# Genetic heterogeneity of the Spy1336/R28 – Spy1337 Virulence Axis in *Streptococcus pyogenes* and Effect on Gene Transcript Levels and Pathogenesis

**DOI:** 10.1101/777987

**Authors:** Jesus M. Eraso, Priyanka Kachroo, Randall J. Olsen, Stephen B. Beres, Luchang Zhu, Traci Badu, Sydney Shannon, Concepcion C. Cantu, Matthew Ojeda Saavedra, Samantha L. Kubiak, Adeline R. Porter, Frank R. DeLeo, James M. Musser

## Abstract

*Streptococcus pyogenes* is a strict human pathogen responsible for more than 700 million infections annually worldwide. Strains of serotype M28 *S. pyogenes* are typically among the five more abundant types causing invasive infections and pharyngitis in adults and children. Type M28 strains also have an unusual propensity to cause puerperal sepsis and neonatal disease. We recently discovered that a one-nucleotide indel in an intergenic homopolymeric tract located between genes *Spy1336/R28* and *Spy1337* altered virulence in a mouse model of infection. In the present study, we analyzed size variation in this homopolymeric tract and determined the extent of heterogeneity in the number of tandemly-repeated 79-amino acid domains in the coding region of *Spy1336/R28* in large samples of strains recovered from humans with invasive infections. Both repeat sequence elements are highly polymorphic in natural populations of M28 strains. Variation in the homopolymeric tract results in (i) changes in transcript levels of *Spy1336/R28* and *Spy1337 in vitro,* (ii) differences in virulence in a mouse model of necrotizing myositis, and (iii) global transcriptome changes as shown by RNAseq analysis of isogenic mutant strains. Variation in the number of tandem repeats in the coding sequence of *Spy1336/R28* is responsible for size variation of R28 protein in natural populations. Isogenic mutant strains in which genes encoding R28 or transcriptional regulator Spy1337 are inactivated are significantly less virulent in a nonhuman primate model of necrotizing myositis. Our findings provide impetus for additional studies addressing the role of R28 and Spy1337 variation in pathogen-host interactions.

## INTRODUCTION

*Streptococcus pyogenes* (group A streptococcus, GAS) is a strict human pathogen responsible for >700 million infections and ∼517,000 deaths annually worldwide (1). Human infections range in severity from relatively mild conditions such as pharyngitis to life-threatening septicemia and necrotizing fasciitis/myositis (2). GAS also causes skin infections such as impetigo and erysipelas (3) and post-infection sequelae, including rheumatic fever (4), rheumatic heart disease (5), and glomerulonephritis (6).

GAS strains are commonly classified based on serologic diversity in M protein, an antiphagocytic cell-surface virulence factor, or allelic variation in the 5’-end of the *emm* gene that encodes this protein (7, 8). More than 250 *emm* types have been identified, but the majority of infections in many countries are caused by a relatively small number of prevalent *emm* types: *emm1*, *emm3*, and *emm12* (9–12). Strains of *emm28* (serotype M28) GAS are of special importance because they are among the top five *emm* types causing invasive infections in the USA (11, 13) and several European countries (14–16). They also have an unusual propensity to cause puerperal sepsis (childbed fever) and neonatal infections (17–21). The molecular mechanisms contributing to the ability of *emm28* strains to cause devastating peripartum infections are poorly understood.

The R28 protein is a surface-associated virulence factor made by *emm28* strains and has been studied as a vaccine candidate (22–24). R28 originally was described by Lancefield and colleagues based on serologic studies (25–27). The gene (*Spy1336/R28*) encoding R protein in *emm28* GAS strains is nearly identical to the *alp3* gene encoding the Alp3 protein in group B streptococcus (GBS), a common cause of neonatal sepsis, pneumonia, and meningitis (24, 28, 29). Genome sequencing of *emm28* GAS strain MGAS6180 discovered that the gene encoding the R28 protein (*Spy1336/R28*) is located on an integrative-conjugative element (ICE)-like element originally designated as region of difference 2 (RD2) (30, 31). In the genome of reference strain MGAS6180 (31), RD2 is a 37.4 Kb segment of DNA that has 34 annotated genes. RD2 is present in a small number of other GAS *emm* types and is >99% identical to a region present in the chromosome of the majority of GBS strains (31). These characteristics suggest that RD2 has been disseminated into different streptococcal strains and species by horizontal gene transfer and recombination (32, 33); published data support this idea (33). The presence of RD2 in GBS and *emm28* GAS strains causing infections associated with the female genital tract also suggests that genes located on this ICE element are causally involved in this clinical phenotype.

The *Spy1337* gene is adjacent to, and divergently transcribed from, *Spy1336/R28* in GAS (Fig. 1A). The Spy1337 protein is a member of the AraC family of prokaryotic transcriptional regulators. Members of this family of regulators commonly have two domains, a conserved C-terminal domain that defines the members of the family, and a variable N-terminal domain that differs among family members. The C-terminal domain comprises approximately 100 amino acids and has two helix-turn-helix (HTH) DNA-binding motifs. The N-terminal domain can be bifunctional, mediating effector-binding and multimerization of the regulator (34, 35). Genes encoding Spy1337/AraC-like proteins are frequently found adjacent to genes encoding surface antigens containing the YSIRK signal sequence motif {(YF)SIRKxxxGxxS}, which is responsible for localized secretion at the division septum (36, 37). This motif is present in the N-terminal domain of R28 at amino acid positions 19 through 30.

**FIG 1.**
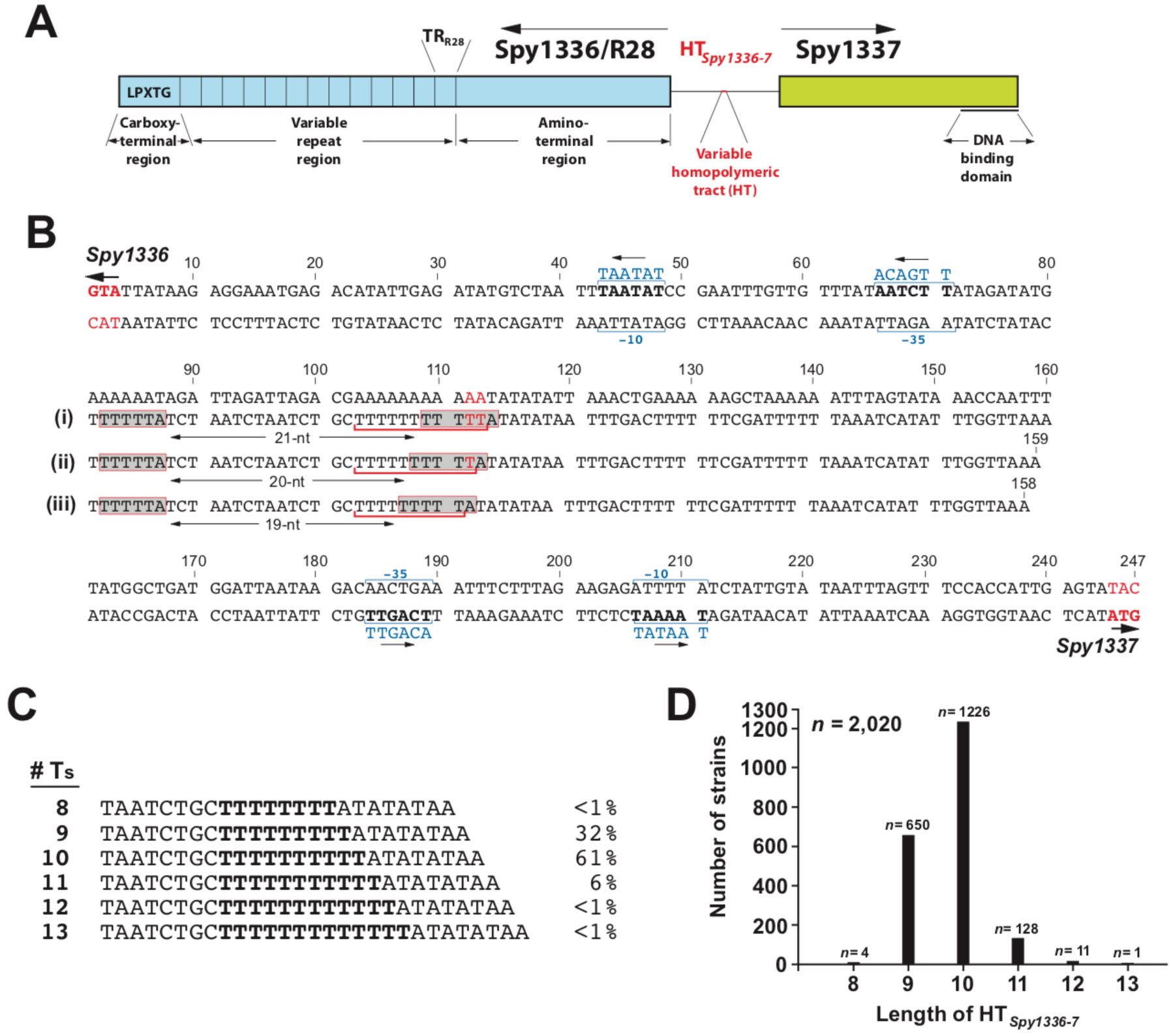
Heterogeneity in the homopolymeric tract located between *Spy1336/R28* and *Spy1337*. (A) Schematic of Spy1336/R28 (R28) and Spy1337. HT*_Spy1336-7_* refers to the variable homopolymeric tract. TR_R28_ refers to one 79-amino acid tandem repeat. LPXTG is the consensus sequence for peptidoglycan attachment. DNA binding domain refers to two predicted helix-turn-helix DNA-binding motifs. Schematic is not drawn to scale. (B) *Spy1336/R28-Spy1337* regulatory region from the reference strain MGAS6180 (31). (i), (ii), and (iii) refer to HT*_Spy1336-7_* containing 11, 10, or 9 T nucleotides, respectively, and the change in the spacing between two ATTTTT (shaded) direct repeats. The HT*_Spy1336-7_* variants are underlined. Inferred promoters are shown. (C) DNA sequence of the prevalent HT*_Spy1336-7_* alleles differing by one T nucleotide. Numbers at right refer to percent of all isolates analyzed with the designated number of Ts. (D) Frequency distribution of the most common HT*_Spy1336-7_* alleles in the *emm28* population. HT*_Spy1336-7_* containing between 8 and 13 T nucleotides are shown with the corresponding number of strains (*n*) in which they are present. The total number of strains represented is 2,020.

Recently, we proposed that the Spy1337 protein is a positive transcriptional regulator of both *Spy1336/R28* and *Spy1337*, and that by regulating expression of *Spy1336/R28* and other genes, Spy1337 is involved in *emm28* GAS virulence (38). To positively regulate virulence gene expression, AraC-like transcriptional regulators usually either bind to chemical effectors present at the site of infection to cause a conformational change favoring DNA-binding to their cognate gene targets (39, 40), or they bind to AraC negative regulators (ANRs) that inhibits such interactions (41). In addition, they can regulate their own expression (42–46).

The R28 protein is a member of a family of streptococcal surface proteins termed <u>a</u>lpha-like <u>p</u>roteins (Alps), characterized by the presence of (i) a conserved amino-terminal domain (pfam08829), (ii) long tandem <u>r</u>epeat (TR) elements (pfam08428), and (iii) a carboxy-terminal domain (pfam00746) containing the LPXTG consensus sequence that serves to attach these proteins to the cell wall (24, 28, 32, 47, 48) (Fig. 1A). Alps participate in adhesion (49–51), can be immunogenic (52, 53), and have been studied in the context of vaccine development (23, 54). The best-studied Alp family proteins are alpha C protein (*bca*) (55), Alp1 (*alp1*) (56), Alp2 (*alp2*), Alp3 (*alp3*) (28), Alp4 (*alp4*) (57), and Rib (*rib*) (58), all made by GBS. Compared to the wild-type parental strain, an isogenic alpha C gene deletion mutant of GBS is less virulent in a mouse model of sepsis (59), demonstrating the role of Alp proteins in virulence. It has been shown that changing the number of tandemly repeated domains in the alpha C protein in GBS altered antigenicity, host immune recognition, and virulence when immunized with alpha C protein.

The molecular mechanism(s) underlying how R28 or Alp proteins contribute to virulence is poorly understood. R28 adheres to human epithelial cells *in vitro* (24) via binding of the N-terminal domain to human integrin receptors α_3_β_1_, α_6_β_1_, and α_6_β_4_ (60), which in turn bind to laminin, an extracellular matrix (ECM) protein (61).

In a recent study of 492 M28 invasive isolates of GAS for which whole genome sequence and transcriptome data were available, we used machine learning to determine that a variable-length T-nucleotide homopolymeric tract (HT, referred to herein as HT*_Spy1336-7_*) in the intergenic region of *Spy1336/R28* and *Spy1337* was associated with differences in transcript levels of these two genes. HTs are commonly characterized by rapid length variation (62). When present in promoter regions, HTs can modify transcript levels by altering the distance between promoter elements (63) or changing the binding of transcription factors (64, 65). In ∼94% of the strains we studied, HT*_Spy1336-7_* had either nine or ten T residues. Comparison of human infection isolates found that strains with 9Ts had significantly lower transcript levels of the *Spy1336/R28* and *Spy1337* genes compared to strains with 10Ts. Compared to a parental strain with 9Ts, an isogenic mutant strain with 10Ts in the HT*_Spy1336-7_* had significantly increased transcript levels of *Spy1336/R28* and *Spy1337* and was significantly more virulent in a mouse necrotizing myositis infection model. In addition, compared to the isogenic strain with 9Ts, the strain with 10Ts was significantly more resistant to killing by human polymorphonuclear leukocytes *ex vivo* and produced more R28 protein. Thus, a one-nucleotide indel in the HT*_Spy1336-7_* caused altered level of these two transcripts and changed the virulence phenotype (38). Of note, there is an HT comprised of 8-15 T residues located in the regulatory region 99 nucleotides (nts) upstream of the *bca* gene encoding the GBS alpha C Alp protein (64).

The *Spy1336/R28* gene has a centrally-located long TR motif (referred to herein as TR_R28_), characteristic of genes encoding Alp family proteins. Due to the homologous nature of repetitive DNA sequence, regions having TRs frequently vary in size as a consequence of mutational events involving either unequal crossover or intramolecular recombination (66–69). The R28 protein made by GAS *emm28* reference strain MGAS6180 (31) has 13 identical TR_R28s_ of 79 amino acids (aa) each (Fig. 1A).

In the present study, we analyzed size variation of the HT*_Spy1336-7_* region located upstream of *Spy1336/R28* in >2,000 strains of *emm28* GAS cultured from invasive human infections and compared transcriptome changes in isogenic strains containing variable lengths of HT*_Spy1336-7_*. We also determined the extent of heterogeneity in the number of TR_R28s_ in *Spy1336/R28* in a subset of 493 M28 invasive clinical isolates, including the reference strain MGAS6180, for which we had whole genome and transcriptome data (38). Finally, we used isogenic mutant strains to determine the contributions of *Spy1336/R28* and *Spy1337* to global gene expression and virulence.

## RESULTS

### Heterogeneity in a homopolymeric tract in the intergenic region between *Spy1336/R28 and Spy1337*

We previously discovered that a single nucleotide indel located in an HT in the intergenic region between the divergently transcribed *Spy1336/R28* and *Spy1337* genes (Fig. 1A and B) significantly altered the transcript levels of the two genes (38). Specifically, strains with 9Ts in the HT produced little or no detectable transcript of these two genes, whereas organisms with 10Ts in this tract produced abundant and significantly increased levels of transcripts (38). Moreover, increased transcript levels of *Spy1336/R28* and *Spy1337* resulted in increased production of the R28 virulence factor and increased virulence in a mouse necrotizing myositis infection model (38). We reported that approximately two-thirds of 493 strains had the 10T variant of the HT*_Spy1336-7_* region, whereas one-third of strains had a 9T variant (38).

Taken together, these observations provided the impetus to expand our study of heterogeneity in the HT*_Spy1336-7_* region to include the entire previously described cohort of 2,095 *emm28* clinical isolates recovered from invasive human infections collected in six countries over a 26-year period (38). The HT*_Spy1336-7_* region was analyzed in three ways, as described in detail in Materials and Methods. First, the contigs from genome assemblies generated with SPAdes (70) for all strains were searched with sequences of 20-nt flanking HT*_Spy1336-7_* on each side using nucleotide Basic Local Alignment Search Tool (BLASTn). The identified HT*_Spy1336-7_* target regions were retrieved, binned by alleles, and alleles were enumerated. Subsequently, the number of T nucleotides in the HT*_Spy1336-7_* for each allele was counted. This method yielded results for the great majority of strains (∼89%). As a second assembly method, we interrogated the Illumina sequencing reads for each strain using a set of eight probes of 31-nts in length corresponding to HT*_Spy1336-7_* alleles with 6 to 13 Ts. In the aggregate, six alleles of HT*_Spy1336-7_* were identified that differed from one another only by the number of T residues, varying in length from 8 to 13Ts (Fig. 1C; Tables S1 and S2). This analysis identified the same approximate frequency distribution for strains containing 9T or 10T nucleotides as described in our previous study (38), namely ∼1/3 (*n* = 650) and ∼2/3 (*n*=1,226), respectively. We also discovered that ∼6% (*n* = 128) of the strains had 11Ts in the HT*_Spy1336-7_* sequence (Fig. 1D). A small number of strains had 8Ts, 12Ts, or 13Ts in the HT*_Spy1336-7_* sequence (Tables S1 and S2). For any strain with discrepant or no results, we visually inspected read alignment {i.e. Binary Alignment Map (BAM)} files corresponding to the *Spy1336*/*R28-Spy1337* intergenic region using TABLET (71).

### Heterogeneity in the number of ALP-family long tandem repeats in the Spy1336/R28 gene and protein

The R28 protein has 3 domains: the amino-terminal and carboxy-terminal domains flank the size-variable central TR_R28_ domain (Fig. 2A). The 424-aa amino-terminal domain has a secretion signal sequence with the YSIRK motif, and the 46-aa carboxy-terminal domain has a cell-wall anchoring sequence with the LPXTG motif characteristic of surface-attached proteins (Fig. 2A). Inasmuch as the number of TRs can vary by recombination or other mechanisms (66–69) we hypothesized that the large number of *emm28* GAS strains we studied could have different size variants of the *Spy1336/R28* gene as a consequence of having different numbers of TR_R28s_. Our previously reported Illumina paired-end 150nt read length whole genome sequence data (38) could not be used to accurately assemble and determine the number of repeats in the TR_R28_ region, as the sequence reads are of insufficient length to span the 237nt repeated motif. Thus, to determine the extent of size variation in TR_R28s_ in the *Spy1336/R28* gene, we used PCR analysis, as described in the Materials and Methods, and studied the 493 strains analyzed previously by RNAseq (38). Overall, we identified TR_R28s_ size variants ranging from 1 to 17 copies of the repeat (Fig. 2B and Table S3). The most common numbers of TR_R28s_ identified was ten (*n*=71, 14.4%) then nine (*n*=69, 14.0%). The inferred molecular weight of several R28 variants was confirmed by Western immunoblot analysis (Fig. 2C).

**FIG 2.**
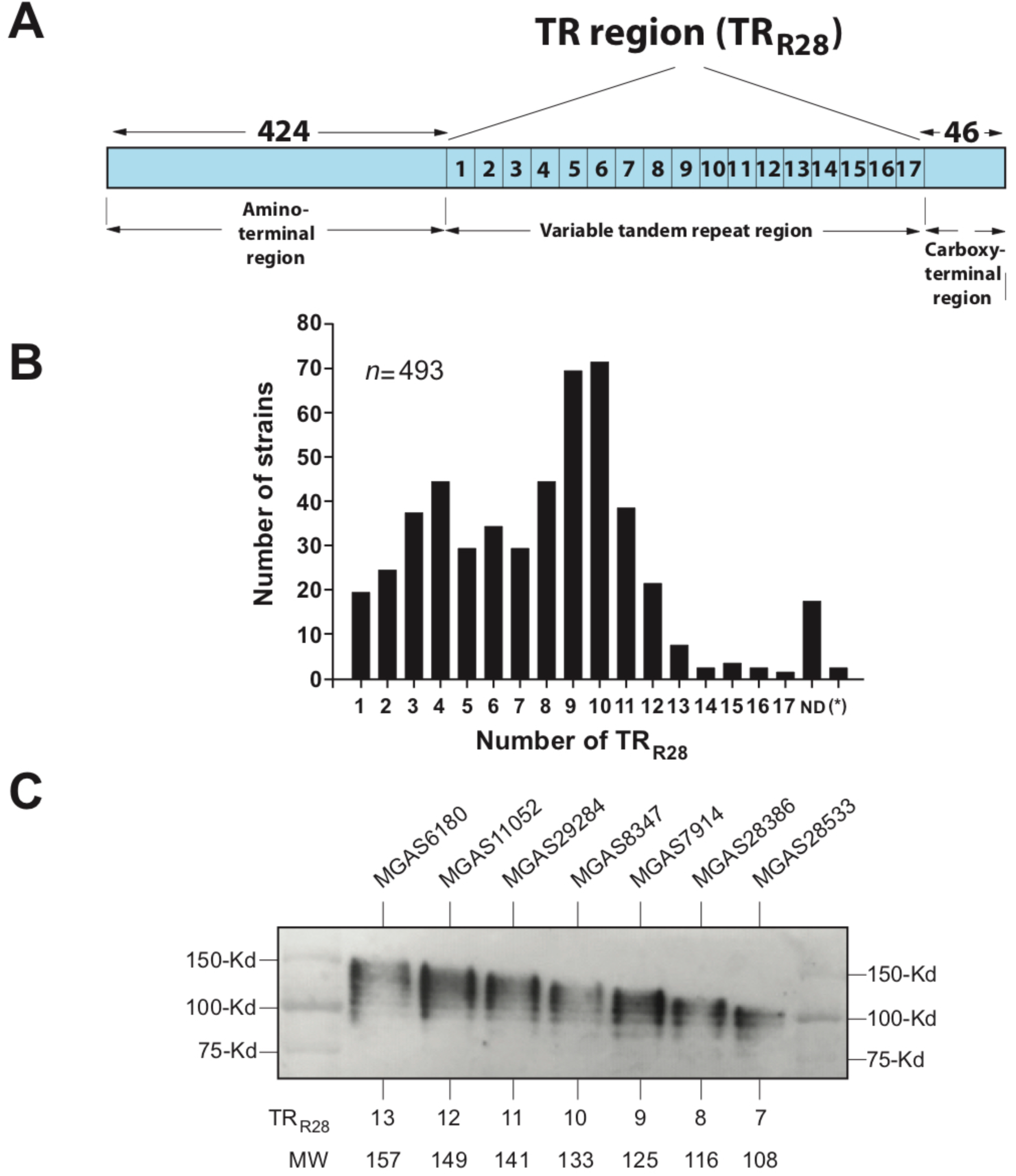
Analysis of the number of tandem repeats in the R28 protein. (A) Schematic of R28. Numbers above the diagram indicate the number of amino acids. TR, tandem repeat. The maximum number of TR_R28s_ found in this study was 17. Schematic is not drawn to scale. (B) Data shown correspond to 493 strains. ND, not determined. (*), two strains studied did not contain the RD2 mobile genetic element, as confirmed by inspection of bam files using TABLET (71). (C) Western immunoblot of strains with R28 proteins containing different numbers of TR_R28_, inferred based on gene sequence data. All strains analyzed had an HT*_Spy1336-7_* with 10Ts because 9T strains do not produce detectable R28 protein. TR_R28_, number of tandem repeats per strain. MW, inferred molecular weight of the R28 protein.

### Construction and growth characteristics of isogenic mutant strains

We previously compared the global transcriptomes of isogenic mutant strains (MGAS27961-9T and MGAS27961-10T) that differ only in the number of T residues in the HT*_Spy1336-7_* region (38). In view of our finding that 6% of strains have 11Ts in this region, we constructed isogenic mutant strain MGAS27961-11T. We also constructed isogenic mutant strains MGAS27961-Δ*Spy1336/R28*, MGAS27961-Δ*Spy1337*, and MGAS27961-Δ*Spy1336/*Δ*Spy1337* in which the target genes were deleted in a parental strain with 10Ts in the HT*_Spy1336-7_* region. The goal of generating these strains was to perform comparative transcriptome and virulence analyses. All strains had a very similar growth curve under the laboratory conditions tested (Fig. S1).

### Transcriptome analysis of clinical strains

We examined the transcriptome data for 442 M28 clinical isolates (38) to determine if variation in the length of HT*_Spy1336-7_* altered the level of transcripts for *Spy1336/R28* and *Spy1337* (Fig. 1C). Among these 442 clinical isolates, 423 strains had alleles exclusively containing indels in HT*_Spy1336-7._* All HT variants were represented in these 423 strains. We restricted examination of the transcriptome data to strains with 8 to 11Ts in the HT*_Spy1336-7_* region because the 12T and 13T variants were represented by only one strain each. Strains with 8 and 9Ts in the HT*_Spy1336-7_* region had low transcript levels of *Spy1336/R28* and *Spy1337* genes (Fig. 3). In contrast, strains with the 10T variant had significantly higher transcript levels of Spy*1336/R28* and *Spy1337* compared to strains with the 9T variants (*P*<0.0001). Strains with the 11T variant (*n*=25) had transcript levels for *Spy1336/R28 (P<*0.0001) and *Spy1337 (P<*0.0024) that are significantly higher than either 10T or 9T strains (Fig. 3).

**FIG 3.**
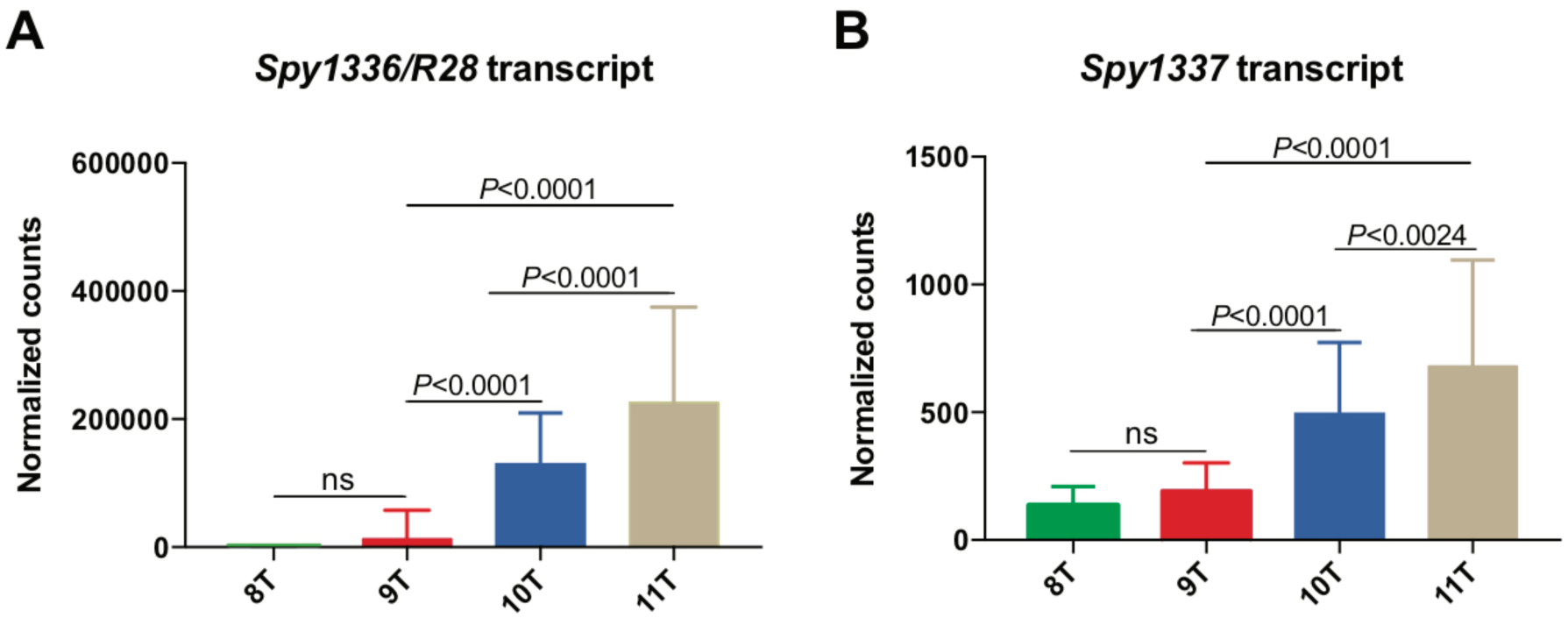
Transcript levels of *Spy1336/R28* and *Spy1337* in 421 M28 clinical isolates. RNAseq was performed on the isolates grown to early stationary phase, as described previously (38). Transcript levels (normalized counts) of (A) *Spy1336/R28* and (B) *Spy1337* for strains with 8Ts (*n*=4), 9Ts (*n*=137), 10Ts (*n*=255) or 11Ts(*n*=25) in the HT*_Spy1336-7_* region. Mean expression levels are plotted and error bars represent standard deviation. *P* values were calculated using the unpaired Student’s t-test. ns, not significant.

### Transcriptome analysis of isogenic mutant strains

We next used RNAseq to test the hypothesis that, when compared to a parental strain with 9Ts (MGAS27961-9T) in the HT*_Spy1336-7_* region, an isogenic mutant strain with 11Ts (MGAS27961-11T) had an altered transcriptome. RNAseq analysis confirmed that the gene expression profile of MGAS27961-11T is substantially altered compared to MGAS27961-9T. Principal component analysis supported these findings (Fig. 4A and B). Moreover, compared to strain MGAS27961-9T, isogenic strain MGAS27961-11T had differential expression of 4.7% (ME) and 6% (ES) of the GAS genome (Fig. S2A). We note that at both phases of growth, transcript levels of *Spy1336/R28* in MGAS27961-11T were significantly higher compared to the MGAS27961-9T strain values (Fig. 4C and D). As expected, no detectable transcript expression of *Spy1336/R28* and *Spy1337* was observed in the isogenic deletion mutant strains, MGAS27961-Δ*Spy1337* and MGAS27961-Δ*Spy1336/*Δ*Spy1337* (Fig. 4, C-F). Virulence genes significantly upregulated included: *nga,* encoding an NADase cytotoxin; *slo*, encoding the cytolytic protein streptolysin O; the *sag* operon encoding streptolysin S; *mga*, a positive transcriptional regulator of multiple virulence genes; *emm28* encoding M protein; and *sof,* encoding serum opacity factor (Fig. S2 and Table S4).

**FIG 4.**
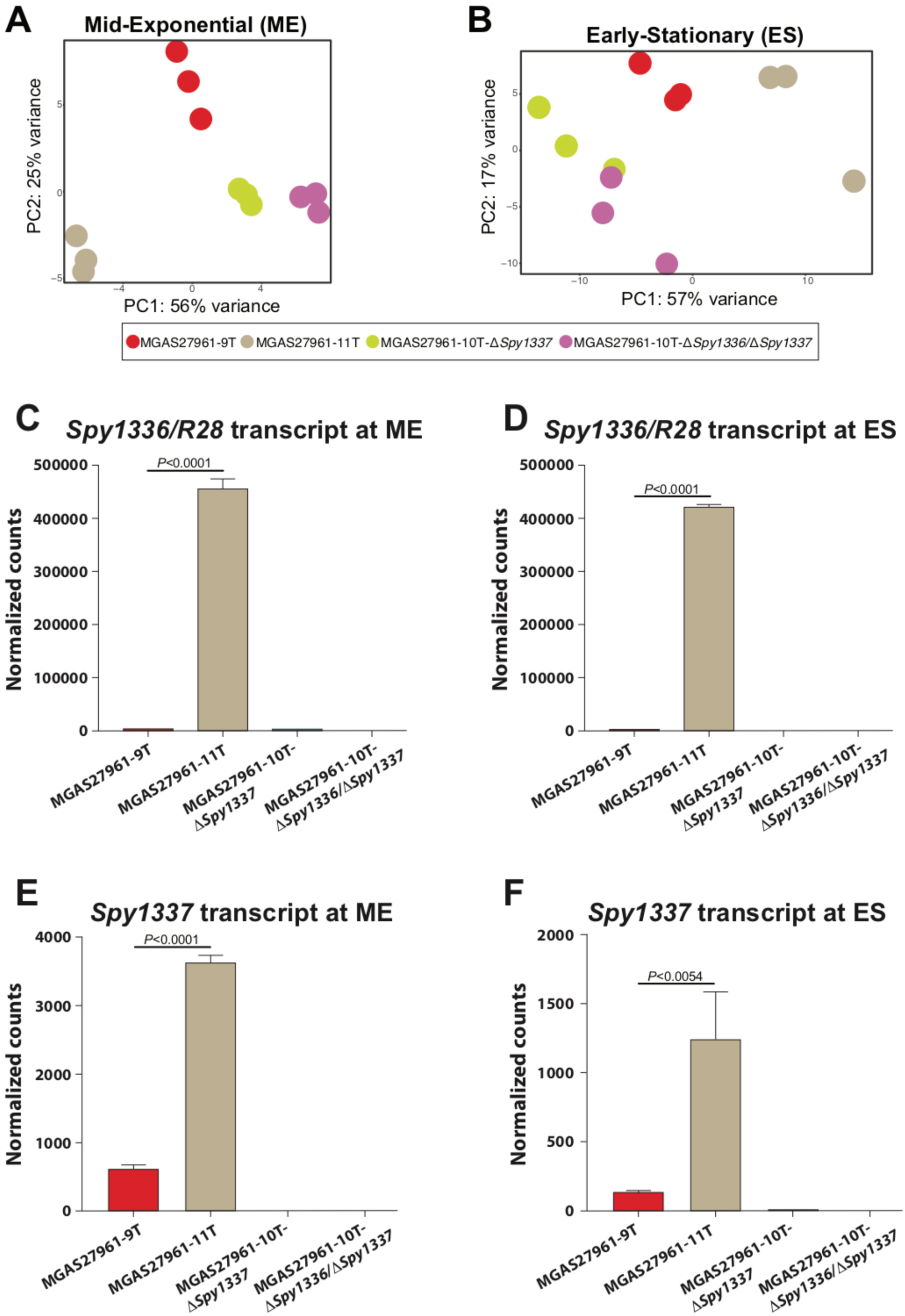
Transcriptome analysis of parental and isogenic mutant strains. RNAseq was performed on isogenic mutant strains MGAS27961-9T, MGAS27961-11T, MGAS27961-Δ*Spy1337*, and MGAS27961-10T-Δ*Spy1336/*Δ*Spy1337* using three biological replicates per strain at two phases of growth: mid-exponential (ME) and early-stationary (ES). Principal component analysis shows distinct clustering of isogenic mutant strains at the (A) ME and (B) ES phases of growth. Strain MGAS27961-11T clusters away from the isogenic strains with mutations that decrease (MGAS27961-9T) or eliminate expression of *Spy1336/R28* and/or *Spy1337* (MGAS27961-Δ*Spy1337* and MGAS27961-10T-Δ*Spy1336/*Δ*Spy1337)*. (C and D) At both ME and ES, transcript levels of *Spy1336/R28* for strain MGAS27961-11T were significantly higher compared to strain MGAS27961-9T. At both phases of growth, no detectable *Spy1336/R28* transcript was observed in the isogenic knockout strains (MGAS27961-Δ*Spy1337* and MGAS27961-10T-Δ*Spy1336/*Δ*Spy1337*. (E and F) Similarly, at both ME and ES, transcript levels of the gene *Spy1337* were significantly higher in MGAS27961-11T compared to strain MGAS27961-9T. Mean transcript levels are plotted and error bars represent standard deviation. *P* values were calculated using the unpaired student’s t-test.

### Heterogeneity in the HT*_Spy1336-7_* region significantly affects virulence in a mouse model of necrotizing myositis

To test the hypothesis that the number of T nucleotides in HT*_Spy1336-7_* region contributes to GAS virulence, we inoculated mice intramuscularly with either the parental strain with 9Ts in HT*_Spy1336-7_* or an isogenic mutant strain with 10Ts or 11Ts. Compared to the parental 9T strain, the isogenic 10T and 11T strains each caused significantly greater mortality and larger lesions with more tissue destruction (Fig. 5). Taken together, the data support the hypothesis that the number of T nucleotides in HT*_Spy1336-7_* significantly affects virulence in this infection model.

**FIG 5.**
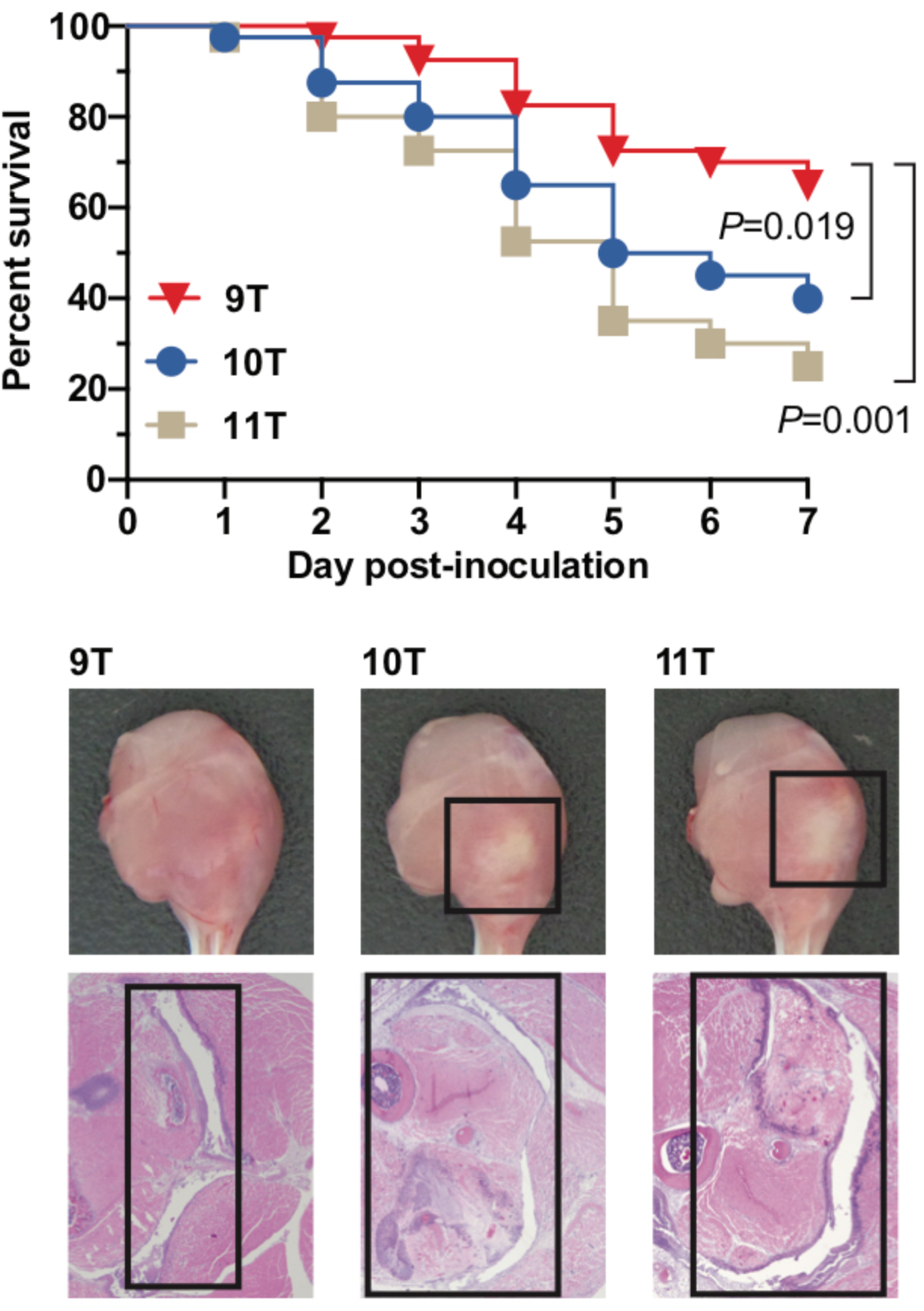
Virulence assessed in a mouse model of necrotizing myositis. (A) Virulence of 9T, 10T, and 11T isogenic strains in a mouse model of necrotizing myositis (*n* = 40 mice per strain). Significantly increased near-mortality was observed for the 10T and 11T strains compared with the 9T strain. Significance was determined using the log-rank test. (B) Representative gross and microscopic pathology images of the hindlimb lesions from mice (*n* = 5 mice per strain). Boxed areas demarcate areas with major lesions.

### Analysis of the R28 protein made by the isogenic mutant strains

In most bacteria, gene transcript levels typically correlate with amounts of the encoded proteins made, but this is not always the case (72, 73). We reported previously that the R28 protein is produced and detected by Western immunoblot in whole cell extracts and supernatants derived from MGAS27961-10T, but not from the isogenic strain MGAS27961-9T. This finding is consistent with RNAseq data showing that the transcript level of *Spy1336/R28* was higher in MGAS27961-10T compared to MGAS27961-9T (38).

To build on our previous observations, we used Western immunoblot analysis to assess R28 production by a strain with an HT*_Spy1336-7_* 11T variant and isogenic strains MGAS27961-11T, MGAS27961-10T-Δ*Spy1336*, and MGAS27961-10T-Δ*Spy1337* (Fig. 6A). As anticipated, R28 protein was undetectable in strain MGAS27961-10T-Δ*Spy1336/R28,* and production by strain MGAS27961-10T-Δ*Spy1337* was also not detected under the conditions tested (Fig. 6A). The lack of discernible R28 production by the MGAS27961-10T-Δ*Spy133*7 strain is consistent with our hypothesis that Spy1337 is a positive transcriptional regulator of *Spy1336/R28* (38).

**FIG 6.**
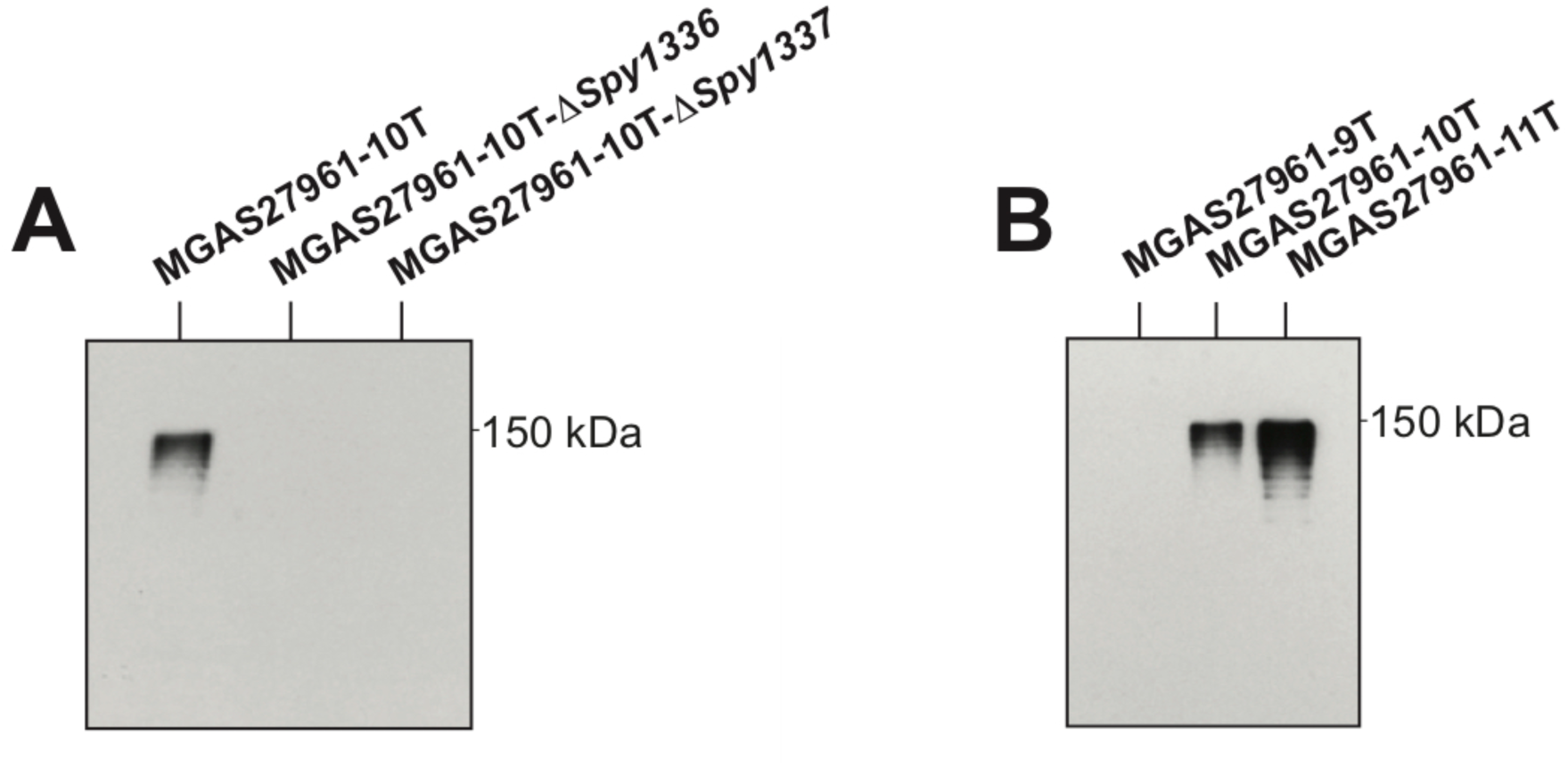
Western immunoblot analysis of Spy1336/R28. Isogenic strains were collected at mid-exponential phase (OD_600_ = ∼0.6). Whole cells were assayed for the presence of immunoreactive R28 protein with an anti-R28 polyclonal antibody (38). (A) Isogenic deletion mutants assayed with anti-R28. (B) Isogenic HT*_Spy1336-7_* size variant mutants assayed with anti-R28. 10% polyacrylamide gels were used.

Next, we analyzed R28 protein production by parental and isogenic mutant strains containing the three different length variants of HT*_Spy1336-7_* (Fig. 6B). Consistent with previous data, we did not detect production of R28 by parental strain MGAS27961-9T, whereas immunoreactive R28 was produced by isogenic mutant strain MGAS27961-10T (38). Also consistent with our hypothesis and RNAseq data, isogenic mutant strain MGAS27961-11T produced greater amounts of immunoreactive R28 compared to strain MGAS27961-10T and parental strain MGAS27961-9T (Fig. 6B).

### *Spy1336/R28* and *Spy1337* contribute significantly to virulence in a non-human primate (NHP) model of necrotizing myositis

To test the hypothesis that *Spy1336/R28* and *Spy1337* contribute to GAS virulence, we inoculated NHPs with parental strain MGAS27961-10T and isogenic mutant strains MGAS27961-10T-Δ*Spy1336/R28* or MGAS27961-10T-Δ*Spy1337*. Compared to the parental strain, each of the two isogenic deletion-mutant strains caused significantly smaller lesions characterized by less tissue destruction (Fig. 7A and C). In addition, compared to the parental strain, significantly fewer CFUs of each isogenic mutant strain were recovered from the inoculation site (Fig. 7B). Taken together, the data support the hypothesis that *Spy1336/R28* and *Spy1337* contribute to virulence in this infection model.

**FIG 7.**
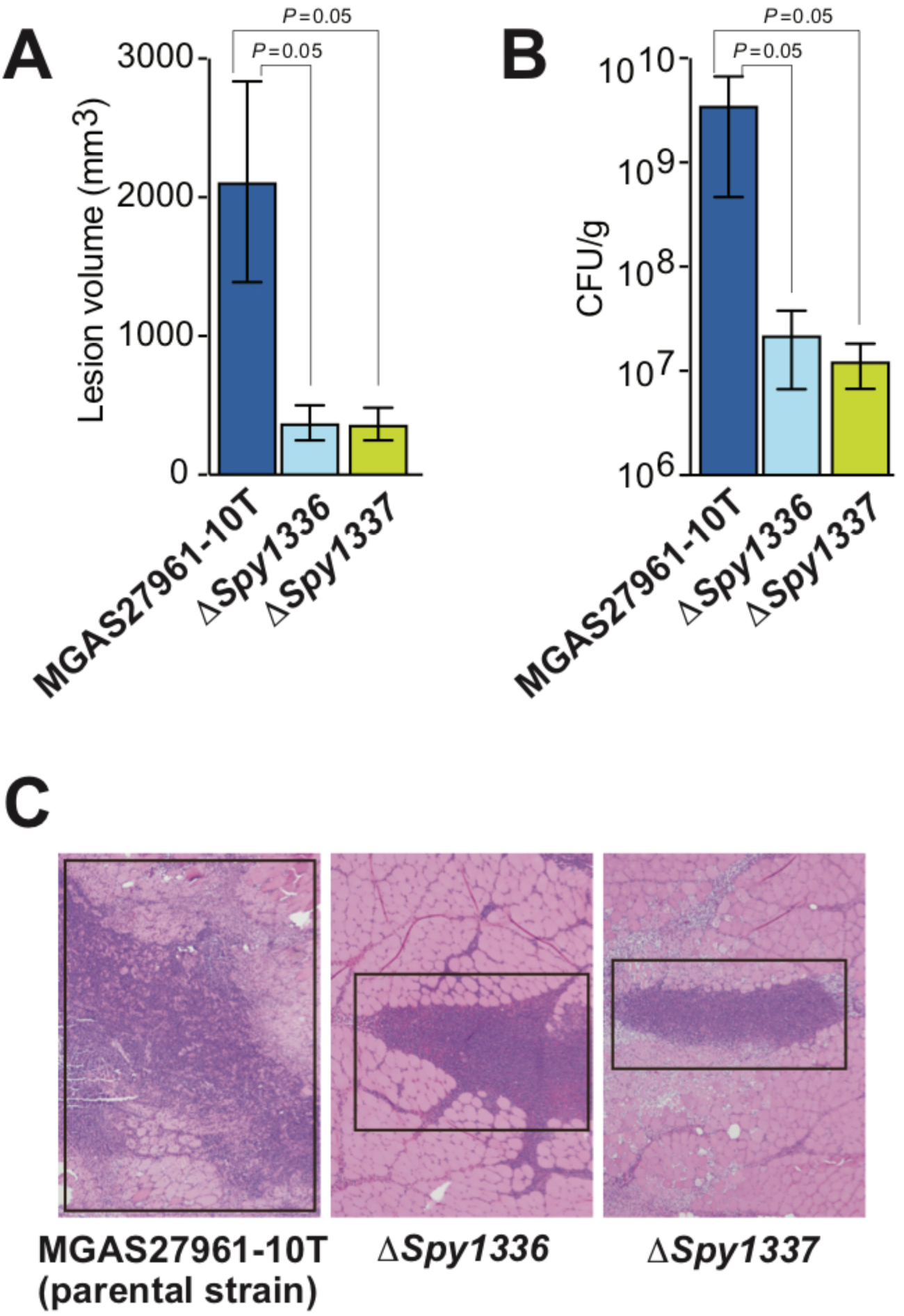
Virulence assessed in a nonhuman primate model of necrotizing myositis. Compared to the parental wild-type strain, deletion of *Spy1336/R28* (Δ*Spy1336*) or *Spy1337* (Δ*Spy1337*) significantly decreases (A) lesion volume, (B) CFU recovery, and (C) tissue destruction. Shown in panel C are representative microscopic images of the necrotic lesion centered at the GAS injection site (black box); hematoxylin and eosin stain, original magnification 4X. *P* = 0.05, Mann-Whitney Test. *n* = 3 NHPs/GAS strain.

## DISCUSSION

*S. pyogenes*, a leading cause of human morbidity, mortality, and healthcare costs globally, produces a large number of extracellular virulence factors. Strains of serotype M28 *S. pyogenes* are repeatedly associated with puerperal sepsis (childbed fever) and are a prominent cause of pharyngitis in many countries. The molecular mechanisms responsible for puerperal sepsis, other invasive infections, and pharyngitis caused by serotype M28 strains are poorly understood. Several lines of evidence indicate that the *Spy1336/R28* gene encodes a cell surface-anchored virulence factor (24, 74) that is involved in *S. pyogenes* pathogenesis. We analyzed the *Spy1336/R28* gene encoding the R28 protein in approximately 2,000 *emm28* invasive strains. We found DNA sequences located in the upstream regulatory region and repeat sequences in its coding sequence that make *Spy1336/R28* highly polymorphic when compared across our population sample; importantly, these polymorphisms affect virulence.

We previously reported a bimodal distribution in the transcripts made by *Spy1336/R28* and *Spy1337* in a study of 492 *emm28* strains (38). Namely, strains with 9Ts in HT*_Spy1336-7_* located between *Spy1336/R28* and *Spy1337* produced low transcript levels (Fig. 1A and B), whereas strains with 10Ts produced significantly greater transcript levels (38). The current work confirmed and extended our observations and found that in the sample of ∼2,000 strains, and when considering alleles exclusively containing indels in the HT*_Spy1336-7_*, (i) the number of T residues varied between 8 and 13, (ii) most strains (∼90.4%) had 9T (32%) or 10T (61%) residues, (iii) an 11T variant was present in 6% of the strains (Fig. 1 and 8), and (iv) non-isogenic clinical isolates and an isogenic strain with an 11T variant had significantly higher transcript levels of *Spy1336/R28* and *Spy1337*, and produced more R28 protein than 9T and 10T variant isogenic strains (Fig. 1, 3, 4, 6, 7, and 8, and Tables S1 and S2).

**FIG 8.**
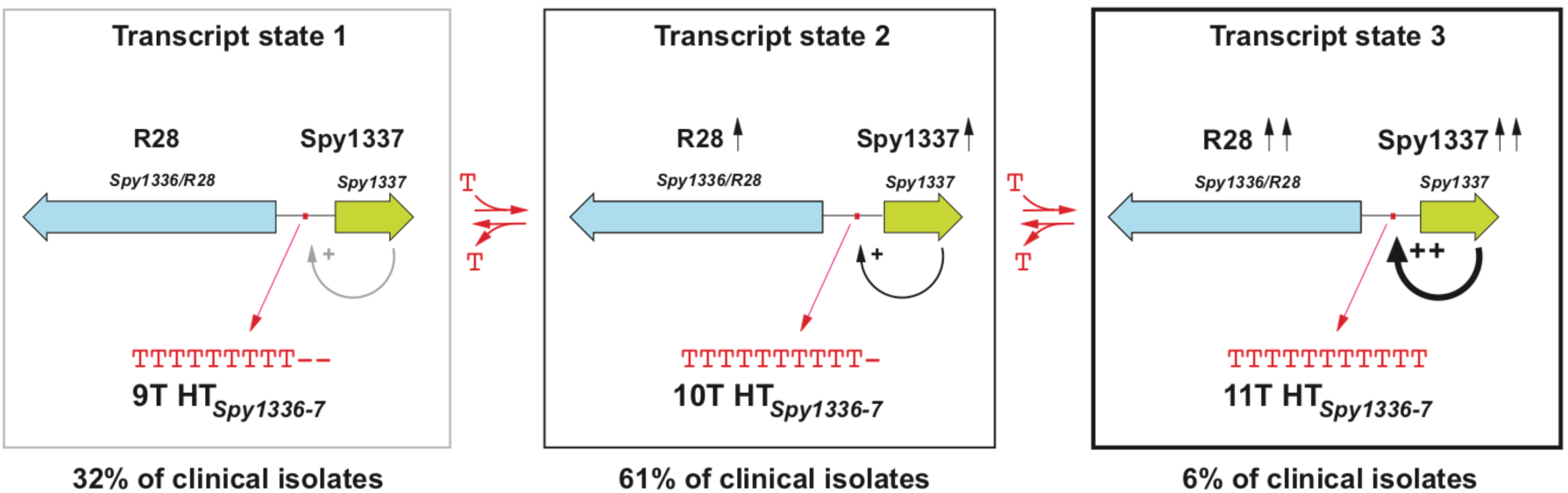
Frequency distribution of HT*_Spy1336-7_* size variants in the *emm28* strains studied and effect on *Spy1336/R28* and *Spy1337* transcript levels. Addition or deletion of one T nucleotide in the homopolymeric tract results in a statistically significant change in transcript levels for *Spy1336/R28* and *Spy1337* and a consequent effect on virulence. There are three transcript states based on number of T residues in the HT*_Spy1336-7_* region. In state 1 (9Ts, left), there are essentially no detectable transcripts of *Spy1336/R28* and *Spy1337*. State 2 (10Ts, center) is characterized by increased transcripts of these two genes, and state 3 (11Ts, right) results in further increase in transcripts of *Spy1336/R28* and *Spy1337*.

Size variants of the HT*_Spy1336-7_* may arise because HTs, especially poly(T)-containing HTs such as HT*_Spy1336-7_*, are able to form transient mispaired regions leading to slipped-strand mispairing during DNA replication, resulting in expansion or contraction of the HT (75–80). HTs located at the 5’ upstream untranslated regions of genes such as the HT*_Spy1336-7_* can contribute to altered regulation of gene transcript expression (64, 81). In this regard, HTs may be involved in phase variation (82) and bacterial adaptation (69, 83). The finding that the vast majority of strains had a 9T or 10T genotype suggests that this system may influence GAS-human interaction in some settings. Under one scenario, lack of or very low transcript production (9T genotype) is advantageous to the organism in certain currently undefined physiologic conditions. Conversely, the significantly-higher transcript level conferred by the 10T genotype could be advantageous in other conditions, also currently not defined. The isogenic mutant strain with the 11T genotype had a higher level of *Spy1336/R28* transcript (Fig. 4C and D), and higher level of production of R28 protein (Fig. 6B), than the strain with the 10T genotype. We hypothesize that the increased R28 protein made by the 11T strain confers only slightly enhanced fitness to the organism (relative to the 10T genotype), an idea that is consistent with the observation that relatively few (6%) natural clinical isolates have this genotype, and substantiated by the fact that the isogenic mutant strain had only modestly increased virulence in the mouse model of necrotizing myositis. Alternatively, too much R28 protein could be detrimental during a natural course of infection in the human host by promoting too tight adherence, leading to decreased dissemination, or even promoting an increased immune response. Consistent with these ideas, the 10T and 11T isogenic mutant strains did not differ significantly in resistance to killing by human PMNs *ex vivo* although compared to the 9T wild-type parental strain, these two mutant strains had significantly enhanced resistance to killing by human PMN (Fig. 9). In terms of the small number of strains containing additional size variants of the HT region (8Ts, 12Ts, and 13Ts), strains with 8Ts produce very minimal levels of *Spy1336/R28* transcripts (Fig. 3A), and those with 12Ts and 13Ts might not further enhance pathogen fitness, or alternatively, global transcript changes occurring in strains with these genotypes in some way decrease fitness. Clearly, additional investigations are required to address these ideas.

**FIG 9.**
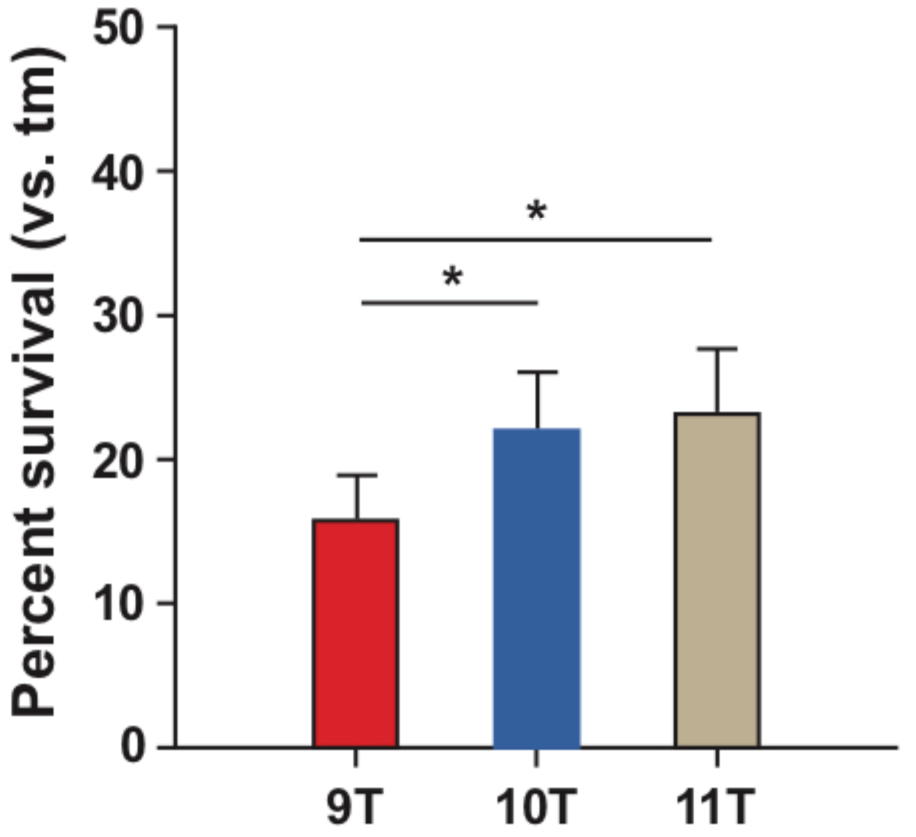
Resistance to killing by human PMNs. Survival of 3 variant HT*_Spy1336-7_* isogenic strains in the presence of purified human PMNs. Bacterial survival data are expressed as the mean ± SEM. \**P* < 0.05 using a repeated-measures one-way ANOVA and Tukey’s posttest (*n* = 7 experiments). tm, time-matched control lacking PMNs.

Puopolo and Madoff reported that deletion or expansion of a 5-nt repeat (AGATT) adjacent to the poly(T) tract is associated with a null phenotype for expression of the *bca* gene encoding the alpha C protein in GBS (64). This region is highly conserved in GAS and GBS. However, only a small number of strains were studied. Here, analysis of 2,095 clinical isolates recovered from human patients with invasive disease identified only five strains with variation in this pentanucleotide motif (Table S2), whereas most of the variation was found in HT*_Spy1336-7_* region. Thus, in the natural populations of GAS we studied, variation in HT*_Spy1336-7_* is the key driver of transcript level changes of *Spy1336/R28* and *Spy1337* (38).

Neither *Spy1336/R28* transcript nor R28 protein production were detectable in an isogenic strain in which the *Spy1337* transcription factor gene was deleted. In addition, deletion of either *Spy1336/R28* or *Spy1337* resulted in a significant decrease in virulence in an NHP model of necrotizing myositis (Fig. 7). Thus, when taken together, our present and previous (38) data are consistent with a model in which the regulation of expression of the gene encoding the R28 virulence factor is partly dependent on a process whereby indels occurring in the HT*_Spy1336-7_* affect binding of the Spy1337 transcriptional regulator to this DNA region by altering its consensus binding site, and/or changing the spacing, and therefore spatial orientation between two adjacent binding sites (Fig. 1B). In this regard, the DNA sequence ATTTT present twice in the *Spy1336/R28-Spy1337* regulatory region resembles part of the consensus binding site for the AraC transcriptional regulator ToxT (84). In our proposed model, Spy1337 positively regulates the expression of both *Spy1336/R28* and *Spy1337* (Fig. 8).

The R28 virulence factor binds through its N-terminal domain to integrin receptors α_3_β_1_, α_6_β_1_, and α_6_β_4_ (60), which in the human host bind to laminin, an ECM protein (61). This constitutes an additional instance in which a pathogen binds to host integrins (85–88), a strategy described for other GAS proteins (89).

For example, GAS PrtF1 binds to α_5_β_1_ integrins and could trigger integrin clustering and internalization of α_5_β_1_ integrin, ultimately resulting in GAS uptake (90). The second region in *Spy1336/R28* containing repeat sequences is located in its coding sequence (Fig. 1A and Fig. 2A). A 237-nt repetitive sequence, corresponding to one 79-aa TR_R28_, is present in *Spy1336/R28* from 1 to 17 tandem copies (Table S3). Ten is the most prevalent number of TR_R28_ copies present in R28, followed by 9 (Fig. 2B). Thus, *emm28* GAS strains make different size variants of the R28 protein (Fig. 2C), likely arising through recombination (66–69, 91). The function of TRs in virulence factors is not well understood. One possible function would be to extend the reach of a surface-anchored protein and expose its N-terminal domain at the bacterial surface without adding new mechanistic functions (92). Variation in the number of TRs may alternatively produce antigenic variation. In this regard, variation in the conserved region of GAS M protein generates antigenic diversity (92), and similarly, variation in the GBS alpha C protein affects antigenicity and protective efficacy (52, 93). TR number variation might decrease adhesion to and entry into host cells (94). No correlation was found between the number of T nucleotides in HT*_Spy1336-7_* and the number of TR_R28_ repeats (Fig. S3). Additional studies designed to address these ideas are warranted.

To summarize, the R28 virulence factor in GAS is highly polymorphic in natural populations, both in level of transcript production and protein expression, and size. Differences in transcription levels are caused by variation in the number of Ts in a homopolymeric tract upstream of the *Spy1336/R28* gene, whereas variance in R28 protein size is caused by variation in the number of identical TR_R28_ tandem repeats in the structural gene. Isogenic mutant strains in which genes encoding R28 or transcriptional regulator Spy1337 are inactivated are significantly less virulent in a NHP model of necrotizing myositis. These findings provide new information about the extent of natural genetic diversity and virulence in this two-gene virulence axis and set the stage for additional studies addressing the role of *Spy1336/R28* and *Spy1337* in pathogen-host interactions.

## MATERIALS AND METHODS

### Bacterial strains and growth conditions

GAS strains were grown at 37°C in Todd-Hewitt broth (Bacto Todd-Hewitt broth; Becton Dickinson and Co.) supplemented with 0.2% yeast extract (THY medium). THY medium was supplemented with chloramphenicol (20 µg ml^-1^) as needed. Trypticase soy agar supplemented with 5% sheep blood (Becton Dickinson and Co.) was used as required. *E. coli* strains were grown in Luria-Bertani (LB) medium at 37°C, unless indicated otherwise. LB medium was supplemented with chloramphenicol (Acros Organics; 20 µg ml^-1^) as needed.

### DNA manipulation

Standard protocols or manufacturer’s instructions were used to isolate plasmid DNA, and conduct restriction endonuclease, DNA ligase, PCR, and other enzymatic treatments of plasmids and DNA fragments. Enzymes were purchased from New England Biolabs, Inc (NEB). Q5 high-fidelity DNA polymerase (NEB) was used. Oligonucleotides were purchased from Sigma Aldrich.

### Analysis of the number of T residues in the HT*_1336-7_*

A blastable database of SPADES (70) assemblies was made for 2,095 *emm28* genomes (38). Using two adjacent DNA sequences of 20-nts that flank HT*_1336-7_*, the DNA region surrounding these two DNA sequences, including the two sequences, was extracted from each strain, and the number of T nucleotides counted. In addition, we analyzed the number of T nucleotides present in HT*_Spy1336-7_* in the 2,095 *emm28* genomes with the command-line program Jellyfish (95), which uses k-mers, and counted the occurrences of each HT*_Spy1336-7_* variant. Using these combined approaches we identified the length of HT*_Spy1336-7_* in 2,074 (∼99%) of the original 2,095 strains, corresponding to 30 different alleles (Table S1 and Table S2). Of these alleles, six contained indels in HT*_Spy1336-7_* exclusively, which resulted in changes in the number of consecutive T nucleotides, with no additional polymorphisms. These six alleles were present in 2,020 (∼97%) strains (Fig. 1C and D).

### Chromosomal DNA extraction and PCR amplification of the *Spy1336/R28* repeat region

Chromosomal DNA extraction was performed as described (96), using Fast-Prep lysing Matrix B beads in 2-ml tubes (M P Biomedicals), or the DNeasy blood and tissue kit (Qiagen). The primer sequences used for PCR-based size determination of the *Spy1336/R28* repeat region are shown in Table S5. Three different primer sets were used. All primers were designed to bind to conserved regions located upstream, in the case of the forward (FWD) primers, or downstream, for the reverse (REV) primers, from the DNA sequence of the *Spy1336/R28* gene encoding the repeat region. Primer set 1 comprised primers JE433 (FWD), binding 60-nt upstream of the repeat region, and JE431 (REV), binding 226-nt downstream of the repeat region. Primer set 2 included primers JE432 (FWD), binding 255-nt upstream of the repeat region, and JE431 (REV). Primer set 3 was comprised of primers JE410 (FWD), binding 2,264-nt upstream of the repeat region, and JE412 (REV), binding 1,098-nt downstream of the repeat region. The extension time used for the PCR reactions was adjusted to accommodate anticipated PCR fragment length. Typically primer set 1 was used first to obtain the desired PCR product and the other 2 primer sets were used if primer set 1 failed to yield an amplified product. Two different DNA ladders were used to determine the size of the PCR products, including the *exACTGene* 100 bp-10,000 bp DNA ladder (Fisher Scientific), and the 100-bp DNA step ladder (Promega). Both ladders were loaded at least 3 times in every agarose gel and used as reference to determine DNA fragment length.

### Western immunoblot analysis

Bacteria grown in THY were collected at OD_600_=∼0.6 (ME), centrifuged at 16,100*g* for 1 min, and pellets were resuspended in PBS. The Western immunoblot procedure used has been described (38), with the following modifications: (i) protein transfer to nitrocellulose membranes was done for 80 (Fig. 6) or 45 min (Fig. 2C) at 120 volts, and (ii) the anti-R28 antibody (38) was diluted to 1:1,250 in PBS-T with 5% nonfat dry milk, whereas the HRP-conjugated anti-rabbit secondary antibody was used at 1:13,500 dilution.

### Construction of isogenic mutant strains

All isogenic mutant strains used in this study are listed in Table S6. Isogenic mutant strain MGAS27961-11T containing an 11T HT*_Spy1336-7_* was generated using allelic exchange as described previously (97). Briefly, primers HPN-1 and HPN-2 (38) were used to amplify a ∼2,690-bp fragment using genomic DNA of MGAS11108, an *emm28* clinical isolate with the naturally occurring 11T nucleotide region. The amplicon encompasses HT*_Spy1336-7_*. The resulting PCR product was cloned into suicide plasmid pBBL740 and transformed into parental strain MGAS27961-9T. The plasmid integrant was used for allelic exchange as described previously (97). To identify strains putatively containing the allelic replacement region encompassing the expected polymorphism (11 T nucleotides), we sequenced the *Spy1336/R28* upstream region using primer HPN-seq (Table S5).

Isogenic mutant strain MGAS27961-10T-Δ*Spy1336* was constructed using MGAS27961-10T genomic DNA as template for amplification. All primers are listed in Table S5. Primer sets 1336-1 and −2 and 1336-3 and −4 were used to amplify two fragments upstream and downstream, respectively, of *Spy1336/R28*. The two PCR fragments were merged by combinatorial PCR and ligated into the BamHI site of suicide vector pBBL740. The recombinant plasmid containing a *Spy1336/R28* deletion encompassing the entire gene was transformed into strain 27961-10T to replace the native *Spy1336/R28* via allelic exchange. Isogenic mutant strains MGAS27961-10T-Δ*Spy1337* and the MGAS27961-10T-Δ*Spy1336*/Δ*Spy1337* double mutant were constructed with methods analogous to those described above. Primer sets 1337-1 and −2 and 1337-3 and −4 were used to generate MGAS27961-10T-Δ*Spy1337*, and 1336-1337-1 and −2 and 1336-1337-3 and −4 were used to generate the double mutant. Whole genome sequence analysis on the isogenic mutant strains confirmed the absence of spurious mutations.

### RNAseq library preparation, sequencing, and analysis

Isogenic *emm28* strains were grown in triplicate in THY and harvested at mid-exponential (ME; OD_600_=0.46-0.52) and early-stationary (ES; OD_600_=1.65-1.7) phases of growth. Bacteria (2 ml) from the ME phase and ES phase (1 ml) were added to 4 ml and 2 ml of RNAprotect Bacteria Reagent (Qiagen), respectively, incubated at room temperature for 20 min, and centrifuged at 4,000 rpm for 15 min. The supernatant was discarded, and the bacterial pellet was frozen in liquid nitrogen and stored at -80°C. The RNeasy kit (Qiagen) was used for total RNA isolation, and the quality of the total RNA was evaluated with RNA Nano chips (Agilent Technologies) and an Agilent 2100 Bioanalyzer. RNA extraction for all *emm28* isogenic strains was performed as described previously (38, 96, 98). The rRNA was depleted with the Ribo-Zero rRNA removal kit for Gram-positive bacteria (Illumina). The quality of the rRNA-depleted RNA was evaluated with RNA Pico chips (Agilent Technologies) and an Agilent 2100 Bioanalyzer. NEBNext Ultra II DNA library prep kit (NEB) was used to prepare the cDNA libraries, according to the manufacturer’s instructions. The quality of the cDNA libraries was evaluated with High-Sensitivity DNA chips (Agilent Technologies) and an Agilent 2100 Bioanalyzer. The cDNA library concentration was measured fluorometrically with Qubit dsDNA BR and HS assay kits (Invitrogen). Analysis of the RNAseq data was performed as described previously (38).

### Necrotizing myositis infection models

A mouse model of necrotizing myositis was used to compare virulence of the 9T, 10T and 11T isogenic strains as previously described (38). Briefly, mice were inoculated in the right hindlimb with 5×10^8^ CFU (*n*=40 mice per strain) and followed for 7 days. A well-described NHP model of necrotizing myositis was used to compare the virulence of the wildtype and isogenic MGAS27961-10T-Δ*Spy1336/R28* and MGAS27961-10T-Δ*Spy1337* strains (96, 99). Five cynomolgus macaques (2-3 years, 2-4 kg) were used. Animals were randomly assigned to different strain treatment groups and inoculated with 5×10^9^ CFU/kg of one strain in the right limb and a different strain in the left limb. Each strain was tested in triplicate. The animals were observed continuously and necropsied at 24 h post-inoculation. Lesions (necrotic tissue) were excised, measured in three dimensions, and volume was calculated using the formula for an ellipsoid. A full-thickness section of tissue taken from the inoculation site was fixed in 10% phosphate buffered formalin and embedded in paraffin using standard automated instruments. Histology of the three sections taken from each limb was scored by a pathologist blinded to the strain treatment groups (99, 100). To obtain the quantitative CFU data, diseased tissue recovered from the inoculation site was weighed, homogenized (Omni International) in 1 mL PBS, and CFUs were determined by plating serial dilutions of the homogenate. Statistical differences between strain groups were determined with the Mann-Whitney test. Animal studies were approved by the Institutional Animal Care and Use Committee at Houston Methodist Research Institute (protocol numbers AUP-1217-0058 and AUP-0318-0016).

### Neutrophil bactericidal activity assays

Neutrophil bactericidal activity assays were performed in accordance with protocol 01-I-N055, approved by the Institutional Review Board for human subjects, National Institute of Allergy and Infectious Diseases. All volunteers gave written informed consent prior to participation in the study. Human neutrophils were isolated from the venous blood of healthy volunteers using a standard method (101). Killing of *S. pyogenes* by human neutrophils was performed as described previously (38), except assay tubes were rotated for 3 h at 37°C.

### Statistical analysis

Unless otherwise stated, error bars represent standard deviation (SD), and *P* values were calculated using either Kruskal-Wallis, or log-rank tests. Differential expression analysis was performed using DESeq2 1.16.1. Genes were considered differentially expressed if the fold-change was greater than 1.5-fold and associated with adjusted *P* value (Bonferroni corrected) < 0.05. For mouse survival studies, results were graphed as Kaplan-Meier curves and data were analyzed using the log-rank test with *P* < 0.05 considered to be significant. For the NHP virulence studies, lesion volume and CFU data were graphed as mean +/-SEM and analyzed using the Kruskal-Wallis test with *P* < 0.05 considered to be significant.

## Data availability

Transcriptome data has been deposited in the Gene Expression Omnibus under accession GSEXXXXXX.

## ACKNOWLEDGMENTS

This study was supported in part by the Fondren Foundation, Houston Methodist Hospital and Research Institute and National Institutes of Health grant AI139369-01A1 (to J.M.M.), and the Intramural Research Program of the National Institute of Allergy and Infectious Diseases, National Institutes of Health (to F.R.D.). We thank Dr. Leslie Jenkins for veterinary assistance and Dr. Kathryn Stockbauer for critical comments to improve the manuscript.

