## Supplemental Figures and Tables for "Genetic heterogeneity of the Spy1336/R28 – Spy1337 Virulence Axis in *Streptococcus pyogenes* and Effect on Gene Transcript Levels and Pathogenesis"

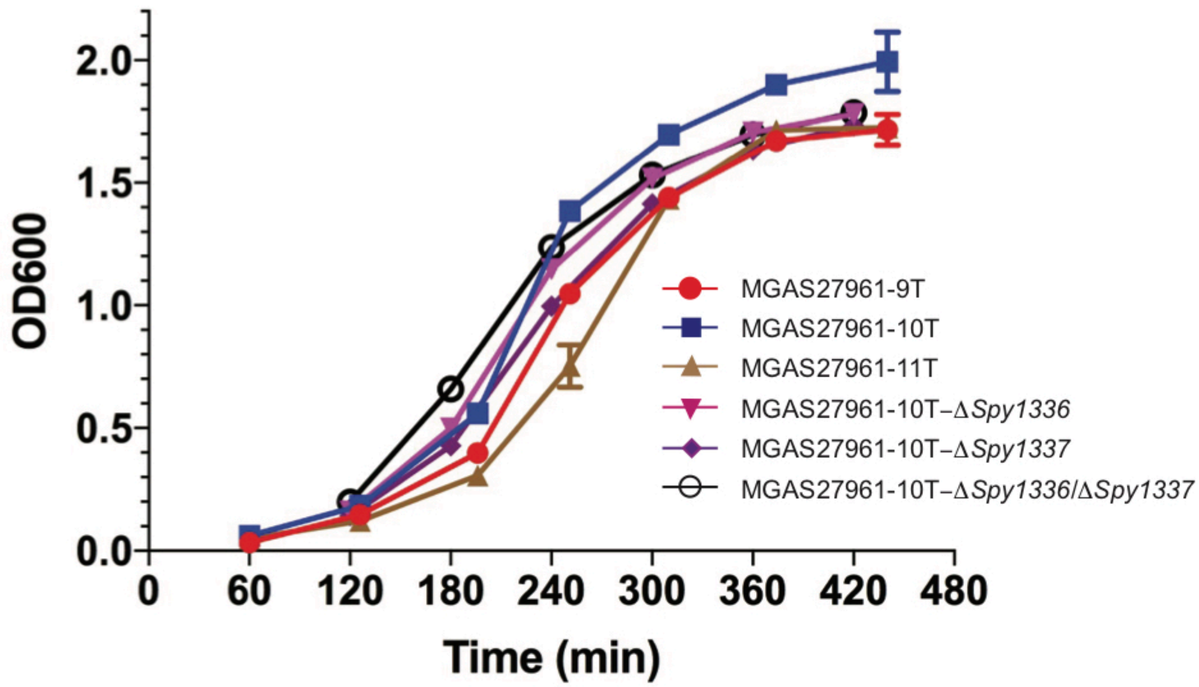


**FIG S1. Growth curves of the parental and engineered isogenic mutant strains in THY**. Strains were grown at 37°C in THY medium.

#### A. Number of differentially expressed genes

| | 10T vs 9T[1] | | 11T vs 9T[2] | | 11T vs 10T[3] | | |
|---|---|---|---|---|---|---|---|
| | Number of genes | %[4] | Number of genes | % | Number of genes | % | Growth phase |
| Upregulated | 18 | 1% | 59 | 3.2% | 8 | 0.4% | ME |
| Downregulated | 13 | 0.7% | 28 | 1.5% | 7 | 0.4% | ME |
| Upregulated | 102 | 5.6% | 12 | 0.7% | 8 | 0.4% | ES |
| Downregulated | 154 | 8.4% | 98 | 5.4% | 5 | 0.3% | ES |

(1) MGAS2761-10T compared to MGAS2761-9T
(2) MGAS2761-11T compared to MGAS2761-9T
(3) MGAS2761-11T compared to MGAS2761-10T
(4) Percentage of genes compared to the entire genome

#### B. Upregulated virulence genes (10T vs 9T)

| | Growth | Spy | gene | Product | FC |
|---|---|---|---|---|---|
| 1 | ME | Spy0137 | *nga* | NAD glycohydrolase | 1.5 |
| 2 | ME | Spy0138 | *ifs* | immunity factor for SPN | 1.6 |
| 3 | ME | Spy0139 | *slo* | streptolysin O | 1.6 |
| 4 | ME | Spy0540 | *sagA* | streptolysin S precursor | 1.6 |
| 5 | ME | Spy1336 | *Spy1336* | R28 protein | 64.7 |
| 6 | ME | Spy1337 | *Spy1337* | transcriptional regulator, AraC family | 4.3 |
| 7 | ME | Spy1675 | *sclA* | collagen-like surface protein A | 2.4 |
| 8 | ME | Spy1699 | - | cell surface protein | 1.6 |
| 9 | ME | Spy1700 | *scpA* | C5A peptidase precursor | 1.7 |
| 10 | ME | Spy1701 | *enn* | enn protein | 1.5 |
| 11 | ME | Spy1702 | *emm* | emm28 protein | 1.7 |
| 1 | ST | Spy0138 | *ifs* | immunity factor for SPN | 1.6 |
| 2 | ST | Spy0139 | *slo* | streptolysin O | 1.5 |
| 3 | ST | Spy1336 | *Spy1336* | R28 protein | 45.3 |
| 4 | ST | Spy1337 | *Spy1337* | transcriptional regulator, AraC family | 7.6 |

#### C. Upregulated virulence genes (11T vs 9T)

| | Growth | Spy | gene | Product | FC |
|---|---|---|---|---|---|
| 1 | ME | Spy0137 | *nga* | NAD glycohydrolase | 1.8 |
| 2 | ME | Spy0138 | *ifs* | immunity factor for SPN | 2.0 |
| 3 | ME | Spy0139 | *slo* | streptolysin O | 1.9 |
| 4 | ME | Spy0329 | *spyCEP* | lactocepin | 2.9 |
| 5 | ME | Spy0540 | *sagA* | streptolysin S precursor | 1.7 |
| 6 | ME | Spy0541 | *sagB* | streptolysin S biosynthesis protein | 1.6 |
| 7 | ME | Spy0542 | *sagC* | streptolysin S biosynthesis protein | 1.6 |
| 8 | ME | Spy0543 | *sagD* | streptolysin S biosynthesis protein | 1.7 |
| 9 | ME | Spy0544 | *sagE* | streptolysin S putative self-immunity protein | 1.8 |
| 10 | ME | Spy0545 | *sagF* | streptolysin S biosynthesis protein | 1.7 |
| 11 | ME | Spy0546 | *sagG* | streptolysin S export ATP-binding protein | 1.8 |
| 12 | ME | Spy0547 | *sagH* | streptolysin S export transmembrane protein | 1.7 |
| 13 | ME | Spy0548 | *sagI* | streptolysin S export transmembrane protein | 1.8 |
| 14 | ME | Spy1336 | *Spy1336* | R28 protein | 115.0 |
| 15 | ME | Spy1337 | *Spy1337* | transcriptional regulator, AraC family | 6.0 |
| 16 | ME | Spy1675 | *sclA* | collagen-like surface protein A | 3.7 |
| 17 | ME | Spy1699 | - | cell surface protein | 2.1 |
| 18 | ME | Spy1700 | *scpA* | C5A peptidase precursor | 2.2 |
| 19 | ME | Spy1701 | *enn* | enn protein | 1.9 |
| 20 | ME | Spy1702 | *emm* | emm28 protein | 1.9 |
| 1 | ST | Spy0137 | *nga* | NAD glycohydrolase | 1.9 |
| 2 | ST | Spy0138 | *ifs* | immunity factor for SPN | 1.8 |
| 3 | ST | Spy0139 | *slo* | streptolysin O | 1.7 |
| 4 | ST | Spy1336 | *Spy1336* | R28 protein | 72.6 |
| 5 | ST | Spy1337 | *Spy1337* | transcriptional regulator, AraC family | 9.1 |
| 6 | ST | Spy1702 | *emm* | emm28 protein | 2.3 |
| 7 | ST | Spy1716 | *sof* | serum opacity factor | 1.5 |

#### D. Upregulated virulence genes (11T vs 10T)

| | Growth | Spy | gene | Product | FC |
|---|---|---|---|---|---|
| 1 | ME | Spy0329 | *spyCEP* | lactocepin | 1.5 |
| 2 | ME | Spy1336 | *Spy1336* | R28 protein | 1.8 |
| 1 | ST | Spy1336 | *Spy1336* | R28 protein | 1.6 |
| 2 | ST | Spy1675 | *sclA* | collagen-like surface protein A | 1.7 |
| 3 | ST | Spy1702 | *emm* | emm28 protein | 1.7 |
| 4 | ST | Spy1621 | *salA* | lantibiotic salivaricin A | 1.8 |

**FIG. S2. RNAseq results comparing isogenic 9T, 10T, and 11T strains**. (**A**) Total number of differentially-expressed genes using a fold cutoff ≥1.5, and *p* value ≤ 0.05. Upregulated virulence genes comparing 10T vs 9T (**B**), 11T vs 9T (**C**), and 11T vs 10T (**D**).

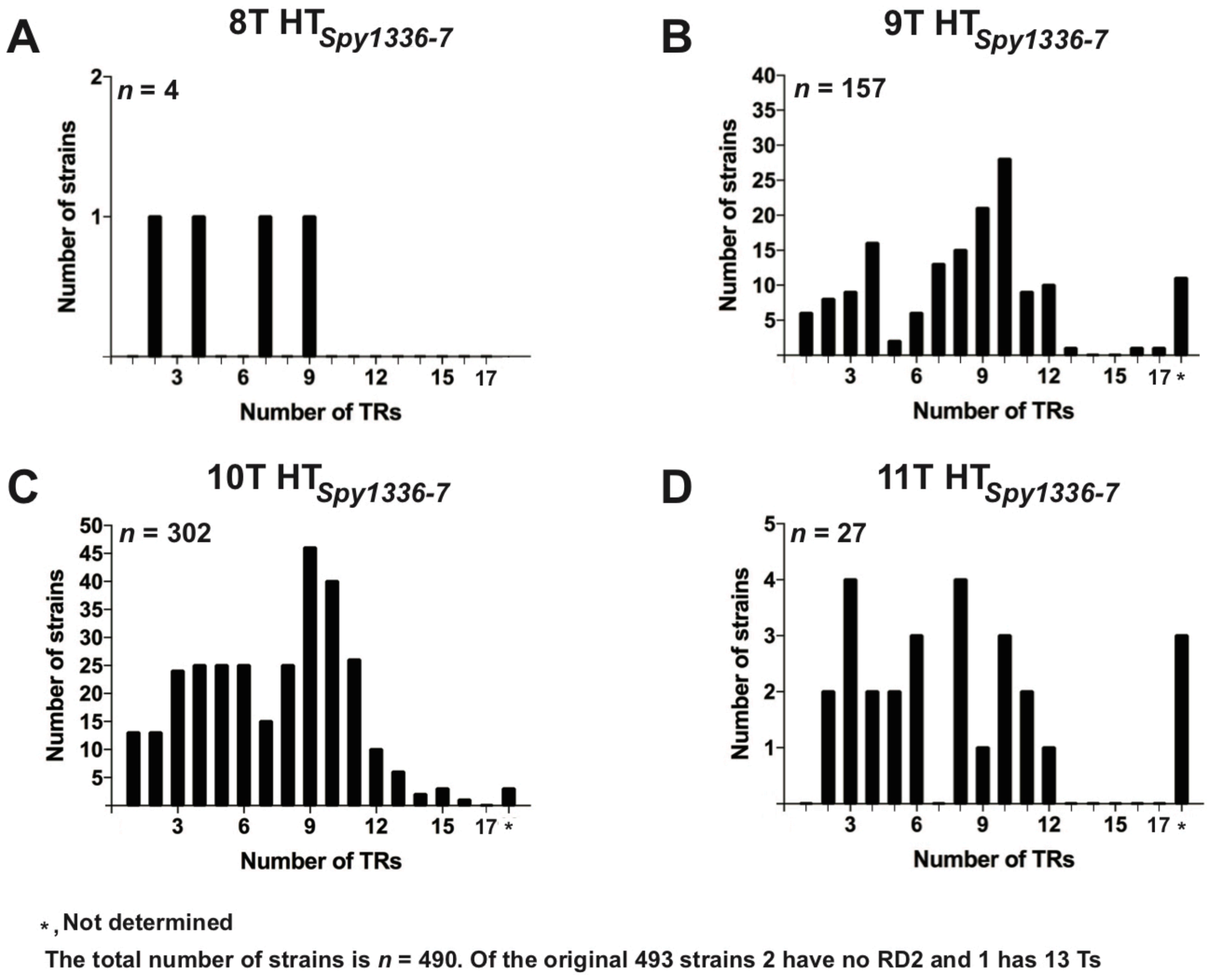


**FIG. S3. Correlation between the number of TR$_{R28}$ repeats in Spy1336/R28 and the number of T nucleotides in the corresponding HT$_{Spy1336\text{-}7}$.** Number of TR$_{R28}$ repeats in strains with HT$_{Spy1336\text{-}7}$ containing 8Ts (**A**), 9Ts (**B**), 10Ts (**C**), and 11Ts (**D**). *n*=490 strains.

**TABLE S1. Number of T nucleotides in $HT_{Spy1336-7}$ in 2,095 *emm28* invasive strains**

| No. | MGAS number[(1)] | Number of Ts[(2)] | Allele number[(3)] | MLST type |
|---|---|---|---|---|
| 1 | **6180** | **10** | 12 | **52** |
| 2 | **7865** | **9** | 2 | **52** |
| 3 | **7866** | **9** | 2 | **52** |
| 4 | **7867** | **9** | 2 | **52** |
| 5 | **7868** | **9** | 2 | **52** |
| 6 | **7869** | **10** | 1 | **52** |
| 7 | **7870** | **10** | 1 | **52** |
| 8 | **7871** | **9** | 2 | **52** |
| 9 | **7872** | **9** | 2 | **52** |
| 10 | **7873** | **10** | 1 | **52** |
| 11 | **7874** | **9** | 2 | **52** |
| 12 | **7876** | **9** | 2 | **52** |
| 13 | **7877** | **10** | 1 | **52** |
| 14 | **7878** | **10** | 1 | **52** |
| 15 | **7879** | **ND[(4)]** | - | **52[(5)]** |
| 16 | **7880** | **9** | 2 | **52** |
| 17 | **7881** | **9** | 2 | **52** |
| 18 | **7882** | **10** | 1 | **52** |
| 19 | **7883** | **10** | 1 | **52** |
| 20 | **7884** | **10** | 1 | **52** |
| 21 | **7885** | **10** | 1 | **52** |
| 22 | **7886** | **10** | 1 | **52** |
| 23 | **7887** | **10** | 1 | **52[(6)]** |
| 24 | **7888** | **10** | 1 | **52** |
| 25 | **7889** | **10** | 1 | **52** |
| 26 | **7890** | **10** | 1 | **52** |
| 27 | **7891** | **9** | 2 | **52** |
| 28 | **7892** | **10** | 1 | **52** |
| 29 | **7893** | **10** | 1 | **52** |
| 30 | **7894** | **10** | 1 | **52** |
| 31 | **7895** | **10** | 1 | **52** |
| 32 | **7896** | **10** | 1 | **52** |
| 33 | **7897** | **10** | 1 | **52** |
| 34 | **7898** | **10** | 1 | **52** |
| 35 | **7899** | **9** | 2 | **52** |
| 36 | **7900** | **10** | 1 | **52** |
| 37 | **7901** | **10** | 1 | **52** |
| 38 | **7902** | **9** | 2 | **52** |
| 39 | **7903** | **9** | 2 | **52** |
| 40 | **7904** | **10** | 1 | **52** |
| 41 | **7905** | **10** | 1 | **52** |
| 42 | **7906** | **10** | 1 | **52** |

| No. | MGAS number(1) | Number of Ts(2) | Allele number(3) | MLST type |
|---|---|---|---|---|
| 43 | **7907** | **10** | 1 | **52** |
| 44 | **7908** | **10** | 1 | **52** |
| 45 | **7909** | **9** | 2 | **52** |
| 46 | **7910** | **9** | 2 | **52** |
| 47 | **7911** | **10** | 1 | **52** |
| 48 | **7912** | **10** | 1 | **52** |
| 49 | **7913** | **10** | 1 | **52** |
| 50 | **7914** | **10** | 1 | **52** |
| 51 | **7916** | **10** | 1 | **52** |
| 52 | **7917** | **10** | 1 | **52** |
| 53 | **7918** | **10** | 1 | **52** |
| 54 | **7919** | **10** | 1 | **52** |
| 55 | **7920** | **10** | 1 | **52** |
| 56 | **7921** | **10** | 1 | **52** |
| 57 | **7922** | **9** | 2 | **52** |
| 58 | **7923** | **10** | 1 | **52** |
| 59 | **7924** | **9** | 2 | **52** |
| 60 | **7925** | **10** | 1 | **52** |
| 61 | **7926** | **10** | 1 | **52** |
| 62 | **7927** | **10** | 1 | **52**(7) |
| 63 | **7928** | **10** | 1 | **52** |
| 64 | **7929** | **11** | 3 | **52** |
| 65 | **7930** | **11** | 3 | **52** |
| 66 | **7931** | **10** | 1 | **52** |
| 67 | **7932** | **10** | 1 | **52** |
| 68 | **7933** | **10** | 1 | **52** |
| 69 | **7934** | **10** | 1 | **52** |
| 70 | **7935** | **10** | 1 | **52** |
| 71 | **7936** | **10** | 1 | **52** |
| 72 | **7937** | **10** | 1 | **52** |
| 73 | **7938** | **10** | 1 | **52** |
| 74 | **7939** | **10** | 1 | **52** |
| 75 | **7940** | **10** | 1 | **52** |
| 76 | **7941** | **10** | 1 | **52** |
| 77 | **7942** | **10** | 1 | **52** |
| 78 | **7943** | **10** | 1 | **52** |
| 79 | **7944** | **9** | 2 | **52** |
| 80 | **7945** | **10** | 1 | **52** |
| 81 | **7946** | **10** | 1 | **52** |
| 82 | **7947** | **10** | 1 | **52** |
| 83 | **7948** | **10** | 1 | **52** |
| 84 | **7949** | **10** | 1 | **52** |
| 85 | **7950** | **10** | 1 | **52**(7) |
| 86 | **7951** | **10** | 1 | **52** |

| No. | MGAS number[1] | Number of Ts[2] | Allele number[3] | MLST type |
|---|---|---|---|---|
| 87 | **7952** | **9** | 2 | **52** |
| 88 | **7953** | **10** | 1 | **52** |
| 89 | **7956** | **10** | 1 | **NF[8,9]** |
| 90 | **7957** | **10** | 1 | **52** |
| 91 | **7958** | **11** | 3 | **52** |
| 92 | **7959** | **10** | 1 | **52** |
| 93 | **7960** | **9** | 2 | **52** |
| 94 | **7961** | **10** | 1 | **52** |
| 95 | **7962** | **10** | 1 | **52** |
| 96 | **7963** | **11** | 3 | **52** |
| 97 | **7964** | **10** | 1 | **52** |
| 98 | **7965** | **9** | 2 | **52** |
| 99 | **7966** | **9** | 2 | **52** |
| 100 | **7967** | **9** | 2 | **52** |
| 101 | **7968** | **10** | 1 | **52** |
| 102 | **7969** | **9** | 2 | **52[10]** |
| 103 | **7970** | **9** | 2 | **52** |
| 104 | **7971** | **10** | 1 | **52** |
| 105 | **7972** | **10** | 1 | **52** |
| 106 | **7973** | **10** | 1 | **52** |
| 107 | **7975** | **9** | 2 | **52** |
| 108 | **7976** | **10** | 1 | **52** |
| 109 | **7977** | **10** | 1 | **52** |
| 110 | **7978** | **10** | 1 | **52** |
| 111 | **7979** | **9** | 2 | **52** |
| 112 | **7980** | **10** | 1 | **52** |
| 113 | **7981** | **10** | 1 | **52[9]** |
| 114 | **7982** | **10** | 1 | **457** |
| 115 | **7983** | **10** | 1 | **52[7]** |
| 116 | **7984** | **10** | 1 | **52** |
| 117 | **7985** | **9** | 2 | **52** |
| 118 | **7986** | **10** | 1 | **52[5]** |
| 119 | **7987** | **10** | 1 | **52** |
| 120 | **7988** | **9** | 2 | **52** |
| 121 | **7989** | **9** | 2 | **52** |
| 122 | **7990** | **10** | 1 | **52** |
| 123 | **7991** | **10** | 1 | **52** |
| 124 | **7993** | **10** | 1 | **52** |
| 125 | **7994** | **10** | 1 | **52** |
| 126 | **7995** | **11** | 3 | **52** |
| 127 | **7996** | **9** | 2 | **52** |
| 128 | **7997** | **9** | 2 | **52** |
| 129 | **7999** | **10** | 1 | **52** |
| 130 | **8000** | **10** | 1 | **52** |

| No. | MGAS number[1] | Number of Ts[2] | Allele number[3] | MLST type |
|---|---|---|---|---|
| 131 | **8001** | **10** | 1 | **52** |
| 132 | **8002** | **10** | 1 | **52** |
| 133 | **8003** | **10** | 1 | **52** |
| 134 | **8004** | **10** | 1 | **52** |
| 135 | **8005** | **10** | 1 | **52** |
| 136 | **8006** | **9** | 2 | **52** |
| 137 | **8007** | **10** | 1 | **52** |
| 138 | **8008** | **9** | 2 | **52** |
| 139 | **8009** | **9** | 2 | **52** |
| 140 | **8010** | **10** | 1 | **52** |
| 141 | **8011** | **9** | 2 | **52** |
| 142 | **8012** | **10** | 1 | **52** |
| 143 | **8013** | **11** | 3 | **52** |
| 144 | **8014** | **12** | 5 | **52** |
| 145 | **8015** | **12** | 5 | **52** |
| 146 | **8016** | **11** | 3 | **52** |
| 147 | **8017** | **10** | 1 | **52** |
| 148 | **8018** | **10** | 1 | **52** |
| 149 | **8019** | **10** | 1 | **52** |
| 150 | **8020** | **9** | 2 | **52** |
| 151 | **8342** | **10** | 1 | **52** |
| 152 | **8343** | **ND** | - | **244** |
| 153 | **8345** | **10** | 1 | **52** |
| 154 | **8346** | **10** | 1 | **52** |
| 155 | **8347** | **9** | 6 | **52** |
| 156 | **8349** | **9** | 2 | **52** |
| 157 | **8350** | **10** | 1 | **52** |
| 158 | **8351** | **10** | 1 | **52** |
| 159 | **8352** | **6** | 20 | **244** |
| 160 | **8353** | **9** | 2 | **52** |
| 161 | **8354** | **9** | 2 | **52** |
| 162 | **8355** | **9** | 2 | **52** |
| 163 | **8356** | **10** | 1 | **52** |
| 164 | **8357** | **10** | 1 | **52** |
| 165 | **8358** | **9** | 2 | **52** |
| 166 | **8359** | **10** | 1 | **52** |
| 167 | **8360** | **10** | 1 | **52** |
| 168 | **8361** | **9** | 6 | **52** |
| 169 | **8362** | **10** | 1 | **52** |
| 170 | **8363** | **11** | 3 | **52** |
| 171 | **8364** | **10** | 1 | **52** |
| 172 | **8365** | **10** | 1 | **52** |
| 173 | **8366** | **10** | 1 | **52** |
| 174 | **8374** | **8** | 4 | **52** |

| No. | MGAS number[1] | Number of Ts[2] | Allele number[3] | MLST type |
|---|---|---|---|---|
| 175 | **8375** | **10** | 1 | **52** |
| 176 | **8376** | **9** | 2 | **52** |
| 177 | **8379** | **9** | 2 | **52** |
| 178 | **8381** | **9** | 2 | **52** |
| 179 | **8383** | **9** | 6 | **52** |
| 180 | **8385** | **9** | 2 | **52** |
| 181 | **8387** | **9** | 2 | **52** |
| 182 | **8389** | **9** | 2 | **52** |
| 183 | **8394** | **10** | 1 | **52** |
| 184 | **8396** | **7** | 7 | **244** |
| 185 | **8405** | **10** | 1 | **52** |
| 186 | **8406** | **9** | 2 | **52** |
| 187 | **8408** | **9** | 2 | **52** |
| 188 | **8410** | **10** | 1 | **52** |
| 189 | **8415** | **10** | 1 | **52** |
| 190 | **8417** | **10** | 1 | **52** |
| 191 | **8423** | **10** | 1 | **52** |
| 192 | **8432** | **10** | 1 | **52** |
| 193 | **8438** | **10** | 1 | **52** |
| 194 | **8439** | **10** | 1 | **52** |
| 195 | **8444** | **11** | 3 | **52** |
| 196 | **8446** | **10** | 1 | **52** |
| 197 | **8447** | **9** | 2 | **52** |
| 198 | **8448** | **9** | 2 | **52** |
| 199 | **10751** | **10** | 1 | **52** |
| 200 | **10752** | **9** | 2 | **52** |
| 201 | **10753** | **10** | 1 | **52** |
| 202 | **10754** | **11** | 3 | **52** |
| 203 | **10755** | **11** | 3 | **52** |
| 204 | **10756** | **9** | 2 | **52** |
| 205 | **10757** | **10** | 1 | **52** |
| 206 | **10758** | **10** | 1 | **52** |
| 207 | **10759** | **10** | 1 | **52** |
| 208 | **10760** | **10** | 1 | **52** |
| 209 | **10761** | **9** | 2 | **52** |
| 210 | **10762** | **10** | 1 | **52** |
| 211 | **10763** | **9** | 2 | **52**[7] |
| 212 | **10764** | **10** | 1 | **52** |
| 213 | **10765** | **10** | 1 | **52** |
| 214 | **10766** | **10** | 1 | **52** |
| 215 | **10767** | **9** | 2 | **52** |
| 216 | **10769** | **11** | 3 | **52** |
| 217 | **10770** | **10** | 1 | **52** |
| 218 | **10771** | **10** | 1 | **52** |

| No. | MGAS number[1] | Number of Ts[2] | Allele number[3] | MLST type |
|---|---|---|---|---|
| 219 | **10772** | **9** | 2 | **52** |
| 220 | **10774** | **10** | 1 | **52** |
| 221 | **10776** | **10** | 1 | **52** |
| 222 | **10777** | **10** | 1 | **52** |
| 223 | **10778** | **10** | 1 | **52** |
| 224 | **10779** | **10** | 1 | **52** |
| 225 | **10780** | **10** | 1 | **52** |
| 226 | **10781** | **10** | 1 | **52** |
| 227 | **10782** | **10** | 1 | **52** |
| 228 | **10783** | **10** | 1 | **52** |
| 229 | **10784** | **10** | 1 | **52** |
| 230 | **10785** | **10** | 1 | **52** |
| 231 | **10786** | **10** | 1 | **52** |
| 232 | **10787** | **10** | 1 | **52** |
| 233 | **10788** | **9** | 2 | **52** |
| 234 | **10789** | **10** | 1 | **52** |
| 235 | **10790** | **10** | 1 | **52** |
| 236 | **10791** | **10** | 1 | **52** |
| 237 | **10792** | **ND** | - | **626** |
| 238 | **10793** | **9** | 2 | **52** |
| 239 | **10794** | **10** | 1 | **52** |
| 240 | **10795** | **10** | 1 | **52** |
| 241 | **10797** | **10** | 1 | **52** |
| 242 | **10798** | **9** | 2 | **52** |
| 243 | **10799** | **10** | 1 | **52** |
| 244 | **10800** | **10** | 1 | **52** |
| 245 | **10801** | **10** | 1 | **52** |
| 246 | **10802** | **10** | 1 | **52** |
| 247 | **10803** | **10** | 1 | **52** |
| 248 | **10804** | **11** | 3 | **52** |
| 249 | **10806** | **9** | 2 | **52** |
| 250 | **10807** | **9** | 2 | **52** |
| 251 | **10808** | **9** | 2 | **52** |
| 252 | **10809** | **9** | 2 | **52** |
| 253 | **10810** | **11** | 3 | **52** |
| 254 | **10811** | **10** | 1 | **52** |
| 255 | **10812** | **9** | 2 | **52** |
| 256 | **10813** | **10** | 1 | **52** |
| 257 | **10814** | **9** | 2 | **52** |
| 258 | **10815** | **9** | 2 | **52** |
| 259 | **10816** | **10** | 1 | **52** |
| 260 | **10817** | **10** | 1 | **52** |
| 261 | **10818** | **9** | 2 | **52** |
| 262 | **10819** | **10** | 1 | **52** |

| No. | MGAS number(1) | Number of Ts(2) | Allele number(3) | MLST type |
|---|---|---|---|---|
| 263 | **10820** | **10** | 1 | **52** |
| 264 | **10821** | **10** | 1 | **52** |
| 265 | **10823** | **9** | 2 | **52** |
| 266 | **10824** | **10** | 1 | **52** |
| 267 | **10825** | **9** | 2 | **52** |
| 268 | **10826** | **9** | 2 | **52** |
| 269 | **10827** | **10** | 1 | **52** |
| 270 | **10828** | **9** | 2 | **52** |
| 271 | **11050** | **10** | 1 | **52** |
| 272 | **11052** | **10** | 1 | **52** |
| 273 | **11053** | **8** | 4 | **52** |
| 274 | **11055** | **10** | 1 | **456** |
| 275 | **11063** | **10** | 1 | **52** |
| 276 | **11064** | **7** | 7 | **244** |
| 277 | **11067** | **9** | 2 | **52** |
| 278 | **11076** | **10** | 1 | **52** |
| 279 | **11080** | **10** | 1 | **52** |
| 280 | **11081** | **10** | 1 | **52** |
| 281 | **11082** | **10** | 1 | **52** |
| 282 | **11083** | **8** | 4 | **52** |
| 283 | **11084** | **10** | 1 | **52** |
| 284 | **11085** | **11** | 3 | **456** |
| 285 | **11086** | **10** | 1 | **52** |
| 286 | **11087** | **9** | 2 | **52** |
| 287 | **11089** | **7** | 21 | **244** |
| 288 | **11090** | **10** | 1 | **52** |
| 289 | **11091** | **10** | 1 | **456** |
| 290 | **11092** | **10** | 1 | **456** |
| 291 | **11093** | **10** | 1 | **456** |
| 292 | **11097** | **ND** | - | **52** |
| 293 | **11098** | **ND** | - | **456** |
| 294 | **11100** | **9** | 2 | **456** |
| 295 | **11102** | **9** | 2 | **52** |
| 296 | **11103** | **9** | 2 | **52** |
| 297 | **11107** | **10** | 1 | **52** |
| 298 | **11108** | **11** | 3 | **456** |
| 299 | **11110** | **10** | 1 | **52** |
| 300 | **11112** | **ND** | - | **52(11)** |
| 301 | **11113** | **10** | 1 | **456** |
| 302 | **11114** | **10** | 1 | **456** |
| 303 | **11115** | **10** | 26 | **456** |
| 304 | **11116** | **10** | 1 | **52** |
| 305 | **11118** | **12** | 5 | **456** |
| 306 | **11119** | **11** | 3 | **52** |

| No. | MGAS number[1] | Number of Ts[2] | Allele number[3] | MLST type |
|---|---|---|---|---|
| 307 | **11120** | **10** | 1 | **52** |
| 308 | **11121** | **10** | 1 | **52** |
| 309 | **11122** | **10** | 1 | **52** |
| 310 | **11123** | **10** | 1 | **456** |
| 311 | **11124** | **10** | 1 | **456** |
| 312 | **11127** | **10** | 1 | **52** |
| 313 | **11128** | **10** | 1 | **456** |
| 314 | **11129** | **9** | 2 | **52** |
| 315 | **11132** | **10** | 1 | **456** |
| 316 | **11133** | **10** | 1 | **456** |
| 317 | **11135** | **10** | 1 | **52** |
| 318 | **11136** | **10** | 1 | **456** |
| 319 | **11138** | **10** | 16 | **456** |
| 320 | **11140** | **10** | 1 | **52** |
| 321 | **11141** | **10** | 1 | **52** |
| 322 | **11142** | **10** | 1 | **52** |
| 323 | **11143** | **10** | 1 | **52** |
| 324 | **11144** | **10** | 1 | **52** |
| 325 | **11145** | **ND** | - | **456** |
| 326 | **11147** | **7** | 7 | **244** |
| 327 | **11149** | **10** | 1 | **52** |
| 328 | **11151** | **7** | 7 | **244** |
| 329 | **11152** | **10** | 1 | **52** |
| 330 | **11153** | **9** | 2 | **52** |
| 331 | **11154** | **10** | 1 | **52** |
| 332 | **11155** | **10** | 1 | **52** |
| 333 | **11156** | **10** | 1 | **52** |
| 334 | **11158** | **9** | 2 | **52** |
| 335 | **12226** | **10** | 1 | **52** |
| 336 | **12227** | **10** | 1 | **52** |
| 337 | **12228** | **9** | 2 | **52** |
| 338 | **12229** | **9** | 2 | **52** |
| 339 | **12230** | **9** | 2 | **52** |
| 340 | **12231** | **10** | 1 | **52** |
| 341 | **12232** | **10** | 1 | **52** |
| 342 | **12233** | **11** | 3 | **52** |
| 343 | **12234** | **9** | 2 | **52** |
| 344 | **12236** | **9** | 11 | **52** |
| 345 | **12237** | **10** | 1 | **52** |
| 346 | **12238** | **10** | 1 | **52** |
| 347 | **12240** | **10** | 1 | **52** |
| 348 | **12241** | **10** | 1 | **52** |
| 349 | **12242** | **9** | 2 | **52** |
| 350 | **12245** | **9** | 2 | **52** |

| No. | MGAS number(1) | Number of Ts(2) | Allele number(3) | MLST type |
|---|---|---|---|---|
| 351 | **12246** | **10** | 1 | **52** |
| 352 | **12247** | **9** | 2 | **52** |
| 353 | **12248** | **9** | 2 | **457** |
| 354 | **12249** | **10** | 1 | **52** |
| 355 | **12250** | **9** | 2 | **52** |
| 356 | **12251** | **10** | 1 | **52** |
| 357 | **12252** | **10** | 1 | **52** |
| 358 | **12253** | **9** | 2 | **52** |
| 359 | **12254** | **10** | 1 | **52** |
| 360 | **12256** | **10** | 1 | **52** |
| 361 | **27720** | **10** | 1 | **52** |
| 362 | **27737** | **10** | 1 | **52** |
| 363 | **27745** | **10** | 1 | **52** |
| 364 | **27754** | **10** | 1 | **52** |
| 365 | **27766** | **ND** | - | **52** |
| 366 | **27767** | **11** | 3 | **52** |
| 367 | **27775** | **10** | 1 | **52** |
| 368 | **27776** | **11** | 3 | **52** |
| 369 | **27784** | **9** | 2 | **52** |
| 370 | **27785** | **10** | 1 | **52** |
| 371 | **27787** | **10** | 1 | **52** |
| 372 | **27798** | **9** | 2 | **52** |
| 373 | **27808** | **11** | 3 | **52** |
| 374 | **27809** | **10** | 1 | **52** |
| 375 | **27813** | **9** | 2 | **52** |
| 376 | **27814** | **10** | 1 | **52** |
| 377 | **27843** | **10** | 1 | **52** |
| 378 | **27844** | **10** | 1 | **52** |
| 379 | **27846** | **10** | 1 | **52** |
| 380 | **27848** | **9** | 2 | **52** |
| 381 | **27859** | **10** | 1 | **52** |
| 382 | **27864** | **9** | 2 | **456** |
| 383 | **27865** | **9** | 2 | **52** |
| 384 | **27866** | **10** | 1 | **456** |
| 385 | **27876** | **9** | 2 | **52** |
| 386 | **27893** | **10** | 1 | **52** |
| 387 | **27895** | **10** | 1 | **52** |
| 388 | **27935** | **9** | 2 | **52** |
| 389 | **27936** | **9** | 2 | **52** |
| 390 | **27937** | **10** | 1 | **52** |
| 391 | **27940** | **9** | 2 | **52** |
| 392 | **27941** | **10** | 1 | **52** |
| 393 | **27942** | **9** | 2 | **52** |
| 394 | **27943** | **10** | 1 | **52** |

| No. | MGAS number[1] | Number of Ts[2] | Allele number[3] | MLST type |
|---|---|---|---|---|
| 395 | **27944** | **10** | 1 | **52** |
| 396 | **27945** | **10** | 1 | **52** |
| 397 | **27946** | **10** | 1 | **52** |
| 398 | **27947** | **10** | 1 | **52** |
| 399 | **27948** | **10** | 1 | **52** |
| 400 | **27949** | **10** | 1 | **52**[12] |
| 401 | **27950** | **10** | 1 | **52** |
| 402 | **27951** | **10** | 1 | **52** |
| 403 | **27952** | **10** | 1 | **52** |
| 404 | **27954** | **10** | 1 | **52** |
| 405 | **27955** | **9** | 2 | **52** |
| 406 | **27956** | **10** | 1 | **52** |
| 407 | **27957** | **10** | 1 | **52** |
| 408 | **27958** | **11** | 3 | **52** |
| 409 | **27959** | **10** | 1 | **52** |
| 410 | **27960** | **10** | 1 | **52** |
| 411 | **27961** | **9** | 2 | **52** |
| 412 | **27962** | **10** | 1 | **52** |
| 413 | **27963** | **10** | 1 | **52** |
| 414 | **27964** | **10** | 1 | **52** |
| 415 | **27965** | **10** | 1 | **52** |
| 416 | **27966** | **10** | 1 | **52** |
| 417 | **27967** | **10** | 1 | **52** |
| 418 | **27968** | **10** | 1 | **52** |
| 419 | **27969** | **10** | 1 | **52** |
| 420 | **27970** | **9** | 2 | **52** |
| 421 | **27972** | **9** | 2 | **52** |
| 422 | **27973** | **10** | 1 | **52** |
| 423 | **27974** | **10** | 1 | **52** |
| 424 | **27975** | **9** | 2 | **52** |
| 425 | **27976** | **10** | 1 | **52** |
| 426 | **27977** | **10** | 1 | **52** |
| 427 | **27978** | **9** | 2 | **52** |
| 428 | **27979** | **10** | 1 | **52**[7] |
| 429 | **27980** | **10** | 1 | **52** |
| 430 | **27981** | **9** | 2 | **52** |
| 431 | **27982** | **9** | 2 | **52** |
| 432 | **27983** | **11** | 3 | **52** |
| 433 | **27984** | **10** | 1 | **52** |
| 434 | **27985** | **10** | 1 | **52** |
| 435 | **27986** | **10** | 1 | **52** |
| 436 | **27987** | **10** | 1 | **52** |
| 437 | **27988** | **10** | 1 | **52** |
| 438 | **27989** | **11** | 3 | **52** |

| No. | MGAS number(1) | Number of Ts(2) | Allele number(3) | MLST type |
|---|---|---|---|---|
| 439 | **27990** | **10** | 1 | **52** |
| 440 | **27991** | **9** | 2 | **52** |
| 441 | **27992** | **9** | 2 | **52** |
| 442 | **27993** | **9** | 2 | **52** |
| 443 | **27994** | **10** | 1 | **52(7)** |
| 444 | **27995** | **10** | 1 | **52** |
| 445 | **27996** | **10** | 10 | **52** |
| 446 | **27997** | **10** | 1 | **52** |
| 447 | **27998** | **10** | 1 | **52** |
| 448 | **27999** | **9** | 2 | **52** |
| 449 | **28000** | **9** | 2 | **52** |
| 450 | **28001** | **9** | 2 | **52** |
| 451 | **28002** | **9** | 2 | **52** |
| 452 | **28003** | **10** | 1 | **52** |
| 453 | **28004** | **9** | 2 | **52** |
| 454 | **28005** | **9** | 2 | **52** |
| 455 | **28006** | **9** | 2 | **52** |
| 456 | **28007** | **9** | 2 | **52** |
| 457 | **28008** | **10** | 1 | **52** |
| 458 | **28009** | **9** | 2 | **52** |
| 459 | **28010** | **10** | 1 | **52** |
| 460 | **28011** | **10** | 1 | **52** |
| 461 | **28012** | **9** | 6 | **52** |
| 462 | **28013** | **10** | 1 | **52** |
| 463 | **28014** | **10** | 1 | **52** |
| 464 | **28015** | **10** | 1 | **52** |
| 465 | **28016** | **10** | 1 | **52** |
| 466 | **28018** | **10** | 1 | **52** |
| 467 | **28019** | **9** | 2 | **52** |
| 468 | **28020** | **10** | 1 | **52** |
| 469 | **28021** | **10** | 1 | **52** |
| 470 | **28022** | **10** | 1 | **52** |
| 471 | **28023** | **9** | 2 | **52** |
| 472 | **28024** | **10** | 1 | **52** |
| 473 | **28025** | **10** | 1 | **52** |
| 474 | **28026** | **10** | 1 | **52** |
| 475 | **28027** | **10** | 1 | **52** |
| 476 | **28028** | **10** | 1 | **52** |
| 477 | **28029** | **10** | 1 | **52** |
| 478 | **28030** | **10** | 1 | **52** |
| 479 | **28031** | **9** | 2 | **52** |
| 480 | **28032** | **10** | 1 | **52** |
| 481 | **28033** | **9** | 2 | **52** |
| 482 | **28034** | **12** | 5 | **52** |

| No. | MGAS number[1] | Number of Ts[2] | Allele number[3] | MLST type |
|---|---|---|---|---|
| 483 | **28035** | **10** | 1 | **52** |
| 484 | **28036** | **9** | 2 | **52** |
| 485 | **28037** | **9** | 2 | **52** |
| 486 | **28038** | **9** | 2 | **52** |
| 487 | **28039** | **10** | 1 | **52**[13] |
| 488 | **28040** | **10** | 1 | **52** |
| 489 | **28041** | **9** | 2 | **52** |
| 490 | **28042** | **10** | 1 | **52** |
| 491 | **28043** | **9** | 2 | **52** |
| 492 | **28044** | **ND** | - | **52** |
| 493 | **28045** | **10** | 1 | **52** |
| 494 | **28046** | **10** | 1 | **52** |
| 495 | **28047** | **10** | 1 | **52** |
| 496 | **28048** | **9** | 2 | **52** |
| 497 | **28049** | **8** | 8 | **52** |
| 498 | **28050** | **9** | 2 | **52**[7] |
| 499 | **28051** | **9** | 2 | **52** |
| 500 | **28052** | **9** | 2 | **52** |
| 501 | **28053** | **10** | 1 | **52** |
| 502 | **28054** | **10** | 1 | **52** |
| 503 | **28055** | **10** | 1 | **52**[13] |
| 504 | **28056** | **9** | 2 | **52** |
| 505 | **28057** | **10** | 1 | **52** |
| 506 | **28058** | **9** | 2 | **52** |
| 507 | **28059** | **ND** | - | **52** |
| 508 | **28060** | **10** | 1 | **52** |
| 509 | **28061** | **9** | 2 | **52** |
| 510 | **28062** | **10** | 1 | **52** |
| 511 | **28063** | **10** | 1 | **52** |
| 512 | **28064** | **10** | 1 | **52** |
| 513 | **28065** | **10** | 1 | **52** |
| 514 | **28066** | **10** | 1 | **52** |
| 515 | **28067** | **11** | 3 | **52** |
| 516 | **28068** | **10** | 1 | **52** |
| 517 | **28069** | **10** | 1 | **52** |
| 518 | **28070** | **10** | 1 | **52** |
| 519 | **28071** | **10** | 1 | **52** |
| 520 | **28072** | **10** | 1 | **52** |
| 521 | **28073** | **11** | 3 | **52** |
| 522 | **28074** | **10** | 1 | **52** |
| 523 | **28075** | **10** | 1 | **52** |
| 524 | **28076** | **9** | 2 | **52** |
| 525 | **28077** | **10** | 1 | **52** |
| 526 | **28078** | **10** | 1 | **52** |

| No. | MGAS number[1] | Number of Ts[2] | Allele number[3] | MLST type |
|---|---|---|---|---|
| 527 | **28079** | **10** | 1 | **52** |
| 528 | **28081** | **9** | 2 | **52** |
| 529 | **28082** | **9** | 2 | **52** |
| 530 | **28083** | **10** | 1 | **52** |
| 531 | **28084** | **11** | 3 | **52** |
| 532 | **28085** | **10** | 1 | **52** |
| 533 | **28086** | **10** | 1 | **52** |
| 534 | **28087** | **9** | 2 | **52** |
| 535 | **28088** | **9** | 2 | **52** |
| 536 | **28089** | **11** | 3 | **52** |
| 537 | **28090** | **11** | 3 | **52** |
| 538 | **28091** | **10** | 1 | **52** |
| 539 | **28092** | **9** | 2 | **52** |
| 540 | **28093** | **9** | 2 | **52** |
| 541 | **28094** | **10** | 1 | **52** |
| 542 | **28095** | **9** | 2 | **52** |
| 543 | **28096** | **10** | 1 | **52** |
| 544 | **28097** | **11** | 3 | **52** |
| 545 | **28098** | **11** | 3 | **52** |
| 546 | **28099** | **9** | 2 | **52** |
| 547 | **28100** | **9** | 2 | **52** |
| 548 | **28101** | **10** | 1 | **52** |
| 549 | **28102** | **10** | 1 | **52** |
| 550 | **28103** | **10** | 1 | **456** |
| 551 | **28104** | **10** | 1 | **52** |
| 552 | **28105** | **10** | 1 | **52** |
| 553 | **28106** | **10** | 1 | **52** |
| 554 | **28107** | **10** | 1 | **52** |
| 555 | **28108** | **9** | 2 | **52** |
| 556 | **28109** | **9** | 2 | **52** |
| 557 | **28111** | **10** | 1 | **52** |
| 558 | **28112** | **9** | 2 | **52** |
| 559 | **28113** | **10** | 1 | **52** |
| 560 | **28114** | **9** | 2 | **52** |
| 561 | **28115** | **10** | 1 | **52** |
| 562 | **28116** | **10** | 1 | **52** |
| 563 | **28117** | **9** | 2 | **52** |
| 564 | **28118** | **9** | 2 | **52** |
| 565 | **28119** | **9** | 2 | **52** |
| 566 | **28121** | **10** | 30 | **52** |
| 567 | **28122** | **9** | 2 | **52** |
| 568 | **28123** | **10** | 1 | **52** |
| 569 | **28127** | **9** | 2 | **52** |
| 570 | **28128** | **10** | 1 | **52** |

| No. | MGAS number(1) | Number of Ts(2) | Allele number(3) | MLST type |
|---|---|---|---|---|
| 571 | **28129** | **10** | 1 | **52** |
| 572 | **28130** | **10** | 1 | **52** |
| 573 | **28131** | **10** | 1 | **52** |
| 574 | **28132** | **9** | 2 | **52** |
| 575 | **28133** | **10** | 1 | **52(12)** |
| 576 | **28136** | **10** | 1 | **52** |
| 577 | **28137** | **10** | 1 | **52** |
| 578 | **28138** | **9** | 2 | **52** |
| 579 | **28140** | **9** | 2 | **52** |
| 580 | **28141** | **ND** | - | **52** |
| 581 | **28142** | **10** | 1 | **52** |
| 582 | **28143** | **10** | 1 | **52(14)** |
| 583 | **28144** | **9** | 2 | **52** |
| 584 | **28145** | **10** | 1 | **52(14)** |
| 585 | **28148** | **10** | 1 | **52** |
| 586 | **28152** | **9** | 2 | **52** |
| 587 | **28154** | **9** | 2 | **52** |
| 588 | **28155** | **9** | 2 | **52** |
| 589 | **28156** | **11** | 3 | **52** |
| 590 | **28157** | **10** | 1 | **52** |
| 591 | **28158** | **9** | 2 | **52** |
| 592 | **28159** | **10** | 1 | **52** |
| 593 | **28160** | **10** | 1 | **52** |
| 594 | **28161** | **10** | 1 | **52** |
| 595 | **28162** | **10** | 1 | **52** |
| 596 | **28163** | **10** | 1 | **52** |
| 597 | **28164** | **11** | 3 | **52** |
| 598 | **28165** | **11** | 3 | **52** |
| 599 | **28166** | **10** | 1 | **52** |
| 600 | **28167** | **9** | 2 | **52** |
| 601 | **28168** | **11** | 3 | **52** |
| 602 | **28172** | **9** | 2 | **52** |
| 603 | **28173** | **11** | 3 | **52** |
| 604 | **28174** | **10** | 1 | **52** |
| 605 | **28175** | **9** | 2 | **52** |
| 606 | **28176** | **9** | 2 | **52** |
| 607 | **28177** | **10** | 1 | **52** |
| 608 | **28182** | **10** | 1 | **52** |
| 609 | **28183** | **9** | 2 | **52** |
| 610 | **28184** | **10** | 1 | **52** |
| 611 | **28185** | **10** | 1 | **52** |
| 612 | **28186** | **10** | 1 | **52** |
| 613 | **28188** | **10** | 1 | **52** |
| 614 | **28189** | **9** | 2 | **52** |

| No. | MGAS number[1] | Number of Ts[2] | Allele number[3] | MLST type |
|---|---|---|---|---|
| 615 | **28190** | **9** | 2 | **52** |
| 616 | **28191** | **10** | 1 | **52** |
| 617 | **28192** | **10** | 1 | **52** |
| 618 | **28195** | **10** | 1 | **52** |
| 619 | **28196** | **10** | 1 | **52** |
| 620 | **28197** | **10** | 1 | **52** |
| 621 | **28200** | **10** | 1 | **52** |
| 622 | **28201** | **10** | 1 | **52** |
| 623 | **28202** | **10** | 1 | **52** |
| 624 | **28203** | **9** | 2 | **52** |
| 625 | **28204** | **9** | 2 | **52** |
| 626 | **28206** | **10** | 1 | **52** |
| 627 | **28207** | **9** | 2 | **52** |
| 628 | **28208** | **9** | 19 | **52** |
| 629 | **28209** | **10** | 1 | **52** |
| 630 | **28210** | **9** | 2 | **52** |
| 631 | **28211** | **9** | 2 | **52** |
| 632 | **28212** | **9** | 2 | **52** |
| 633 | **28213** | **10** | 1 | **52** |
| 634 | **28214** | **10** | 1 | **52**[13] |
| 635 | **28215** | **9** | 2 | **52** |
| 636 | **28216** | **9** | 2 | **52** |
| 637 | **28217** | **10** | 1 | **52** |
| 638 | **28218** | **9** | 2 | **52** |
| 639 | **28219** | **9** | 2 | **52** |
| 640 | **28220** | **9** | 2 | **52** |
| 641 | **28221** | **9** | 2 | **52** |
| 642 | **28222** | **10** | 1 | **52** |
| 643 | **28223** | **10** | 1 | **52** |
| 644 | **28224** | **9** | 2 | **52** |
| 645 | **28225** | **9** | 2 | **52**[7] |
| 646 | **28226** | **9** | 2 | **52** |
| 647 | **28227** | **9** | 2 | **52** |
| 648 | **28228** | **9** | 2 | **52** |
| 649 | **28229** | **9** | 2 | **52** |
| 650 | **28230** | **10** | 1 | **52** |
| 651 | **28231** | **9** | 2 | **52** |
| 652 | **28232** | **11** | 3 | **52** |
| 653 | **28233** | **9** | 2 | **52** |
| 654 | **28234** | **10** | 1 | **52** |
| 655 | **28235** | **10** | 1 | **52** |
| 656 | **28236** | **10** | 1 | **52** |
| 657 | **28237** | **9** | 2 | **52** |
| 658 | **28238** | **9** | 2 | **52** |

| No. | MGAS number(1) | Number of Ts(2) | Allele number(3) | MLST type |
|---|---|---|---|---|
| 659 | **28239** | **10** | 1 | **52** |
| 660 | **28240** | **10** | 1 | **52** |
| 661 | **28241** | **9** | 2 | **52** |
| 662 | **28243** | **9** | 2 | **52** |
| 663 | **28244** | **9** | 2 | **52** |
| 664 | **28245** | **10** | 1 | **52** |
| 665 | **28246** | **10** | 1 | **52** |
| 666 | **28247** | **10** | 1 | **52** |
| 667 | **28248** | **10** | 1 | **52** |
| 668 | **28249** | **11** | 3 | **52** |
| 669 | **28250** | **11** | 3 | **52** |
| 670 | **28251** | **11** | 3 | **52** |
| 671 | **28252** | **9** | 2 | **52** |
| 672 | **28253** | **9** | 2 | **52** |
| 673 | **28254** | **9** | 2 | **52** |
| 674 | **28255** | **9** | 2 | **52** |
| 675 | **28256** | **10** | 1 | **52** |
| 676 | **28257** | **9** | 2 | **52** |
| 677 | **28258** | **10** | 1 | **52** |
| 678 | **28259** | **11** | 3 | **52** |
| 679 | **28260** | **10** | 1 | **52** |
| 680 | **28261** | **10** | 1 | **52** |
| 681 | **28263** | **9** | 2 | **52** |
| 682 | **28264** | **10** | 1 | **52** |
| 683 | **28265** | **10** | 1 | **52** |
| 684 | **28266** | **10** | 1 | **52** |
| 685 | **28267** | **10** | 1 | **52** |
| 686 | **28268** | **10** | 1 | **52** |
| 687 | **28269** | **10** | 1 | **52** |
| 688 | **28270** | **9** | 2 | **52** |
| 689 | **28271** | **10** | 1 | **52** |
| 690 | **28272** | **10** | 1 | **52** |
| 691 | **28273** | **11** | 3 | **52** |
| 692 | **28274** | **10** | 1 | **52** |
| 693 | **28275** | **10** | 1 | **52** |
| 694 | **28276** | **11** | 3 | **52** |
| 695 | **28277** | **10** | 1 | **52** |
| 696 | **28278** | **10** | 1 | **52** |
| 697 | **28279** | **10** | 1 | **52** |
| 698 | **28280** | **9** | 2 | **52(13)** |
| 699 | **28281** | **9** | 2 | **52** |
| 700 | **28282** | **11** | 3 | **52** |
| 701 | **28283** | **10** | 1 | **52** |
| 702 | **28284** | **10** | 1 | **52** |

| No. | MGAS number(1) | Number of Ts(2) | Allele number(3) | MLST type |
|---|---|---|---|---|
| 703 | **28285** | **9** | 2 | **52**(9) |
| 704 | **28286** | **10** | 1 | **52** |
| 705 | **28287** | **10** | 1 | **52** |
| 706 | **28288** | **9** | 2 | **52** |
| 707 | **28290** | **9** | 2 | **52** |
| 708 | **28291** | **10** | 1 | **52** |
| 709 | **28292** | **9** | 2 | **52** |
| 710 | **28294** | **9** | 2 | **52** |
| 711 | **28295** | **10** | 1 | **52** |
| 712 | **28296** | **10** | 1 | **52** |
| 713 | **28297** | **10** | 1 | **52** |
| 714 | **28298** | **9** | 11 | **52** |
| 715 | **28299** | **9** | 2 | **52** |
| 716 | **28300** | **10** | 1 | **52** |
| 717 | **28302** | **9** | 2 | **52**(15) |
| 718 | **28303** | **10** | 1 | **52** |
| 719 | **28304** | **10** | 1 | **52** |
| 720 | **28305** | **10** | 1 | **52** |
| 721 | **28306** | **12** | 5 | **52** |
| 722 | **28307** | **9** | 2 | **52** |
| 723 | **28308** | **11** | 3 | **52** |
| 724 | **28309** | **9** | 2 | **52** |
| 725 | **28310** | **10** | 1 | **52** |
| 726 | **28311** | **9** | 2 | **52** |
| 727 | **28312** | **10** | 1 | **52** |
| 728 | **28313** | **9** | 2 | **52**(13) |
| 729 | **28314** | **10** | 1 | **52** |
| 730 | **28315** | **9** | 2 | **52** |
| 731 | **28316** | **11** | 3 | **52** |
| 732 | **28317** | **10** | 1 | **52** |
| 733 | **28318** | **9** | 2 | **52** |
| 734 | **28319** | **10** | 1 | **52** |
| 735 | **28320** | **10** | 1 | **52** |
| 736 | **28321** | **10** | 1 | **52** |
| 737 | **28322** | **10** | 1 | **52** |
| 738 | **28323** | **10** | 1 | **52** |
| 739 | **28324** | **10** | 1 | **52** |
| 740 | **28325** | **10** | 1 | **52** |
| 741 | **28327** | **9** | 2 | **52** |
| 742 | **28328** | **10** | 1 | **52** |
| 743 | **28329** | **10** | 1 | **52** |
| 744 | **28330** | **10** | 1 | **52** |
| 745 | **28331** | **10** | 1 | **52** |
| 746 | **28332** | **10** | 1 | **52** |

| No. | MGAS number[1] | Number of Ts[2] | Allele number[3] | MLST type |
|---|---|---|---|---|
| 747 | **28333** | **9** | 2 | **52** |
| 748 | **28334** | **10** | 1 | **52** |
| 749 | **28335** | **10** | 1 | **52** |
| 750 | **28336** | **10** | 1 | **52** |
| 751 | **28337** | **11** | 3 | **52** |
| 752 | **28338** | **10** | 1 | **52** |
| 753 | **28339** | **10** | 1 | **52** |
| 754 | **28340** | **9** | 2 | **52** |
| 755 | **28341** | **10** | 1 | **52** |
| 756 | **28342** | **9** | 2 | **52** |
| 757 | **28343** | **9** | 2 | **52** |
| 758 | **28345** | **10** | 1 | **52**[7] |
| 759 | **28346** | **9** | 2 | **52** |
| 760 | **28347** | **10** | 1 | **52** |
| 761 | **28348** | **9** | 2 | **52** |
| 762 | **28350** | **10** | 1 | **52** |
| 763 | **28351** | **10** | 1 | **52**[13] |
| 764 | **28353** | **9** | 2 | **52** |
| 765 | **28354** | **10** | 1 | **52** |
| 766 | **28355** | **10** | 1 | **52** |
| 767 | **28356** | **9** | 2 | **52** |
| 768 | **28357** | **9** | 2 | **52** |
| 769 | **28358** | **9** | 2 | **52** |
| 770 | **28359** | **9** | 2 | **52** |
| 771 | **28360** | **9** | 2 | **52** |
| 772 | **28361** | **10** | 1 | **52** |
| 773 | **28362** | **10** | 1 | **52** |
| 774 | **28363** | **9** | 2 | **52** |
| 775 | **28364** | **10** | 1 | **52** |
| 776 | **28365** | **10** | 1 | **52** |
| 777 | **28366** | **9** | 2 | **52** |
| 778 | **28367** | **10** | 1 | **52** |
| 779 | **28368** | **10** | 1 | **52** |
| 780 | **28369** | **10** | 1 | **52** |
| 781 | **28370** | **10** | 1 | **52** |
| 782 | **28371** | **9** | 2 | **52** |
| 783 | **28372** | **10** | 1 | **52** |
| 784 | **28373** | **9** | 2 | **52** |
| 785 | **28374** | **9** | 2 | **52** |
| 786 | **28375** | **9** | 2 | **52** |
| 787 | **28376** | **9** | 2 | **52** |
| 788 | **28378** | **9** | 2 | **52** |
| 789 | **28379** | **10** | 1 | **52** |
| 790 | **28380** | **9** | 2 | **52** |

| No. | MGAS number[1] | Number of Ts[2] | Allele number[3] | MLST type |
|---|---|---|---|---|
| 791 | **28381** | **9** | 2 | **456** |
| 792 | **28382** | **10** | 1 | **52** |
| 793 | **28383** | **10** | 1 | **52** |
| 794 | **28384** | **10** | 1 | **52** |
| 795 | **28385** | **9** | 15 | **52** |
| 796 | **28386** | **11** | 3 | **52** |
| 797 | **28387** | **10** | 1 | **52** |
| 798 | **28388** | **10** | 1 | **52** |
| 799 | **28389** | **10** | 1 | **52** |
| 800 | **28390** | **10** | 1 | **52** |
| 801 | **28391** | **10** | 1 | **52** |
| 802 | **28392** | **10** | 1 | **52** |
| 803 | **28393** | **9** | 2 | **52** |
| 804 | **28394** | **10** | 1 | **52** |
| 805 | **28395** | **9** | 18 | **52** |
| 806 | **28396** | **10** | 1 | **52** |
| 807 | **28397** | **9** | 2 | **52** |
| 808 | **28398** | **10** | 1 | **52** |
| 809 | **28400** | **ND** | - | **52** |
| 810 | **28401** | **10** | 1 | **52** |
| 811 | **28402** | **10** | 1 | **52** |
| 812 | **28403** | **10** | 1 | **52**[12] |
| 813 | **28404** | **11** | 3 | **52** |
| 814 | **28406** | **10** | 1 | **52** |
| 815 | **28409** | **9** | 2 | **52** |
| 816 | **28410** | **8** | 4 | **52** |
| 817 | **28411** | **10** | 1 | **52** |
| 818 | **28412** | **9** | 2 | **52** |
| 819 | **28414** | **9** | 2 | **52** |
| 820 | **28415** | **ND** | - | **52** |
| 821 | **28416** | **9** | 2 | **52** |
| 822 | **28417** | **11** | 3 | **52** |
| 823 | **28418** | **10** | 1 | **52** |
| 824 | **28419** | **10** | 1 | **52** |
| 825 | **28420** | **10** | 1 | **52** |
| 826 | **28421** | **10** | 1 | **52** |
| 827 | **28422** | **10** | 1 | **52** |
| 828 | **28423** | **10** | 1 | **52** |
| 829 | **28424** | **8** | 8 | **52** |
| 830 | **28425** | **10** | 1 | **52** |
| 831 | **28426** | **9** | 2 | **52** |
| 832 | **28428** | **10** | 1 | **52** |
| 833 | **28429** | **9** | 2 | **52** |
| 834 | **28430** | **10** | 1 | **52** |

| No. | MGAS number(1) | Number of Ts(2) | Allele number(3) | MLST type |
|---|---|---|---|---|
| 835 | **28431** | **11** | 3 | **52** |
| 836 | **28433** | **10** | 1 | **52** |
| 837 | **28434** | **9** | 2 | **52** |
| 838 | **28435** | **9** | 2 | **52** |
| 839 | **28436** | **9** | 2 | **52** |
| 840 | **28437** | **10** | 1 | **52** |
| 841 | **28438** | **10** | 1 | **52** |
| 842 | **28439** | **10** | 1 | **52** |
| 843 | **28440** | **10** | 1 | **52** |
| 844 | **28441** | **9** | 2 | **52** |
| 845 | **28442** | **10** | 25 | **52** |
| 846 | **28443** | **9** | 2 | **52** |
| 847 | **28444** | **11** | 3 | **52** |
| 848 | **28445** | **9** | 2 | **52** |
| 849 | **28446** | **10** | 1 | **52** |
| 850 | **28447** | **9** | 2 | **52** |
| 851 | **28448** | **9** | 2 | **52** |
| 852 | **28449** | **10** | 1 | **52** |
| 853 | **28450** | **10** | 1 | **52** |
| 854 | **28451** | **10** | 1 | **52** |
| 855 | **28452** | **10** | 1 | **52** |
| 856 | **28453** | **10** | 1 | **52** |
| 857 | **28454** | **9** | 2 | **52** |
| 858 | **28455** | **10** | 1 | **52** |
| 859 | **28456** | **ND** | - | **52** |
| 860 | **28457** | **9** | 2 | **52** |
| 861 | **28458** | **9** | 2 | **52** |
| 862 | **28459** | **10** | 1 | **52** |
| 863 | **28460** | **9** | 2 | **52** |
| 864 | **28461** | **10** | 1 | **52** |
| 865 | **28462** | **10** | 1 | **52** |
| 866 | **28463** | **10** | 1 | **52** |
| 867 | **28464** | **10** | 1 | **52** |
| 868 | **28465** | **9** | 2 | **52** |
| 869 | **28466** | **11** | 27 | **52** |
| 870 | **28467** | **10** | 1 | **52** |
| 871 | **28468** | **11** | 3 | **52** |
| 872 | **28469** | **10** | 1 | **52** |
| 873 | **28470** | **9** | 2 | **52** |
| 874 | **28471** | **10** | 1 | **52** |
| 875 | **28472** | **10** | 1 | **52** |
| 876 | **28473** | **10** | 1 | **52** |
| 877 | **28474** | **9** | 2 | **52** |
| 878 | **28475** | **10** | 1 | **52** |

| No. | MGAS number[1] | Number of Ts[2] | Allele number[3] | MLST type |
|---|---|---|---|---|
| 879 | **28476** | **10** | 1 | **52** |
| 880 | **28477** | **10** | 1 | **52** |
| 881 | **28478** | **10** | 1 | **52** |
| 882 | **28479** | **10** | 1 | **52** |
| 883 | **28480** | **10** | 1 | **52** |
| 884 | **28481** | **9** | 2 | **52** |
| 885 | **28482** | **10** | 1 | **52** |
| 886 | **28483** | **9** | 2 | **52** |
| 887 | **28484** | **10** | 1 | **52** |
| 888 | **28485** | **10** | 1 | **52** |
| 889 | **28486** | **10** | 1 | **52** |
| 890 | **28487** | **9** | 2 | **52** |
| 891 | **28490** | **9** | 2 | **52** |
| 892 | **28491** | **9** | 2 | **52** |
| 893 | **28492** | **9** | 2 | **52** |
| 894 | **28493** | **11** | 3 | **52** |
| 895 | **28494** | **9** | 2 | **52** |
| 896 | **28495** | **10** | 1 | **52**[7] |
| 897 | **28496** | **9** | 2 | **52** |
| 898 | **28498** | **10** | 1 | **52** |
| 899 | **28499** | **9** | 2 | **52** |
| 900 | **28500** | **9** | 2 | **52** |
| 901 | **28501** | **10** | 1 | **52** |
| 902 | **28502** | **10** | 1 | **52** |
| 903 | **28503** | **9** | 2 | **52**[13] |
| 904 | **28504** | **9** | 2 | **52** |
| 905 | **28505** | **10** | 1 | **52** |
| 906 | **28506** | **10** | 1 | **52** |
| 907 | **28507** | **10** | 1 | **52** |
| 908 | **28508** | **9** | 2 | **52** |
| 909 | **28509** | **9** | 2 | **52** |
| 910 | **28510** | **10** | 1 | **52** |
| 911 | **28511** | **10** | 1 | **52** |
| 912 | **28512** | **11** | 3 | **52** |
| 913 | **28513** | **10** | 1 | **52** |
| 914 | **28514** | **9** | 2 | **52** |
| 915 | **28515** | **9** | 2 | **52** |
| 916 | **28516** | **10** | 1 | **52** |
| 917 | **28517** | **11** | 3 | **52** |
| 918 | **28518** | **10** | 1 | **52** |
| 919 | **28519** | **9** | 2 | **52**[14] |
| 920 | **28520** | **10** | 1 | **52** |
| 921 | **28521** | **10** | 1 | **52** |
| 922 | **28522** | **11** | 3 | **52** |

| No. | MGAS number[1] | Number of Ts[2] | Allele number[3] | MLST type |
|---|---|---|---|---|
| 923 | **28523** | **10** | 1 | **52** |
| 924 | **28524** | **10** | 1 | **52** |
| 925 | **28525** | **10** | 1 | **52** |
| 926 | **28527** | **10** | 1 | **52** |
| 927 | **28528** | **10** | 1 | **52** |
| 928 | **28529** | **10** | 1 | **52** |
| 929 | **28530** | **10** | 1 | **52** |
| 930 | **28531** | **10** | 1 | **52** |
| 931 | **28532** | **10** | 1 | **52** |
| 932 | **28533** | **10** | 1 | **52** |
| 933 | **28534** | **9** | 2 | **52** |
| 934 | **28535** | **10** | 1 | **52** |
| 935 | **28536** | **9** | 2 | **52** |
| 936 | **28537** | **9** | 2 | **52** |
| 937 | **28538** | **10** | 1 | **52** |
| 938 | **28539** | **10** | 1 | **52** |
| 939 | **28540** | **10** | 1 | **52** |
| 940 | **28542** | **10** | 1 | **52** |
| 941 | **28543** | **10** | 1 | **52** |
| 942 | **28544** | **10** | 1 | **52** |
| 943 | **28545** | **10** | 1 | **52** |
| 944 | **28546** | **10** | 1 | **52** |
| 945 | **28547** | **10** | 1 | **52** |
| 946 | **28548** | **10** | 1 | **52** |
| 947 | **28549** | **9** | 2 | **52** |
| 948 | **28550** | **9** | 2 | **52** |
| 949 | **28551** | **11** | 3 | **52**[13] |
| 950 | **28552** | **10** | 1 | **52** |
| 951 | **28553** | **11** | 3 | **52** |
| 952 | **28554** | **9** | 2 | **52** |
| 953 | **28555** | **10** | 1 | **52** |
| 954 | **28556** | **10** | 1 | **52** |
| 955 | **28557** | **10** | 1 | **52** |
| 956 | **28558** | **10** | 1 | **52** |
| 957 | **28559** | **10** | 1 | **52** |
| 958 | **28560** | **10** | 1 | **52** |
| 959 | **28561** | **10** | 1 | **52** |
| 960 | **28562** | **10** | 1 | **52** |
| 961 | **28563** | **10** | 1 | **52** |
| 962 | **28564** | **10** | 1 | **52** |
| 963 | **28565** | **10** | 1 | **52** |
| 964 | **28566** | **9** | 2 | **52** |
| 965 | **28567** | **10** | 1 | **52** |
| 966 | **28568** | **9** | 2 | **52** |

| No. | MGAS number[1] | Number of Ts[2] | Allele number[3] | MLST type |
|---|---|---|---|---|
| 967 | **28569** | **10** | 1 | **52** |
| 968 | **28570** | **9** | 2 | **52** |
| 969 | **28571** | **9** | 2 | **52** |
| 970 | **28572** | **10** | 1 | **52** |
| 971 | **28573** | **10** | 1 | **52** |
| 972 | **28574** | **10** | 1 | **52** |
| 973 | **28575** | **ND** | - | **626** |
| 974 | **28576** | **10** | 1 | **52** |
| 975 | **28578** | **10** | 1 | **52** |
| 976 | **28579** | **10** | 1 | **52**[16] |
| 977 | **28580** | **10** | 1 | **52** |
| 978 | **28581** | **10** | 1 | **52** |
| 979 | **28582** | **10** | 1 | **52** |
| 980 | **28583** | **9** | 2 | **52** |
| 981 | **28584** | **9** | 2 | **52** |
| 982 | **28585** | **9** | 2 | **52**[17] |
| 983 | **28586** | **10** | 1 | **52** |
| 984 | **28587** | **10** | 1 | **52** |
| 985 | **28588** | **11** | 3 | **52** |
| 986 | **28589** | **10** | 1 | **52** |
| 987 | **28591** | **9** | 2 | **52** |
| 988 | **28592** | **9** | 2 | **52** |
| 989 | **28593** | **9** | 2 | **52** |
| 990 | **28594** | **9** | 2 | **52** |
| 991 | **28595** | **10** | 1 | **52** |
| 992 | **28596** | **10** | 1 | **52** |
| 993 | **28597** | **10** | 1 | **52** |
| 994 | **28598** | **10** | 1 | **52** |
| 995 | **28599** | **10** | 1 | **52** |
| 996 | **28600** | **10** | 1 | **52** |
| 997 | **28601** | **9** | 2 | **52** |
| 998 | **28602** | **10** | 1 | **52** |
| 999 | **28603** | **9** | 2 | **52** |
| 1000 | **28604** | **10** | 1 | **52** |
| 1001 | **28605** | **11** | 3 | **52** |
| 1002 | **28606** | **10** | 1 | **52** |
| 1003 | **28607** | **9** | 2 | **52** |
| 1004 | **28609** | **10** | 1 | **52** |
| 1005 | **28612** | **10** | 1 | **52** |
| 1006 | **28613** | **10** | 1 | **52** |
| 1007 | **28614** | **10** | 1 | **52** |
| 1008 | **28615** | **10** | 1 | **52** |
| 1009 | **28616** | **9** | 2 | **52** |
| 1010 | **28617** | **10** | 1 | **52** |

| No. | MGAS number(1) | Number of Ts(2) | Allele number(3) | MLST type |
|---|---|---|---|---|
| 1011 | **28618** | **11** | 3 | **52** |
| 1012 | **28619** | **9** | 2 | **52** |
| 1013 | **28620** | **10** | 1 | **52** |
| 1014 | **28621** | **10** | 1 | **52** |
| 1015 | **28622** | **10** | 1 | **52** |
| 1016 | **28623** | **8** | 4 | **52** |
| 1017 | **28625** | **9** | 2 | **52** |
| 1018 | **28626** | **11** | 3 | **52** |
| 1019 | **28627** | **9** | 2 | **52** |
| 1020 | **28628** | **ND** | - | **626** |
| 1021 | **28629** | **10** | 1 | **52** |
| 1022 | **28630** | **10** | 1 | **52** |
| 1023 | **28631** | **10** | 1 | **52** |
| 1024 | **28632** | **10** | 1 | **52**(7) |
| 1025 | **28633** | **10** | 1 | **52** |
| 1026 | **28634** | **10** | 1 | **52** |
| 1027 | **28635** | **9** | 2 | **52** |
| 1028 | **28636** | **10** | 1 | **52** |
| 1029 | **28637** | **10** | 1 | **52** |
| 1030 | **28638** | **9** | 2 | **52**(7) |
| 1031 | **28639** | **10** | 1 | **52** |
| 1032 | **28640** | **10** | 1 | **52** |
| 1033 | **28641** | **10** | 1 | **52**(16) |
| 1034 | **28642** | **10** | 1 | **52** |
| 1035 | **28643** | **10** | 1 | **52** |
| 1036 | **28644** | **10** | 1 | **52** |
| 1037 | **28645** | **11** | 3 | **458** |
| 1038 | **28646** | **10** | 1 | **52** |
| 1039 | **28647** | **10** | 1 | **52** |
| 1040 | **28648** | **8** | 23 | **52** |
| 1041 | **28649** | **9** | 2 | **52** |
| 1042 | **28650** | **10** | 1 | **52** |
| 1043 | **28651** | **10** | 1 | **52** |
| 1044 | **28652** | **10** | 1 | **52** |
| 1045 | **28653** | **10** | 1 | **52** |
| 1046 | **28654** | **9** | 2 | **52** |
| 1047 | **28655** | **10** | 1 | **52** |
| 1048 | **28656** | **10** | 1 | **52** |
| 1049 | **28657** | **11** | 3 | **52** |
| 1050 | **28658** | **9** | 2 | **52** |
| 1051 | **28659** | **10** | 1 | **52**(13) |
| 1052 | **28660** | **10** | 1 | **52** |
| 1053 | **28661** | **9** | 2 | **52** |
| 1054 | **28662** | **9** | 2 | **52** |

| No. | MGAS number[1] | Number of Ts[2] | Allele number[3] | MLST type |
|---|---|---|---|---|
| 1055 | **28664** | **9** | 2 | **52** |
| 1056 | **28665** | **9** | 2 | **52** |
| 1057 | **28666** | **9** | 2 | **52** |
| 1058 | **28667** | **9** | 2 | **52** |
| 1059 | **28668** | **10** | 1 | **52** |
| 1060 | **28669** | **9** | 2 | **52** |
| 1061 | **28670** | **10** | 1 | **52** |
| 1062 | **28671** | **9** | 2 | **52** |
| 1063 | **28673** | **10** | 1 | **52** |
| 1064 | **28674** | **10** | 1 | **52** |
| 1065 | **28675** | **9** | 2 | **52** |
| 1066 | **28677** | **10** | 1 | **52** |
| 1067 | **28678** | **9** | 2 | **52** |
| 1068 | **28679** | **9** | 2 | **52** |
| 1069 | **28680** | **9** | 2 | **52** |
| 1070 | **28681** | **9** | 2 | **52** |
| 1071 | **28682** | **10** | 1 | **52** |
| 1072 | **28683** | **10** | 1 | **52** |
| 1073 | **28684** | **11** | 3 | **52** |
| 1074 | **28685** | **10** | 1 | **52** |
| 1075 | **28686** | **9** | 2 | **52** |
| 1076 | **28687** | **10** | 1 | **52** |
| 1077 | **28688** | **10** | 1 | **52** |
| 1078 | **28689** | **9** | 2 | **52** |
| 1079 | **28690** | **10** | 1 | **52**[7] |
| 1080 | **28691** | **10** | 1 | **52** |
| 1081 | **28692** | **10** | 1 | **52** |
| 1082 | **28693** | **10** | 1 | **52** |
| 1083 | **28694** | **10** | 1 | **52** |
| 1084 | **28695** | **10** | 1 | **52** |
| 1085 | **28696** | **10** | 1 | **52** |
| 1086 | **28697** | **9** | 2 | **52**[7] |
| 1087 | **28698** | **10** | 1 | **52** |
| 1088 | **28699** | **10** | 1 | **52** |
| 1089 | **28700** | **11** | 3 | **52** |
| 1090 | **28701** | **9** | 2 | **52** |
| 1091 | **28702** | **9** | 2 | **52** |
| 1092 | **28703** | **9** | 2 | **52** |
| 1093 | **28704** | **10** | 1 | **52** |
| 1094 | **28705** | **9** | 6 | **52** |
| 1095 | **28706** | **10** | 1 | **52** |
| 1096 | **28707** | **10** | 1 | **458** |
| 1097 | **28708** | **9** | 2 | **52** |
| 1098 | **28709** | **9** | 2 | **52** |

| No. | MGAS number(1) | Number of Ts(2) | Allele number(3) | MLST type |
|---|---|---|---|---|
| 1099 | **28710** | **10** | 1 | **52** |
| 1100 | **28711** | **9** | 2 | **52** |
| 1101 | **28712** | **10** | 1 | **52** |
| 1102 | **28713** | **9** | 2 | **52** |
| 1103 | **28714** | **10** | 1 | **52** |
| 1104 | **28715** | **10** | 1 | **52**(7) |
| 1105 | **28716** | **9** | 2 | **52** |
| 1106 | **28717** | **11** | 3 | **52** |
| 1107 | **28718** | **10** | 1 | **52** |
| 1108 | **28719** | **10** | 1 | **52** |
| 1109 | **28720** | **10** | 1 | **52** |
| 1110 | **28721** | **10** | 1 | **52** |
| 1111 | **28722** | **11** | 3 | **52** |
| 1112 | **28723** | **10** | 1 | **52** |
| 1113 | **28724** | **10** | 1 | **52** |
| 1114 | **28725** | **ND** | - | **52** |
| 1115 | **28726** | **10** | 1 | **52** |
| 1116 | **28727** | **9** | 2 | **52** |
| 1117 | **28728** | **10** | 1 | **52** |
| 1118 | **28729** | **10** | 1 | **52**(13) |
| 1119 | **28730** | **9** | 2 | **52** |
| 1120 | **28731** | **9** | 2 | **52** |
| 1121 | **28732** | **9** | 2 | **52** |
| 1122 | **28733** | **10** | 1 | **52** |
| 1123 | **28734** | **10** | 1 | **52** |
| 1124 | **28735** | **9** | 2 | **52** |
| 1125 | **28736** | **10** | 1 | **52** |
| 1126 | **28737** | **9** | 2 | **52** |
| 1127 | **28738** | **9** | 2 | **52** |
| 1128 | **28739** | **9** | 2 | **52** |
| 1129 | **28740** | **9** | 2 | **52** |
| 1130 | **28741** | **10** | 1 | **52**(10) |
| 1131 | **28742** | **10** | 1 | **52** |
| 1132 | **28743** | **9** | 2 | **52** |
| 1133 | **28744** | **10** | 1 | **52** |
| 1134 | **28745** | **11** | 3 | **52** |
| 1135 | **28746** | **9** | 2 | **52** |
| 1136 | **28747** | **10** | 1 | **52** |
| 1137 | **28748** | **9** | 2 | **52**(7) |
| 1138 | **28749** | **10** | 1 | **52** |
| 1139 | **28750** | **9** | 2 | **52** |
| 1140 | **28751** | **10** | 1 | **52** |
| 1141 | **28752** | **10** | 1 | **52** |
| 1142 | **28753** | **10** | 1 | **52** |

| No. | MGAS number(1) | Number of Ts(2) | Allele number(3) | MLST type |
|---|---|---|---|---|
| 1143 | **28754** | **10** | 1 | **52** |
| 1144 | **28755** | **10** | 1 | **52** |
| 1145 | **28756** | **9** | 2 | **52** |
| 1146 | **28757** | **9** | 2 | **52** |
| 1147 | **28758** | **9** | 2 | **52** |
| 1148 | **28759** | **11** | 3 | **52** |
| 1149 | **28760** | **11** | 3 | **52(13)** |
| 1150 | **28761** | **10** | 1 | **52** |
| 1151 | **28762** | **10** | 1 | **52** |
| 1152 | **28763** | **10** | 1 | **52** |
| 1153 | **28764** | **10** | 1 | **52** |
| 1154 | **28765** | **10** | 1 | **52** |
| 1155 | **28766** | **9** | 2 | **458** |
| 1156 | **28767** | **10** | 1 | **52** |
| 1157 | **28768** | **9** | 2 | **52** |
| 1158 | **28769** | **10** | 1 | **52** |
| 1159 | **28770** | **9** | 2 | **52** |
| 1160 | **28771** | **9** | 2 | **52** |
| 1161 | **28772** | **10** | 1 | **52** |
| 1162 | **28773** | **10** | 17 | **52** |
| 1163 | **28774** | **10** | 1 | **52** |
| 1164 | **28775** | **9** | 2 | **52** |
| 1165 | **28776** | **10** | 1 | **52(13)** |
| 1166 | **28777** | **9** | 2 | **52(13)** |
| 1167 | **28778** | **9** | 2 | **52** |
| 1168 | **28779** | **10** | 1 | **52** |
| 1169 | **28780** | **10** | 1 | **52** |
| 1170 | **28781** | **10** | 1 | **52** |
| 1171 | **28782** | **11** | 3 | **52** |
| 1172 | **28783** | **10** | 1 | **52** |
| 1173 | **28784** | **10** | 1 | **52** |
| 1174 | **28785** | **10** | 1 | **52** |
| 1175 | **28786** | **10** | 1 | **52** |
| 1176 | **28787** | **10** | 1 | **52** |
| 1177 | **28788** | **9** | 2 | **52** |
| 1178 | **28789** | **ND** | - | **52** |
| 1179 | **28791** | **10** | 1 | **52** |
| 1180 | **28792** | **10** | 1 | **52** |
| 1181 | **28793** | **10** | 1 | **52** |
| 1182 | **28794** | **9** | 2 | **52** |
| 1183 | **28795** | **9** | 2 | **52** |
| 1184 | **28796** | **10** | 1 | **52** |
| 1185 | **28797** | **11** | 3 | **52** |
| 1186 | **28798** | **10** | 1 | **52** |

| No. | MGAS number(1) | Number of Ts(2) | Allele number(3) | MLST type |
|---|---|---|---|---|
| 1187 | **28799** | **10** | 1 | **52** |
| 1188 | **28801** | **9** | 2 | **52** |
| 1189 | **28802** | **9** | 2 | **52** |
| 1190 | **28803** | **10** | 1 | **52** |
| 1191 | **28804** | **10** | 1 | **52** |
| 1192 | **28805** | **10** | 1 | **52** |
| 1193 | **28806** | **10** | 1 | **52** |
| 1194 | **28807** | **10** | 1 | **52** |
| 1195 | **28808** | **9** | 2 | **52** |
| 1196 | **28809** | **10** | 1 | **52** |
| 1197 | **28810** | **10** | 1 | **52** |
| 1198 | **28811** | **9** | 2 | **52** |
| 1199 | **28812** | **9** | 2 | **52** |
| 1200 | **28813** | **10** | 1 | **52** |
| 1201 | **28814** | **10** | 1 | **52** |
| 1202 | **28815** | **10** | 1 | **52** |
| 1203 | **28816** | **10** | 1 | **52** |
| 1204 | **28817** | **10** | 1 | **52** |
| 1205 | **28818** | **11** | 3 | **52** |
| 1206 | **28819** | **10** | 1 | **52** |
| 1207 | **28820** | **10** | 1 | **52** |
| 1208 | **28821** | **10** | 1 | **52** |
| 1209 | **28822** | **10** | 1 | **52** |
| 1210 | **28823** | **10** | 1 | **52** |
| 1211 | **28824** | **9** | 2 | **52** |
| 1212 | **28825** | **10** | 1 | **52** |
| 1213 | **28826** | **9** | 2 | **52** |
| 1214 | **28828** | **10** | 1 | **52** |
| 1215 | **28829** | **9** | 2 | **52** |
| 1216 | **28830** | **10** | 1 | **52** |
| 1217 | **28831** | **10** | 1 | **52** |
| 1218 | **28832** | **9** | 2 | **52** |
| 1219 | **28833** | **10** | 1 | **52** |
| 1220 | **28834** | **10** | 1 | **52** |
| 1221 | **28835** | **10** | 1 | **52** |
| 1222 | **28836** | **9** | 2 | **52** |
| 1223 | **28837** | **9** | 2 | **52** |
| 1224 | **28838** | **10** | 1 | **52** |
| 1225 | **28839** | **10** | 1 | **52** |
| 1226 | **28840** | **10** | 1 | **52** |
| 1227 | **28841** | **9** | 2 | **52** |
| 1228 | **28842** | **10** | 1 | **52** |
| 1229 | **28843** | **9** | 2 | **52** |
| 1230 | **28844** | **10** | 1 | **52** |

| No. | MGAS number[1] | Number of Ts[2] | Allele number[3] | MLST type |
|---|---|---|---|---|
| 1231 | **28845** | **9** | 2 | **52** |
| 1232 | **28846** | **9** | 2 | **52** |
| 1233 | **28847** | **9** | 2 | **52** |
| 1234 | **28848** | **9** | 2 | **52** |
| 1235 | **28849** | **10** | 1 | **52** |
| 1236 | **28850** | **10** | 1 | **52** |
| 1237 | **28851** | **9** | 2 | **52** |
| 1238 | **28852** | **9** | 2 | **52** |
| 1239 | **28853** | **10** | 1 | **52** |
| 1240 | **28854** | **10** | 1 | **52** |
| 1241 | **28855** | **10** | 1 | **52** |
| 1242 | **28857** | **10** | 1 | **52** |
| 1243 | **28858** | **10** | 1 | **52** |
| 1244 | **28859** | **9** | 2 | **52** |
| 1245 | **28860** | **11** | 3 | **52** |
| 1246 | **28863** | **10** | 1 | **52** |
| 1247 | **28864** | **9** | 2 | **52** |
| 1248 | **28865** | **9** | 2 | **52** |
| 1249 | **28866** | **9** | 2 | **52** |
| 1250 | **28867** | **11** | 3 | **52** |
| 1251 | **28868** | **9** | 2 | **52** |
| 1252 | **28869** | **9** | 2 | **52** |
| 1253 | **28871** | **9** | 2 | **52** |
| 1254 | **28872** | **8** | 8 | **52** |
| 1255 | **28873** | **10** | 1 | **52** |
| 1256 | **28874** | **9** | 2 | **52** |
| 1257 | **28875** | **10** | 1 | **52** |
| 1258 | **28876** | **10** | 1 | **52** |
| 1259 | **28877** | **10** | 1 | **52** |
| 1260 | **28878** | **10** | 1 | **52** |
| 1261 | **28879** | **9** | 2 | **52** |
| 1262 | **28880** | **10** | 1 | **52** |
| 1263 | **28881** | **10** | 1 | **52** |
| 1264 | **28882** | **9** | 2 | **52** |
| 1265 | **28883** | **10** | 1 | **52** |
| 1266 | **28884** | **8** | 4 | **52** |
| 1267 | **28885** | **10** | 1 | **52** |
| 1268 | **28886** | **10** | 14 | **52** |
| 1269 | **28887** | **10** | 1 | **52**[10] |
| 1270 | **28888** | **9** | 2 | **52** |
| 1271 | **28889** | **10** | 1 | **52** |
| 1272 | **28890** | **10** | 1 | **52** |
| 1273 | **28891** | **10** | 1 | **52** |
| 1274 | **28892** | **10** | 1 | **52** |

| No. | MGAS number[1] | Number of Ts[2] | Allele number[3] | MLST type |
|---|---|---|---|---|
| 1275 | **28893** | **10** | 1 | **52** |
| 1276 | **28894** | **10** | 1 | **52** |
| 1277 | **28895** | **10** | 1 | **52** |
| 1278 | **28896** | **10** | 1 | **52** |
| 1279 | **28897** | **9** | 2 | **52** |
| 1280 | **28898** | **9** | 2 | **52** |
| 1281 | **28899** | **9** | 2 | **52** |
| 1282 | **28900** | **9** | 2 | **52** |
| 1283 | **28901** | **9** | 2 | **52** |
| 1284 | **28902** | **10** | 1 | **52** |
| 1285 | **28903** | **9** | 2 | **52**[7] |
| 1286 | **28904** | **10** | 1 | **52** |
| 1287 | **28905** | **9** | 2 | **52** |
| 1288 | **28906** | **9** | 2 | **52** |
| 1289 | **28907** | **10** | 1 | **52** |
| 1290 | **28908** | **9** | 2 | **52** |
| 1291 | **28909** | **9** | 2 | **52** |
| 1292 | **28910** | **11** | 3 | **52** |
| 1293 | **28911** | **9** | 2 | **52** |
| 1294 | **28912** | **10** | 1 | **52** |
| 1295 | **28913** | **10** | 1 | **52** |
| 1296 | **28914** | **9** | 2 | **52** |
| 1297 | **28915** | **10** | 1 | **52** |
| 1298 | **28916** | **9** | 2 | **52** |
| 1299 | **28917** | **10** | 1 | **845** |
| 1300 | **28918** | **11** | 3 | **52** |
| 1301 | **28919** | **9** | 2 | **52** |
| 1302 | **28920** | **10** | 1 | **52** |
| 1303 | **28921** | **10** | 1 | **52** |
| 1304 | **28922** | **10** | 1 | **52** |
| 1305 | **28923** | **10** | 1 | **52** |
| 1306 | **28924** | **10** | 1 | **456** |
| 1307 | **28925** | **11** | 3 | **52** |
| 1308 | **28926** | **9** | 2 | **52** |
| 1309 | **28927** | **10** | 1 | **52** |
| 1310 | **28928** | **11** | 3 | **850** |
| 1311 | **28929** | **10** | 1 | **52** |
| 1312 | **28930** | **10** | 1 | **52** |
| 1313 | **28931** | **9** | 2 | **52** |
| 1314 | **28932** | **10** | 1 | **52** |
| 1315 | **28933** | **11** | 3 | **52** |
| 1316 | **28934** | **10** | 1 | **52** |
| 1317 | **28935** | **10** | 1 | **52** |
| 1318 | **28936** | **9** | 2 | **52** |

| No. | MGAS number(1) | Number of Ts(2) | Allele number(3) | MLST type |
|---|---|---|---|---|
| 1319 | **28937** | **10** | 1 | **52** |
| 1320 | **28938** | **9** | 2 | **52** |
| 1321 | **28939** | **9** | 2 | **52** |
| 1322 | **28940** | **9** | 2 | **52** |
| 1323 | **28941** | **9** | 2 | **52** |
| 1324 | **28942** | **10** | 1 | **52** |
| 1325 | **28943** | **10** | 1 | **52** |
| 1326 | **28944** | **10** | 1 | **52** |
| 1327 | **28945** | **9** | 2 | **52** |
| 1328 | **28946** | **10** | 1 | **52** |
| 1329 | **28947** | **10** | 1 | **52** |
| 1330 | **28948** | **9** | 2 | **52** |
| 1331 | **28949** | **9** | 2 | **52** |
| 1332 | **28950** | **11** | 3 | **52** |
| 1333 | **28951** | **10** | 1 | **52** |
| 1334 | **28952** | **10** | 1 | **52** |
| 1335 | **28953** | **10** | 1 | **52** |
| 1336 | **29041** | **10** | 1 | **458** |
| 1337 | **29042** | **9** | 2 | **52** |
| 1338 | **29043** | **9** | 2 | **458** |
| 1339 | **29044** | **11** | 3 | **458** |
| 1340 | **29045** | **10** | 1 | **458** |
| 1341 | **29046** | **10** | 1 | **456** |
| 1342 | **29047** | **10** | 1 | **456** |
| 1343 | **29048** | **9** | 2 | **458** |
| 1344 | **29049** | **10** | 1 | **52** |
| 1345 | **29050** | **10** | 1 | **456** |
| 1346 | **29051** | **10** | 1 | **456** |
| 1347 | **29052** | **10** | 1 | **456** |
| 1348 | **29053** | **11** | 3 | **456** |
| 1349 | **29054** | **10** | 1 | **456** |
| 1350 | **29055** | **10** | 1 | **456** |
| 1351 | **29056** | **10** | 1 | **456** |
| 1352 | **29057** | **11** | 3 | **458** |
| 1353 | **29058** | **10** | 1 | **456** |
| 1354 | **29059** | **10** | 1 | **52** |
| 1355 | **29060** | **10** | 1 | **456** |
| 1356 | **29061** | **10** | 1 | **456** |
| 1357 | **29062** | **9** | 2 | **458** |
| 1358 | **29063** | **10** | 1 | **456** |
| 1359 | **29064** | **10** | 1 | **458** |
| 1360 | **29065** | **10** | 1 | **458** |
| 1361 | **29066** | **9** | 2 | **456** |
| 1362 | **29067** | **10** | 1 | **456** |

| No. | MGAS number(1) | Number of Ts(2) | Allele number(3) | MLST type |
|---|---|---|---|---|
| 1363 | **29068** | **10** | 1 | **456** |
| 1364 | **29069** | **10** | 1 | **52** |
| 1365 | **29070** | **10** | 1 | **456** |
| 1366 | **29071** | **9** | 2 | **456** |
| 1367 | **29072** | **9** | 2 | **458** |
| 1368 | **29073** | **11** | 3 | **456** |
| 1369 | **29074** | **9** | 2 | **52** |
| 1370 | **29075** | **10** | 1 | **458** |
| 1371 | **29076** | **9** | 2 | **456** |
| 1372 | **29077** | **10** | 1 | **458** |
| 1373 | **29078** | **9** | 2 | **52** |
| 1374 | **29079** | **10** | 1 | **52** |
| 1375 | **29080** | **9** | 2 | **458** |
| 1376 | **29081** | **10** | 1 | **456** |
| 1377 | **29082** | **10** | 1 | **456** |
| 1378 | **29083** | **9** | 2 | **458** |
| 1379 | **29084** | **10** | 1 | **456** |
| 1380 | **29085** | **10** | 12 | **52** |
| 1381 | **29086** | **10** | 1 | **52** |
| 1382 | **29087** | **10** | 1 | **456** |
| 1383 | **29088** | **9** | 2 | **456** |
| 1384 | **29089** | **11** | 3 | **458** |
| 1385 | **29090** | **10** | 1 | **456** |
| 1386 | **29091** | **10** | 1 | **456** |
| 1387 | **29092** | **10** | 1 | **52** |
| 1388 | **29093** | **9** | 2 | **456** |
| 1389 | **29094** | **10** | 1 | **52** |
| 1390 | **29095** | **10** | 1 | **458** |
| 1391 | **29096** | **10** | 1 | **456** |
| 1392 | **29097** | **9** | 2 | **458** |
| 1393 | **29098** | **9** | 2 | **52** |
| 1394 | **29099** | **9** | 2 | **52** |
| 1395 | **29100** | **10** | 1 | **456** |
| 1396 | **29101** | **10** | 1 | **458** |
| 1397 | **29102** | **10** | 1 | **52** |
| 1398 | **29103** | **9** | 2 | **456** |
| 1399 | **29104** | **10** | 1 | **456** |
| 1400 | **29105** | **10** | 1 | **456** |
| 1401 | **29106** | **10** | 1 | **456** |
| 1402 | **29107** | **9** | 2 | **456** |
| 1403 | **29108** | **7** | 22 | **458** |
| 1404 | **29109** | **9** | 2 | **456** |
| 1405 | **29110** | **10** | 1 | **458** |
| 1406 | **29111** | **10** | 1 | **52** |

| No. | MGAS number(1) | Number of Ts(2) | Allele number(3) | MLST type |
|---|---|---|---|---|
| 1407 | **29112** | **9** | 2 | **456** |
| 1408 | **29114** | **11** | 3 | **456** |
| 1409 | **29115** | **10** | 1 | **456** |
| 1410 | **29116** | **10** | 1 | **52** |
| 1411 | **29117** | **9** | 2 | **456** |
| 1412 | **29118** | **9** | 2 | **458** |
| 1413 | **29119** | **10** | 1 | **456** |
| 1414 | **29120** | **9** | 2 | **458** |
| 1415 | **29121** | **10** | 1 | **456** |
| 1416 | **29122** | **10** | 1 | **52** |
| 1417 | **29123** | **10** | 1 | **52** |
| 1418 | **29124** | **10** | 1 | **458** |
| 1419 | **29125** | **10** | 1 | **456** |
| 1420 | **29126** | **10** | 1 | **458** |
| 1421 | **29127** | **9** | 2 | **456** |
| 1422 | **29128** | **10** | 1 | **456** |
| 1423 | **29129** | **10** | 1 | **456** |
| 1424 | **29130** | **10** | 1 | **458** |
| 1425 | **29131** | **12** | 5 | **456** |
| 1426 | **29133** | **10** | 1 | **52** |
| 1427 | **29135** | **9** | 2 | **456** |
| 1428 | **29136** | **9** | 2 | **458** |
| 1429 | **29137** | **9** | 2 | **458** |
| 1430 | **29138** | **9** | 2 | **456** |
| 1431 | **29139** | **10** | 1 | **458** |
| 1432 | **29140** | **9** | 2 | **456** |
| 1433 | **29141** | **10** | 1 | **456** |
| 1434 | **29142** | **9** | 2 | **52** |
| 1435 | **29143** | **9** | 2 | **52** |
| 1436 | **29144** | **9** | 2 | **456** |
| 1437 | **29145** | **10** | 1 | **458** |
| 1438 | **29146** | **10** | 1 | **458** |
| 1439 | **29147** | **10** | 1 | **52** |
| 1440 | **29148** | **9** | 2 | **458** |
| 1441 | **29149** | **10** | 1 | **458** |
| 1442 | **29150** | **10** | 1 | **52** |
| 1443 | **29151** | **10** | 1 | **458** |
| 1444 | **29152** | **10** | 1 | **456**(7) |
| 1445 | **29153** | **10** | 1 | **456** |
| 1446 | **29154** | **10** | 1 | **52** |
| 1447 | **29155** | **10** | 1 | **52** |
| 1448 | **29156** | **10** | 1 | **52** |
| 1449 | **29157** | **10** | 1 | **456** |
| 1450 | **29158** | **10** | 1 | **52** |

| No. | MGAS number[(1)] | Number of Ts[(2)] | Allele number[(3)] | MLST type |
|---|---|---|---|---|
| 1451 | **29159** | **10** | 1 | **458** |
| 1452 | **29160** | **9** | 2 | **458** |
| 1453 | **29161** | **10** | 1 | **52** |
| 1454 | **29162** | **10** | 1 | **52** |
| 1455 | **29163** | **10** | 1 | **456** |
| 1456 | **29164** | **10** | 1 | **456** |
| 1457 | **29166** | **10** | 1 | **52** |
| 1458 | **29167** | **10** | 1 | **456** |
| 1459 | **29168** | **11** | 3 | **458** |
| 1460 | **29169** | **10** | 1 | **456** |
| 1461 | **29170** | **9** | 2 | **458** |
| 1462 | **29171** | **10** | 1 | **52** |
| 1463 | **29172** | **10** | 1 | **456** |
| 1464 | **29173** | **10** | 1 | **456** |
| 1465 | **29174** | **9** | 2 | **456** |
| 1466 | **29175** | **10** | 1 | **458** |
| 1467 | **29176** | **10** | 1 | **456** |
| 1468 | **29177** | **9** | 2 | **52** |
| 1469 | **29178** | **10** | 1 | **456** |
| 1470 | **29179** | **10** | 1 | **52** |
| 1471 | **29180** | **9** | 2 | **52** |
| 1472 | **29181** | **10** | 1 | **456** |
| 1473 | **29182** | **10** | 1 | **456** |
| 1474 | **29183** | **10** | 1 | **456** |
| 1475 | **29184** | **10** | 1 | **458** |
| 1476 | **29185** | **10** | 1 | **458** |
| 1477 | **29186** | **9** | 2 | **52** |
| 1478 | **29187** | **10** | 1 | **458** |
| 1479 | **29188** | **9** | 2 | **456** |
| 1480 | **29189** | **10** | 1 | **52** |
| 1481 | **29190** | **10** | 1 | **52** |
| 1482 | **29191** | **9** | 2 | **458** |
| 1483 | **29192** | **9** | 2 | **458** |
| 1484 | **29193** | **9** | 2 | **458** |
| 1485 | **29194** | **10** | 1 | **456** |
| 1486 | **29195** | **10** | 1 | **456** |
| 1487 | **29196** | **9** | 2 | **458** |
| 1488 | **29197** | **10** | 1 | **458** |
| 1489 | **29198** | **10** | 1 | **456** |
| 1490 | **29199** | **10** | 1 | **458** |
| 1491 | **29200** | **9** | 2 | **52** |
| 1492 | **29201** | **10** | 1 | **456** |
| 1493 | **29202** | **9** | 2 | **52** |
| 1494 | **29203** | **10** | 1 | **458** |

| No. | MGAS number(1) | Number of Ts(2) | Allele number(3) | MLST type |
|---|---|---|---|---|
| 1495 | **29204** | **10** | 1 | **52** |
| 1496 | **29205** | **9** | 2 | **456** |
| 1497 | **29208** | **10** | 1 | **458** |
| 1498 | **29209** | **11** | 3 | **456** |
| 1499 | **29210** | **10** | 1 | **52** |
| 1500 | **29211** | **9** | 2 | **456** |
| 1501 | **29212** | **8** | 8 | **52** |
| 1502 | **29213** | **10** | 1 | **52** |
| 1503 | **29214** | **9** | 2 | **456** |
| 1504 | **29215** | **10** | 1 | **456** |
| 1505 | **29216** | **10** | 1 | **52** |
| 1506 | **29217** | **10** | 1 | **458** |
| 1507 | **29218** | **9** | 2 | **456** |
| 1508 | **29219** | **10** | 1 | **458** |
| 1509 | **29220** | **10** | 1 | **456** |
| 1510 | **29221** | **9** | 2 | **52** |
| 1511 | **29222** | **10** | 1 | **458** |
| 1512 | **29223** | **10** | 1 | **456** |
| 1513 | **29224** | **9** | 2 | **52** |
| 1514 | **29225** | **10** | 1 | **52** |
| 1515 | **29226** | **10** | 1 | **456** |
| 1516 | **29228** | **11** | 3 | **458** |
| 1517 | **29229** | **10** | 1 | **456** |
| 1518 | **29230** | **10** | 1 | **458** |
| 1519 | **29231** | **10** | 1 | **52** |
| 1520 | **29232** | **9** | 2 | **458** |
| 1521 | **29233** | **9** | 2 | **52** |
| 1522 | **29234** | **9** | 2 | **456** |
| 1523 | **29235** | **10** | 1 | **456** |
| 1524 | **29236** | **9** | 2 | **456** |
| 1525 | **29237** | **10** | 1 | **458** |
| 1526 | **29238** | **10** | 1 | **456** |
| 1527 | **29239** | **10** | 1 | **52** |
| 1528 | **29240** | **9** | 2 | **456** |
| 1529 | **29241** | **10** | 1 | **52** |
| 1530 | **29242** | **9** | 2 | **52** |
| 1531 | **29243** | **10** | 1 | **458** |
| 1532 | **29244** | **9** | 2 | **456** |
| 1533 | **29245** | **9** | 2 | **456** |
| 1534 | **29246** | **9** | 2 | **456** |
| 1535 | **29247** | **9** | 2 | **458** |
| 1536 | **29248** | **10** | 1 | **52** |
| 1537 | **29249** | **10** | 1 | **52** |
| 1538 | **29250** | **9** | 2 | **52** |

| No. | MGAS number(1) | Number of Ts(2) | Allele number(3) | MLST type |
|---|---|---|---|---|
| 1539 | **29251** | **10** | 1 | **458** |
| 1540 | **29253** | **9** | 2 | **456** |
| 1541 | **29254** | **10** | 1 | **456** |
| 1542 | **29255** | **11** | 3 | **456** |
| 1543 | **29256** | **10** | 1 | **456** |
| 1544 | **29257** | **9** | 2 | **456** |
| 1545 | **29258** | **9** | 2 | **456** |
| 1546 | **29259** | **10** | 1 | **458** |
| 1547 | **29260** | **10** | 1 | **458** |
| 1548 | **29261** | **10** | 1 | **458** |
| 1549 | **29262** | **11** | 3 | **458** |
| 1550 | **29263** | **10** | 1 | **458** |
| 1551 | **29264** | **10** | 1 | **458** |
| 1552 | **29265** | **10** | 1 | **456** |
| 1553 | **29266** | **11** | 3 | **456** |
| 1554 | **29267** | **10** | 1 | **458** |
| 1555 | **29268** | **10** | 1 | **456** |
| 1556 | **29269** | **10** | 1 | **52** |
| 1557 | **29270** | **10** | 1 | **52** |
| 1558 | **29271** | **10** | 1 | **52** |
| 1559 | **29272** | **10** | 1 | **52** |
| 1560 | **29273** | **10** | 1 | **52** |
| 1561 | **29274** | **11** | 3 | **456** |
| 1562 | **29275** | **9** | 2 | **458** |
| 1563 | **29276** | **10** | 1 | **456** |
| 1564 | **29277** | **10** | 1 | **456** |
| 1565 | **29278** | **10** | 1 | **458** |
| 1566 | **29279** | **10** | 1 | **456** |
| 1567 | **29280** | **10** | 1 | **52** |
| 1568 | **29281** | **10** | 1 | **458** |
| 1569 | **29282** | **8** | 4 | **52** |
| 1570 | **29283** | **10** | 1 | **458** |
| 1571 | **29284** | **10** | 1 | **458** |
| 1572 | **29285** | **9** | 2 | **456** |
| 1573 | **29286** | **10** | 1 | **52** |
| 1574 | **29287** | **10** | 1 | **456** |
| 1575 | **29289** | **11** | 3 | **456** |
| 1576 | **29290** | **9** | 2 | **456**(7) |
| 1577 | **29291** | **9** | 2 | **52** |
| 1578 | **29292** | **10** | 1 | **52** |
| 1579 | **29293** | **10** | 1 | **52** |
| 1580 | **29294** | **9** | 2 | **52** |
| 1581 | **29295** | **10** | 1 | **458** |
| 1582 | **29296** | **10** | 1 | **456** |

| No. | MGAS number[1] | Number of Ts[2] | Allele number[3] | MLST type |
|---|---|---|---|---|
| 1583 | **29297** | **9** | 2 | **52** |
| 1584 | **29298** | **10** | 1 | **456** |
| 1585 | **29299** | **10** | 1 | **52** |
| 1586 | **29300** | **9** | 2 | **456** |
| 1587 | **29301** | **9** | 2 | **456** |
| 1588 | **29302** | **9** | 2 | **458** |
| 1589 | **29303** | **10** | 1 | **52** |
| 1590 | **29304** | **10** | 1 | **456** |
| 1591 | **29305** | **10** | 1 | **52** |
| 1592 | **29306** | **10** | 1 | **52** |
| 1593 | **29307** | **8** | 4 | **456** |
| 1594 | **29308** | **9** | 2 | **458** |
| 1595 | **29310** | **10** | 1 | **458** |
| 1596 | **29311** | **10** | 1 | **456** |
| 1597 | **29312** | **9** | 2 | **456** |
| 1598 | **29313** | **9** | 2 | **52** |
| 1599 | **29314** | **10** | 1 | **458** |
| 1600 | **29315** | **9** | 2 | **456** |
| 1601 | **29316** | **10** | 1 | **456?** |
| 1602 | **29317** | **9** | 2 | **458** |
| 1603 | **29318** | **11** | 3 | **456** |
| 1604 | **29319** | **11** | 3 | **456** |
| 1605 | **29320** | **10** | 1 | **456** |
| 1606 | **29321** | **10** | 1 | **456** |
| 1607 | **29322** | **12** | 5 | **456** |
| 1608 | **29323** | **10** | 1 | **456** |
| 1609 | **29324** | **9** | 2 | **456** |
| 1610 | **29325** | **9** | 2 | **456** |
| 1611 | **29326** | **9** | 2 | **52** |
| 1612 | **29327** | **10** | 1 | **456** |
| 1613 | **29328** | **10** | 1 | **456** |
| 1614 | **29329** | **10** | 1 | **52** |
| 1615 | **29330** | **10** | 1 | **456** |
| 1616 | **29331** | **10** | 1 | **52** |
| 1617 | **29332** | **9** | 2 | **458** |
| 1618 | **29333** | **9** | 2 | **456** |
| 1619 | **29334** | **9** | 2 | **456** |
| 1620 | **29335** | **10** | 1 | **458** |
| 1621 | **29336** | **9** | 2 | **52** |
| 1622 | **29337** | **10** | 1 | **456** |
| 1623 | **29338** | **8** | 4 | **456** |
| 1624 | **29339** | **8** | 4 | **456** |
| 1625 | **29340** | **11** | 3 | **52** |
| 1626 | **29341** | **9** | 2 | **52** |

| No. | MGAS number(1) | Number of Ts(2) | Allele number(3) | MLST type |
|---|---|---|---|---|
| 1627 | **29342** | **8** | 4 | **456** |
| 1628 | **29343** | **8** | 4 | **456** |
| 1629 | **29344** | **9** | 2 | **458** |
| 1630 | **29345** | **9** | 2 | **456** |
| 1631 | **29346** | **9** | 2 | **456** |
| 1632 | **29347** | **11** | 3 | **456** |
| 1633 | **29348** | **9** | 2 | **456** |
| 1634 | **29349** | **10** | 1 | **52** |
| 1635 | **29350** | **10** | 1 | **456** |
| 1636 | **29351** | **9** | 6 | **456** |
| 1637 | **29352** | **10** | 1 | **456** |
| 1638 | **29353** | **9** | 2 | **456** |
| 1639 | **29354** | **10** | 1 | **52** |
| 1640 | **29355** | **10** | 1 | **458** |
| 1641 | **29356** | **10** | 1 | **456** |
| 1642 | **29357** | **11** | 3 | **458** |
| 1643 | **29358** | **10** | 1 | **458** |
| 1644 | **29359** | **10** | 1 | **52** |
| 1645 | **29360** | **10** | 1 | **52** |
| 1646 | **29361** | **10** | 1 | **52** |
| 1647 | **29362** | **9** | 2 | **458** |
| 1648 | **29363** | **10** | 1 | **456** |
| 1649 | **29364** | **9** | 2 | **458** |
| 1650 | **29365** | **10** | 1 | **52** |
| 1651 | **29366** | **10** | 1 | **458** |
| 1652 | **29367** | **10** | 1 | **456** |
| 1653 | **29368** | **10** | 1 | **456** |
| 1654 | **29369** | **10** | 1 | **458** |
| 1655 | **29370** | **10** | 1 | **458** |
| 1656 | **29371** | **10** | 1 | **52** |
| 1657 | **29372** | **9** | 2 | **458** |
| 1658 | **29373** | **10** | 1 | **52** |
| 1659 | **29374** | **9** | 2 | **52** |
| 1660 | **29375** | **9** | 2 | **458** |
| 1661 | **29376** | **9** | 2 | **52** |
| 1662 | **29377** | **9** | 2 | **456** |
| 1663 | **29378** | **10** | 1 | **456** |
| 1664 | **29379** | **10** | 1 | **456** |
| 1665 | **29380** | **9** | 2 | **52** |
| 1666 | **29381** | **9** | 2 | **52** |
| 1667 | **29382** | **9** | 2 | **458** |
| 1668 | **29383** | **10** | 1 | **52** |
| 1669 | **29384** | **9** | 2 | **456** |
| 1670 | **29385** | **9** | 2 | **458** |

| No. | MGAS number(1) | Number of Ts(2) | Allele number(3) | MLST type |
|---|---|---|---|---|
| 1671 | **29386** | **8** | 4 | **456** |
| 1672 | **29387** | **10** | 1 | **458** |
| 1673 | **29388** | **9** | 2 | **456** |
| 1674 | **29389** | **10** | 1 | **456** |
| 1675 | **29390** | **10** | 1 | **52** |
| 1676 | **29391** | **9** | 2 | **456** |
| 1677 | **29392** | **10** | 1 | **456** |
| 1678 | **29393** | **9** | 2 | **52** |
| 1679 | **29394** | **10** | 1 | **52** |
| 1680 | **29395** | **10** | 1 | **52** |
| 1681 | **29396** | **10** | 1 | **458** |
| 1682 | **29397** | **10** | 1 | **458** |
| 1683 | **29398** | **10** | 1 | **456** |
| 1684 | **29399** | **9** | 2 | **456** |
| 1685 | **29400** | **10** | 1 | **52** |
| 1686 | **29402** | **10** | 1 | **456** |
| 1687 | **29403** | **10** | 1 | **456** |
| 1688 | **29404** | **9** | 2 | **456** |
| 1689 | **29406** | **10** | 1 | **458** |
| 1690 | **29407** | **10** | 1 | **52** |
| 1691 | **29408** | **10** | 1 | **52** |
| 1692 | **29409** | **10** | 1 | **458** |
| 1693 | **29410** | **10** | 1 | **52** |
| 1694 | **29411** | **10** | 1 | **52** |
| 1695 | **29412** | **9** | 2 | **456** |
| 1696 | **29413** | **10** | 1 | **456** |
| 1697 | **29414** | **10** | 1 | **456** |
| 1698 | **29415** | **10** | 1 | **52** |
| 1699 | **29416** | **10** | 1 | **52** |
| 1700 | **29417** | **10** | 1 | **456** |
| 1701 | **29418** | **10** | 1 | **458** |
| 1702 | **29419** | **10** | 1 | **456** |
| 1703 | **29420** | **9** | 2 | **458** |
| 1704 | **29421** | **10** | 1 | **458** |
| 1705 | **29422** | **10** | 1 | **456** |
| 1706 | **29423** | **10** | 1 | **456** |
| 1707 | **29424** | **10** | 1 | **456** |
| 1708 | **29425** | **11** | 3 | **456** |
| 1709 | **29426** | **10** | 1 | **456** |
| 1710 | **29427** | **9** | 2 | **52** |
| 1711 | **29428** | **9** | 2 | **52** |
| 1712 | **29429** | **10** | 1 | **52** |
| 1713 | **29430** | **10** | 1 | **458** |
| 1714 | **29431** | **10** | 1 | **52** |

| No. | MGAS number[(1)] | Number of Ts[(2)] | Allele number[(3)] | MLST type |
|---|---|---|---|---|
| 1715 | **29432** | **10** | 1 | **456** |
| 1716 | **29433** | **10** | 1 | **458** |
| 1717 | **29434** | **10** | 1 | **458** |
| 1718 | **29435** | **10** | 1 | **52** |
| 1719 | **29436** | **10** | 1 | **458** |
| 1720 | **29439** | **10** | 1 | **456** |
| 1721 | **29440** | **10** | 1 | **456** |
| 1722 | **29441** | **9** | 2 | **52** |
| 1723 | **29442** | **10** | 1 | **456** |
| 1724 | **29443** | **9** | 2 | **456** |
| 1725 | **29444** | **10** | 1 | **52** |
| 1726 | **29445** | **10** | 1 | **52** |
| 1727 | **29446** | **10** | 1 | **52** |
| 1728 | **29447** | **13** | 29 | **52** |
| 1729 | **29448** | **11** | 3 | **456** |
| 1730 | **29450** | **9** | 2 | **52** |
| 1731 | **29451** | **10** | 1 | **456** |
| 1732 | **29452** | **10** | 1 | **52** |
| 1733 | **29453** | **10** | 1 | **456** |
| 1734 | **29454** | **10** | 1 | **456** |
| 1735 | **29455** | **9** | 2 | **52** |
| 1736 | **29456** | **10** | 1 | **456** |
| 1737 | **29457** | **10** | 1 | **456** |
| 1738 | **29458** | **9** | 2 | **456** |
| 1739 | **29459** | **10** | 1 | **52** |
| 1740 | **29460** | **10** | 1 | **456** |
| 1741 | **29461** | **10** | 1 | **456** |
| 1742 | **29462** | **9** | 2 | **456** |
| 1743 | **29463** | **9** | 2 | **52** |
| 1744 | **29464** | **9** | 2 | **52** |
| 1745 | **29465** | **9** | 2 | **458** |
| 1746 | **29466** | **10** | 1 | **456** |
| 1747 | **29467** | **10** | 1 | **52** |
| 1748 | **29468** | **9** | 2 | **456** |
| 1749 | **29469** | **9** | 2 | **52** |
| 1750 | **29470** | **10** | 1 | **456** |
| 1751 | **29471** | **10** | 1 | **456** |
| 1752 | **29472** | **10** | 1 | **52** |
| 1753 | **29473** | **10** | 1 | **52** |
| 1754 | **29474** | **9** | 2 | **456** |
| 1755 | **29475** | **9** | 2 | **52** |
| 1756 | **29476** | **10** | 1 | **456** |
| 1757 | **29477** | **10** | 1 | **52** |
| 1758 | **29478** | **9** | 2 | **456** |

| No. | MGAS number(1) | Number of Ts(2) | Allele number(3) | MLST type |
|---|---|---|---|---|
| 1759 | **29479** | **9** | 2 | **52(5)** |
| 1760 | **29480** | **10** | 1 | **456** |
| 1761 | **29481** | **10** | 1 | **456** |
| 1762 | **29483** | **10** | 1 | **456** |
| 1763 | **29484** | **11** | 3 | **52** |
| 1764 | **29485** | **10** | 1 | **52** |
| 1765 | **29486** | **10** | 1 | **456** |
| 1766 | **29487** | **9** | 2 | **456** |
| 1767 | **29488** | **10** | 1 | **52** |
| 1768 | **29489** | **9** | 2 | **52** |
| 1769 | **29490** | **10** | 1 | **52** |
| 1770 | **29491** | **11** | 3 | **456** |
| 1771 | **29492** | **10** | 1 | **456** |
| 1772 | **29493** | **9** | 2 | **456** |
| 1773 | **29494** | **9** | 2 | **52** |
| 1774 | **29495** | **10** | 1 | **456** |
| 1775 | **29496** | **9** | 2 | **52** |
| 1776 | **29497** | **9** | 2 | **52** |
| 1777 | **29498** | **9** | 2 | **52** |
| 1778 | **29500** | **10** | 1 | **458** |
| 1779 | **29501** | **10** | 1 | **456** |
| 1780 | **29502** | **9** | 2 | **456** |
| 1781 | **29503** | **10** | 1 | **456** |
| 1782 | **29504** | **10** | 1 | **52** |
| 1783 | **29505** | **9** | 2 | **52** |
| 1784 | **29506** | **10** | 1 | **52** |
| 1785 | **29507** | **10** | 1 | **52** |
| 1786 | **29508** | **11** | 3 | **456** |
| 1787 | **29509** | **10** | 1 | **52** |
| 1788 | **29510** | **9** | 2 | **456** |
| 1789 | **29511** | **9** | 2 | **456** |
| 1790 | **29512** | **11** | 3 | **52** |
| 1791 | **29513** | **10** | 1 | **456** |
| 1792 | **29514** | **11** | 3 | **458** |
| 1793 | **29515** | **9** | 2 | **456** |
| 1794 | **29516** | **10** | 1 | **52** |
| 1795 | **29517** | **10** | 1 | **456** |
| 1796 | **29518** | **10** | 1 | **52** |
| 1797 | **29519** | **9** | 2 | **52** |
| 1798 | **29520** | **10** | 1 | **456** |
| 1799 | **29521** | **9** | 2 | **456** |
| 1800 | **29522** | **9** | 2 | **52** |
| 1801 | **29523** | **10** | 1 | **52** |
| 1802 | **29524** | **9** | 2 | **52** |

| No. | MGAS number(1) | Number of Ts(2) | Allele number(3) | MLST type |
|---|---|---|---|---|
| 1803 | **29525** | **9** | 2 | **456** |
| 1804 | **29526** | **11** | 3 | **456** |
| 1805 | **29527** | **10** | 1 | **456** |
| 1806 | **29528** | **10** | 1 | **456** |
| 1807 | **29529** | **10** | 1 | **52** |
| 1808 | **29530** | **10** | 1 | **456** |
| 1809 | **29531** | **10** | 1 | **456** |
| 1810 | **29532** | **10** | 1 | **52** |
| 1811 | **29533** | **10** | 1 | **52** |
| 1812 | **29534** | **9** | 2 | **52** |
| 1813 | **29536** | **10** | 1 | **456** |
| 1814 | **29537** | **10** | 1 | **456** |
| 1815 | **29538** | **10** | 1 | **52** |
| 1816 | **29539** | **10** | 13 | **456** |
| 1817 | **29540** | **9** | 2 | **52** |
| 1818 | **29541** | **10** | 1 | **456** |
| 1819 | **29542** | **9** | 2 | **52** |
| 1820 | **29543** | **10** | 1 | **52** |
| 1821 | **29544** | **10** | 1 | **52** |
| 1822 | **29545** | **10** | 1 | **456** |
| 1823 | **29546** | **10** | 1 | **456** |
| 1824 | **29547** | **10** | 1 | **52** |
| 1825 | **29548** | **9** | 2 | **52** |
| 1826 | **29549** | **10** | 1 | **52** |
| 1827 | **29550** | **10** | 1 | **52** |
| 1828 | **29551** | **9** | 2 | **52** |
| 1829 | **29552** | **9** | 2 | **52** |
| 1830 | **29553** | **10** | 1 | **52** |
| 1831 | **29554** | **10** | 1 | **52** |
| 1832 | **29555** | **11** | 3 | **456** |
| 1833 | **29556** | **10** | 1 | **52** |
| 1834 | **29557** | **9** | 2 | **456** |
| 1835 | **29558** | **10** | 1 | **456** |
| 1836 | **29559** | **9** | 2 | **456** |
| 1837 | **29560** | **10** | 1 | **456** |
| 1838 | **29561** | **9** | 2 | **52** |
| 1839 | **29562** | **9** | 2 | **456** |
| 1840 | **29563** | **9** | 2 | **52** |
| 1841 | **29564** | **9** | 2 | **52** |
| 1842 | **29565** | **9** | 2 | **456** |
| 1843 | **29566** | **8** | 4 | **52** |
| 1844 | **29567** | **9** | 2 | **456** |
| 1845 | **29568** | **11** | 3 | **52** |
| 1846 | **29569** | **9** | 2 | **52** |

| No. | MGAS number(1) | Number of Ts(2) | Allele number(3) | MLST type |
|---|---|---|---|---|
| 1847 | **29570** | **9** | 2 | **456** |
| 1848 | **29571** | **10** | 1 | **456** |
| 1849 | **29572** | **9** | 2 | **456** |
| 1850 | **29573** | **9** | 2 | **456** |
| 1851 | **29575** | **10** | 1 | **52** |
| 1852 | **29576** | **10** | 1 | **52** |
| 1853 | **29577** | **9** | 2 | **456** |
| 1854 | **29578** | **10** | 1 | **456** |
| 1855 | **29579** | **11** | 3 | **52** |
| 1856 | **29580** | **10** | 1 | **456** |
| 1857 | **29581** | **9** | 2 | **456** |
| 1858 | **29582** | **10** | 1 | **52** |
| 1859 | **29583** | **10** | 1 | **456** |
| 1860 | **29584** | **9** | 2 | **52** |
| 1861 | **29585** | **9** | 2 | **456** |
| 1862 | **29586** | **10** | 1 | **456** |
| 1863 | **29587** | **10** | 1 | **52** |
| 1864 | **29588** | **10** | 1 | **456** |
| 1865 | **29589** | **9** | 2 | **52** |
| 1866 | **29590** | **9** | 2 | **52** |
| 1867 | **29591** | **9** | 2 | **456** |
| 1868 | **29592** | **9** | 2 | **456** |
| 1869 | **29593** | **10** | 1 | **52** |
| 1870 | **29594** | **11** | 3 | **456** |
| 1871 | **29596** | **10** | 1 | **456** |
| 1872 | **29597** | **10** | 1 | **456** |
| 1873 | **29598** | **9** | 2 | **52** |
| 1874 | **29599** | **10** | 1 | **456** |
| 1875 | **29600** | **10** | 1 | **52** |
| 1876 | **29601** | **10** | 1 | **52** |
| 1877 | **29602** | **10** | 1 | **456** |
| 1878 | **29603** | **10** | 1 | **456** |
| 1879 | **29604** | **10** | 1 | **456** |
| 1880 | **29605** | **10** | 1 | **52** |
| 1881 | **29606** | **9** | 2 | **456** |
| 1882 | **29607** | **11** | 3 | **52** |
| 1883 | **29608** | **9** | 2 | **52** |
| 1884 | **29609** | **10** | 1 | **52** |
| 1885 | **29610** | **10** | 1 | **52** |
| 1886 | **29611** | **11** | 3 | **52** |
| 1887 | **29612** | **10** | 1 | **52** |
| 1888 | **29613** | **10** | 1 | **456** |
| 1889 | **29614** | **9** | 2 | **52** |
| 1890 | **29615** | **9** | 2 | **456** |

| No. | MGAS number(1) | Number of Ts(2) | Allele number(3) | MLST type |
|---|---|---|---|---|
| 1891 | **29616** | **9** | 2 | **456** |
| 1892 | **29617** | **10** | 1 | **52** |
| 1893 | **29618** | **10** | 1 | **52** |
| 1894 | **29619** | **10** | 1 | **52** |
| 1895 | **29620** | **10** | 1 | **456** |
| 1896 | **29621** | **9** | 2 | **456** |
| 1897 | **29622** | **9** | 2 | **456** |
| 1898 | **29623** | **10** | 1 | **456** |
| 1899 | **29625** | **9** | 2 | **52** |
| 1900 | **29626** | **9** | 2 | **52** |
| 1901 | **29627** | **10** | 1 | **52** |
| 1902 | **29628** | **11** | 3 | **456** |
| 1903 | **29629** | **10** | 1 | **52** |
| 1904 | **29630** | **9** | 2 | **456** |
| 1905 | **29631** | **10** | 1 | **456** |
| 1906 | **29632** | **10** | 1 | **52** |
| 1907 | **29633** | **10** | 1 | **52** |
| 1908 | **29634** | **10** | 1 | **52** |
| 1909 | **29635** | **10** | 1 | **52** |
| 1910 | **29636** | **10** | 1 | **456** |
| 1911 | **30066** | **10** | 1 | **458** |
| 1912 | **30067** | **9** | 2 | **456** |
| 1913 | **30068** | **10** | 1 | **456** |
| 1914 | **30069** | **10** | 1 | **456** |
| 1915 | **30070** | **10** | 1 | **458** |
| 1916 | **30071** | **9** | 2 | **52** |
| 1917 | **30072** | **10** | 1 | **456** |
| 1918 | **30073** | **9** | 2 | **458** |
| 1919 | **30074** | **10** | 1 | **458** |
| 1920 | **30076** | **10** | 1 | **458** |
| 1921 | **30077** | **10** | 1 | **458** |
| 1922 | **30078** | **9** | 2 | **456** |
| 1923 | **30080** | **9** | 2 | **456** |
| 1924 | **30081** | **10** | 1 | **458** |
| 1925 | **30084** | **9** | 2 | **458** |
| 1926 | **31876** | **10** | 1 | **52** |
| 1927 | **31877** | **10** | 1 | **52** |
| 1928 | **31878** | **10** | 1 | **52** |
| 1929 | **31879** | **9** | 2 | **52** |
| 1930 | **31880** | **10** | 1 | **456** |
| 1931 | **31881** | **10** | 1 | **52** |
| 1932 | **31882** | **8** | 4 | **52** |
| 1933 | **31883** | **10** | 1 | **456** |
| 1934 | **31884** | **10** | 1 | **52** |

| No. | MGAS number(1) | Number of Ts(2) | Allele number(3) | MLST type |
|---|---|---|---|---|
| 1935 | **31885** | **10** | 1 | **52** |
| 1936 | **31886** | **10** | 1 | **52** |
| 1937 | **31887** | **9** | 2 | **456** |
| 1938 | **31888** | **10** | 1 | **456** |
| 1939 | **31889** | **10** | 1 | **52** |
| 1940 | **31890** | **10** | 1 | **52** |
| 1941 | **31891** | **10** | 1 | **52** |
| 1942 | **31892** | **10** | 1 | **52** |
| 1943 | **31893** | **9** | 2 | **456** |
| 1944 | **31894** | **10** | 1 | **52** |
| 1945 | **31895** | **10** | 1 | **52** |
| 1946 | **31896** | **10** | 1 | **52** |
| 1947 | **31897** | **11** | 3 | **52** |
| 1948 | **31898** | **10** | 1 | **52** |
| 1949 | **31899** | **12** | 5 | **52** |
| 1950 | **31900** | **9** | 2 | **52** |
| 1951 | **31901** | **10** | 1 | **52** |
| 1952 | **31902** | **10** | 1 | **52** |
| 1953 | **31903** | **10** | 1 | **52** |
| 1954 | **31904** | **10** | 1 | **52** |
| 1955 | **31905** | **10** | 1 | **52** |
| 1956 | **31906** | **10** | 1 | **456** |
| 1957 | **31907** | **10** | 1 | **456** |
| 1958 | **31908** | **10** | 1 | **456** |
| 1959 | **31909** | **9** | 2 | **52** |
| 1960 | **31910** | **10** | 1 | **52** |
| 1961 | **31911** | **9** | 2 | **52** |
| 1962 | **31912** | **9** | 2 | **52** |
| 1963 | **31913** | **10** | 1 | **52** |
| 1964 | **31914** | **9** | 2 | **456** |
| 1965 | **31915** | **10** | 1 | **52** |
| 1966 | **31916** | **10** | 1 | **52** |
| 1967 | **31917** | **10** | 1 | **456** |
| 1968 | **31918** | **10** | 1 | **456** |
| 1969 | **31919** | **10** | 1 | **458** |
| 1970 | **31920** | **9** | 2 | **52** |
| 1971 | **31921** | **10** | 1 | **52** |
| 1972 | **31922** | **ND** | - | **52** |
| 1973 | **31923** | **ND** | - | **52** |
| 1974 | **31924** | **ND** | - | **458(5)** |
| 1975 | **31925** | **10** | 1 | **52** |
| 1976 | **31926** | **7** | 7 | **244** |
| 1977 | **31927** | **10** | 1 | **52** |
| 1978 | **31928** | **9** | 2 | **52** |

| No. | MGAS number(1) | Number of Ts(2) | Allele number(3) | MLST type |
|---|---|---|---|---|
| 1979 | **31929** | **11** | 3 | **52** |
| 1980 | **31930** | **10** | 1 | **52** |
| 1981 | **31931** | **10** | 1 | **52** |
| 1982 | **31932** | **10** | 1 | **52** |
| 1983 | **31933** | **10** | 1 | **52** |
| 1984 | **31934** | **10** | 1 | **52** |
| 1985 | **31935** | **9** | 2 | **52** |
| 1986 | **31936** | **10** | 1 | **52** |
| 1987 | **31937** | **9** | 2 | **52** |
| 1988 | **31938** | **10** | 1 | **52** |
| 1989 | **31939** | **10** | 1 | **52** |
| 1990 | **31940** | **10** | 1 | **52** |
| 1991 | **31941** | **9** | 2 | **52** |
| 1992 | **31942** | **10** | 1 | **52** |
| 1993 | **31943** | **9** | 2 | **52** |
| 1994 | **31944** | **9** | 2 | **52** |
| 1995 | **31945** | **10** | 1 | **52** |
| 1996 | **31946** | **10** | 1 | **458** |
| 1997 | **31947** | **9** | 2 | **52** |
| 1998 | **31948** | **10** | 1 | **52** |
| 1999 | **31949** | **10** | 1 | **456** |
| 2000 | **31950** | **10** | 1 | **52** |
| 2001 | **31951** | **10** | 1 | **458** |
| 2002 | **31952** | **10** | 1 | **52** |
| 2003 | **31953** | **10** | 1 | **52** |
| 2004 | **31954** | **10** | 1 | **52** |
| 2005 | **31955** | **9** | 2 | **52** |
| 2006 | **31956** | **9** | 2 | **52** |
| 2007 | **31957** | **9** | 2 | **52** |
| 2008 | **31958** | **9** | 2 | **52** |
| 2009 | **31959** | **11** | 3 | **458** |
| 2010 | **31960** | **10** | 9 | **52** |
| 2011 | **31961** | **10** | 1 | **52** |
| 2012 | **31962** | **10** | 1 | **52** |
| 2013 | **31963** | **11** | 3 | **52** |
| 2014 | **31964** | **9** | 2 | **52** |
| 2015 | **31965** | **10** | 1 | **52** |
| 2016 | **31966** | **9** | 2 | **52** |
| 2017 | **31967** | **10** | 1 | **458** |
| 2018 | **31968** | **10** | 9 | **52** |
| 2019 | **31969** | **9** | 2 | **52** |
| 2020 | **31970** | **10** | 1 | **52** |
| 2021 | **31971** | **11** | 28 | **52** |
| 2022 | **31972** | **11** | 3 | **52** |

| No. | MGAS number(1) | Number of Ts(2) | Allele number(3) | MLST type |
|---|---|---|---|---|
| 2023 | **31973** | **11** | 3 | **52** |
| 2024 | **31974** | **10** | 10 | **52** |
| 2025 | **31975** | **9** | 2 | **52** |
| 2026 | **31976** | **10** | 1 | **52** |
| 2027 | **31977** | **10** | 1 | **458** |
| 2028 | **31978** | **9** | 6 | **52** |
| 2029 | **31979** | **9** | 2 | **52** |
| 2030 | **31980** | **10** | 1 | **458** |
| 2031 | **31981** | **9** | 2 | **52** |
| 2032 | **31982** | **9** | 2 | **52** |
| 2033 | **31983** | **10** | 1 | **838** |
| 2034 | **31984** | **10** | 1 | **52** |
| 2035 | **31985** | **9** | 2 | **52** |
| 2036 | **31986** | **10** | 1 | **52** |
| 2037 | **31987** | **10** | 1 | **52** |
| 2038 | **31988** | **9** | 2 | **52** |
| 2039 | **31989** | **10** | 1 | **458** |
| 2040 | **31990** | **9** | 2 | **52** |
| 2041 | **31991** | **9** | 2 | **52** |
| 2042 | **31992** | **9** | 2 | **52** |
| 2043 | **31993** | **9** | 2 | **52** |
| 2044 | **31994** | **10** | 1 | **458** |
| 2045 | **31995** | **9** | 2 | **52** |
| 2046 | **31996** | **10** | 1 | **52** |
| 2047 | **31997** | **10** | 1 | **458** |
| 2048 | **31998** | **10** | 1 | **52** |
| 2049 | **31999** | **9** | 2 | **456** |
| 2050 | **32001** | **10** | 1 | **52** |
| 2051 | **32002** | **10** | 1 | **52** |
| 2052 | **32003** | **10** | 1 | **52** |
| 2053 | **32004** | **10** | 1 | **458** |
| 2054 | **32005** | **10** | 1 | **458** |
| 2055 | **32007** | **10** | 1 | **458** |
| 2056 | **32008** | **9** | 2 | **458** |
| 2057 | **32009** | **9** | 2 | **456** |
| 2058 | **32010** | **10** | 1 | **52** |
| 2059 | **32011** | **11** | 3 | **458**(13) |
| 2060 | **32012** | **9** | 2 | **52** |
| 2061 | **32013** | **10** | 9 | **52** |
| 2062 | **32014** | **10** | 9 | **52** |
| 2063 | **32015** | **10** | 1 | **52** |
| 2064 | **32016** | **10** | 1 | **52** |
| 2065 | **32017** | **10** | 1 | **52** |
| 2066 | **32018** | **10** | 1 | **52** |

| No. | MGAS number[1] | Number of Ts[2] | Allele number[3] | MLST type |
|---|---|---|---|---|
| 2067 | **32019** | **10** | 1 | **458** |
| 2068 | **32020** | **10** | 1 | **52** |
| 2069 | **32021** | **10** | 1 | **52** |
| 2070 | **32022** | **9** | 2 | **52** |
| 2071 | **32023** | **9** | 2 | **52** |
| 2072 | **32024** | **10** | 1 | **458** |
| 2073 | **32025** | **10** | 1 | **52** |
| 2074 | **32026** | **11** | 3 | **52** |
| 2075 | **32027** | **11** | 3 | **52** |
| 2076 | **32029** | **12** | 5 | **458** |
| 2077 | **32030** | **10** | 1 | **52** |
| 2078 | **32031** | **9** | 24 | **52** |
| 2079 | **32032** | **9** | 2 | **52** |
| 2080 | **32033** | **10** | 1 | **458** |
| 2081 | **32034** | **9** | 2 | **52** |
| 2082 | **32035** | **10** | 1 | **52** |
| 2083 | **32036** | **12** | 5 | **52** |
| 2084 | **32037** | **12** | 5 | **52** |
| 2085 | **32038** | **10** | 1 | **52** |
| 2086 | **32039** | **11** | 3 | **52** |
| 2087 | **32040** | **10** | 1 | **52** |
| 2088 | **32041** | **10** | 1 | **52** |
| 2089 | **32053** | **10** | 1 | **52** |
| 2090 | **32094** | **10** | 1 | **52** |
| 2091 | **32120** | **9** | 2 | **52** |
| 2092 | **32179** | **10** | 1 | **52** |
| 2093 | **32182** | **9** | 2 | **456** |
| 2094 | **32194** | **11** | 3 | **52** |
| 2095 | **32199** | **10** | 1 | **52** |

**(1)** MGAS number. Musser GAS strain number

**(2)** Number of T nucleotides in the homopolymeric tract upstream of *Spy1336/R28*

**(3)** Allele number for the 30 $HT_{Spy1336\text{-}7}$ alleles found in 2,074 *emm28* GAS invasive strains (Table S2)

**(4)** ND, not determined

**(5)** Low-depth bases or truncation at *murI*

**(6)** Low-depth bases or truncation at *gtr*

**(7)** SNP in *xpt*

[8] NF, MLST allele not found in the SRST2 database

[9] SNP in *mutS*

[10] SNP in *gtr*

[11] Truncation or large deletion in *mutS*

[12] Indel in *yqiL*

[13] SNP in *yqiL*

[14] SNP in *gki*

[15] Multiple SNPs and deletions

[16] SNP in *recP*

[17] Three indels in *mutS*

### Table S2. HT$_{Spy1336-7}$ alleles found in 2,074 *emm28* GAS invasive strains

| Allele[1] | DNA Sequence[2] | | HT length | Count[3] |
|---|---|---|---|---|
| 1[4] | TTTTATCTAATCTAATCTGCTTTTTTTTTT | ATATATAATTTGACTTTTTC | 10 | **1226** |
| 2 | TTTTATCTAATCTAATCTGCTTTTTTTTT | ATATATAATTTGACTTTTTC | 9 | **650** |
| 3 | TTTTATCTAATCTAATCTGCTTTTTTTTTTT | ATATATAATTTGACTTTTTC | 11 | **128** |
| 4 | TTTTATCTAATCTAATCTGCTTTTTTTTAT | ATATATAATTTGACTTTTTC | 8 | **15** |
| 5 | TTTTATCTAATCTAATCTGCTTTTTTTTTTTT | ATATATAATTTGACTTTTTC | 12 | **11** |
| 6 | TTTTATCTAATCTAATCTGCTTTTTTTTTAT | ATATATAATTTGACTTTTTC | 9 | **7** |
| 7 | TTTTATCTAATCTAATCTGCTTTTTTTATAT | ATATATAATTTGACTTTTTC | 7 | **5** |
| 8 | TTTTATCTAATCTAATCTGCTTTTTTTT | ATATATAATTTGACTTTTTC | 8 | **4** |
| 9 | TTTTATCTAATCTAATCTGCTTTTTTTTTT | ATATGTAATTTGACTTTTTC | 10 | **4** |
| 10 | ----ATCTAATCTAATCTGCTTTTTTTTTT | ATATATAATTTGACTTTTTC | 10 | **2** |
| 11 | TTTTATCTAATCTAATCTGCTTTTTTTTT | ATATATAATTTGACTTTTT- | 9 | **2** |
| 12 | TTTTATCTAATCTAATCTGCTTTTTTTTTT | ATATATAATTTGACTTTTT- | 10 | **2** |
| 13 | ---------ATCTAATCTGCTTTTTTTTTT | ATATATAATTTGACTTTTTC | 10 | **1** |
| 14 | -------TAATCTAATCTTCTTTTTTTTTT | ATATATAATTTGACTTTTTC | 10 | **1** |
| 15 | ---TATCTAATCTAATCTGCTTTTTTTTT | ATATATAATTTGACTTTTTC | 9 | **1** |
| 16 | ---TATCTAATCTAATCTGCTTTTTTTTTT | ATATATAATTTGACTTTTTC | 10 | **1** |
| 17 | TTTTATCTAATATAATCTGCTTTTTTTTTT | ATATATAATTTGACTTTTTC | 10 | **1** |
| 18 | TTTTATCTAATCAAATCTGCTTTTTTTTT | ATATATAATTTGACTTTTTC | 9 | **1** |
| 19 | TTTTATCTAATCTAATCTGCCTTTTTTTTT | ATATATAATTTGACTTTTTC | 9 | **1** |
| 20 | TTTTATCTAATCTAATCTGCTTTTTTATAT | ATATATAATTTGACTTTTTC | 6 | **1** |
| 21 | TTTTATCTAATCTAATCTGCTTTTTTTAT | ATATATAATTTGACTTTTTC | 7 | **1** |
| 22 | TTTTATCTAATCTAATCTGCTTTTTTTATT | ATATATAATTTGACTTTTTC | 7 | **1** |
| 23 | TTTTATCTAATCTAATCTGCTTTTTTTT | ATATATAATTTGACTTTTT- | 8 | **1** |
| 24 | TTTTATCTAATCTAATCTGCTTTTTTTTT | ATATGTAATTTGACTTTTTC | 9 | **1** |
| 25 | TTTTATCTAATCTAATCTGCTTTTTTTTTT | ATATTTAATTTGACTTTTTC | 10 | **1** |
| 26 | TTTTATCTAATCTAATCTGCTTTTTTTTTT | ATTTATAATTTGACTTTTTC | 10 | **1** |
| 27 | TTTTATCTAATCTAATCTGCTTTTTTTTTTT | --ATAATTTGACTTTTTC | 11 | **1** |
| 28 | TTTTATCTAATCTAATCTGCTTTTTTTTTTT | ATATGTAATTTGACTTTTTC | 11 | **1** |
| 29 | TTTTATCTAATCTAATCTGCTTTTTTTTTTTTT | ATATATAATTTGACTTTTTC | 13 | **1** |
| 30 | TTTTATCTAATCTATTCTGCTTTTTTTTTT | ATATATAATTTGACTTTTTC | 10 | **1** |
| Total | | | | **2074** |

**[1] Allele number**

**[2] DNA sequence comprising 20 nucleotides upstream and downstream from HT$_{Spy1336-7}$**

**[3] Count refers to the number of strains found to contain a specific allele.**

**[4] Alleles in grey contain indels in HT$_{Spy1336-7}$ exclusively, and are shown in Figure 1D.**

**SNPs and additional indels are indicated in red. Deletions are indicated by -**

**Table S3. Number of $T_{R28}$ repeats in 493 *emm28* invasive strains**

| No. | MGAS number | $TR_{R28}$ number | Number of Ts in $HT_{Spy1336-7}$ |
|---|---|---|---|
| 1 | **7867** | **1** | 10 |
| 2 | **8417** | **1** | 10 |
| 3 | **10820** | **1** | 10 |
| 4 | **11115** | **1** | 10 |
| 5 | **28035** | **1** | 10 |
| 6 | **28118** | **1** | 9 |
| 7 | **28201** | **1** | 10 |
| 8 | **28374** | **1** | 9 |
| 9 | **28463** | **1** | 10 |
| 10 | **28550** | **1** | 9 |
| 11 | **28776** | **1** | 10 |
| 12 | **28788** | **1** | 9 |
| 13 | **28935** | **1** | 10 |
| 14 | **29069** | **1** | 10 |
| 15 | **29080** | **1** | 9 |
| 16 | **29164** | **1** | 10 |
| 17 | **29431** | **1** | 10 |
| 18 | **29569** | **1** | 9 |
| 19 | **29587** | **1** | 10 |
| 20 | **11147** | **2** | 9 |
| 21 | **11151** | **2** | 9 |
| 22 | **12231** | **2** | 10 |
| 23 | **27846** | **2** | 10 |
| 24 | **28008** | **2** | 10 |
| 25 | **28323** | **2** | 10 |
| 26 | **28424** | **2** | 8 |
| 27 | **28484** | **2** | 10 |
| 28 | **28530** | **2** | 10 |
| 29 | **28633** | **2** | 10 |
| 30 | **28648** | **2** | 10 |
| 31 | **28764** | **2** | 10 |
| 32 | **28811** | **2** | 9 |
| 33 | **28842** | **2** | 10 |
| 34 | **29073** | **2** | 11 |
| 35 | **29074** | **2** | 9 |
| 36 | **29117** | **2** | 9 |

| No. | MGAS number | $TR_{R28}$ number | Number of Ts in $HT_{Spy1336-7}$ |
|---|---|---|---|
| 37 | **29253** | **2** | 9 |
| 38 | **29268** | **2** | 10 |
| 39 | **29276** | **2** | 10 |
| 40 | **29321** | **2** | 10 |
| 41 | **29372** | **2** | 9 |
| 42 | **29412** | **2** | 9 |
| 43 | **29611** | **2** | 11 |
| 44 | **7912** | **3** | 10 |
| 45 | **7927** | **3** | 10 |
| 46 | **7941** | **3** | 10 |
| 47 | **10757** | **3** | 10 |
| 48 | **10771** | **3** | 10 |
| 49 | **12254** | **3** | 10 |
| 50 | **28058** | **3** | 9 |
| 51 | **28059** | **3** | 10 |
| 52 | **28225** | **3** | 9 |
| 53 | **28232** | **3** | 11 |
| 54 | **28249** | **3** | 11 |
| 55 | **28273** | **3** | 11 |
| 56 | **28296** | **3** | 10 |
| 57 | **28305** | **3** | 10 |
| 58 | **28307** | **3** | 9 |
| 59 | **28331** | **3** | 10 |
| 60 | **28334** | **3** | 10 |
| 61 | **28356** | **3** | 9 |
| 62 | **28362** | **3** | 10 |
| 63 | **28425** | **3** | 10 |
| 64 | **28430** | **3** | 10 |
| 65 | **28443** | **3** | 9 |
| 66 | **28562** | **3** | 10 |
| 67 | **28566** | **3** | 9 |
| 68 | **28718** | **3** | 10 |
| 69 | **29109** | **3** | 9 |
| 70 | **29299** | **3** | 10 |
| 71 | **29338** | **3** | 10 |
| 72 | **29394** | **3** | 10 |
| 73 | **29476** | **3** | 10 |

| No. | MGAS number | $TR_{R28}$ number | Number of Ts in $HT_{Spy1336-7}$ |
|---|---|---|---|
| 74 | **29501** | **3** | 10 |
| 75 | **29508** | **3** | 11 |
| 76 | **29553** | **3** | 10 |
| 77 | **29570** | **3** | 9 |
| 78 | **29608** | **3** | 9 |
| 79 | **29617** | **3** | 10 |
| 80 | **7872** | **4** | 9 |
| 81 | **7886** | **4** | 10 |
| 82 | **7960** | **4** | 9 |
| 83 | **7995** | **4** | 10 |
| 84 | **8375** | **4** | 10 |
| 85 | **8447** | **4** | 9 |
| 86 | **11140** | **4** | 10 |
| 87 | **11149** | **4** | 10 |
| 88 | **27951** | **4** | 10 |
| 89 | **27970** | **4** | 9 |
| 90 | **27979** | **4** | 10 |
| 91 | **27983** | **4** | 11 |
| 92 | **28207** | **4** | 9 |
| 93 | **28271** | **4** | 10 |
| 94 | **28341** | **4** | 10 |
| 95 | **28367** | **4** | 10 |
| 96 | **28418** | **4** | 10 |
| 97 | **28440** | **4** | 10 |
| 98 | **28441** | **4** | 9 |
| 99 | **28446** | **4** | 10 |
| 100 | **28447** | **4** | 9 |
| 101 | **28481** | **4** | 9 |
| 102 | **28576** | **4** | 10 |
| 103 | **28650** | **4** | 10 |
| 104 | **28669** | **4** | 9 |
| 105 | **28687** | **4** | 10 |
| 106 | **28708** | **4** | 9 |
| 107 | **28724** | **4** | 10 |
| 108 | **28772** | **4** | 10 |
| 109 | **28784** | **4** | 10 |
| 110 | **28785** | **4** | 10 |

| No. | MGAS number | $TR_{R28}$ number | Number of Ts in $HT_{Spy1336-7}$ |
|---|---|---|---|
| 111 | **28822** | **4** | 10 |
| 112 | **28826** | **4** | 9 |
| 113 | **28880** | **4** | 10 |
| 114 | **28897** | **4** | 9 |
| 115 | **29050** | **4** | 10 |
| 116 | **29108** | **4** | 10 |
| 117 | **29138** | **4** | 9 |
| 118 | **29170** | **4** | 9 |
| 119 | **29184** | **4** | 10 |
| 120 | **29212** | **4** | 9 |
| 121 | **29342** | **4** | 10 |
| 122 | **29351** | **4** | 11 |
| 123 | **29384** | **4** | 9 |
| 124 | **29464** | **4** | 9 |
| 125 | **10819** | **5** | 10 |
| 126 | **10824** | **5** | 10 |
| 127 | **10826** | **5** | 9 |
| 128 | **28020** | **5** | 10 |
| 129 | **28029** | **5** | 10 |
| 130 | **28121** | **5** | 10 |
| 131 | **28127** | **5** | 9 |
| 132 | **28185** | **5** | 10 |
| 133 | **28402** | **5** | 10 |
| 134 | **28415** | **5** | 10 |
| 135 | **28461** | **5** | 10 |
| 136 | **28479** | **5** | 10 |
| 137 | **28543** | **5** | 10 |
| 138 | **28596** | **5** | 10 |
| 139 | **28622** | **5** | 10 |
| 140 | **28722** | **5** | 11 |
| 141 | **28751** | **5** | 10 |
| 142 | **28752** | **5** | 10 |
| 143 | **28804** | **5** | 10 |
| 144 | **28805** | **5** | 10 |
| 145 | **28942** | **5** | 10 |
| 146 | **29159** | **5** | 10 |
| 147 | **29198** | **5** | 10 |

| No. | MGAS number | $TR_{R28}$ number | Number of Ts in $HT_{Spy1336-7}$ |
|---|---|---|---|
| 148 | **29209** | **5** | 11 |
| 149 | **29305** | **5** | 10 |
| 150 | **29371** | **5** | 10 |
| 151 | **29423** | **5** | 10 |
| 152 | **29485** | **5** | 10 |
| 153 | **29537** | **5** | 10 |
| 154 | **7893** | **6** | 10 |
| 155 | **7946** | **6** | 10 |
| 156 | **10817** | **6** | 10 |
| 157 | **10827** | **6** | 10 |
| 158 | **27865** | **6** | 9 |
| 159 | **28047** | **6** | 10 |
| 160 | **28085** | **6** | 10 |
| 161 | **28142** | **6** | 10 |
| 162 | **28204** | **6** | 9 |
| 163 | **28278** | **6** | 10 |
| 164 | **28338** | **6** | 10 |
| 165 | **28347** | **6** | 10 |
| 166 | **28455** | **6** | 10 |
| 167 | **28462** | **6** | 10 |
| 168 | **28553** | **6** | 11 |
| 169 | **28567** | **6** | 10 |
| 170 | **28645** | **6** | 11 |
| 171 | **28647** | **6** | 10 |
| 172 | **28688** | **6** | 10 |
| 173 | **28710** | **6** | 10 |
| 174 | **28840** | **6** | 10 |
| 175 | **28883** | **6** | 10 |
| 176 | **29075** | **6** | 10 |
| 177 | **29133** | **6** | 10 |
| 178 | **29140** | **6** | 9 |
| 179 | **29243** | **6** | 10 |
| 180 | **29258** | **6** | 9 |
| 181 | **29293** | **6** | 10 |
| 182 | **29323** | **6** | 10 |
| 183 | **29474** | **6** | 9 |
| 184 | **29491** | **6** | 11 |

| No. | MGAS number | $TR_{R28}$ number | Number of Ts in $HT_{Spy1336-7}$ |
|---|---|---|---|
| 185 | **29502** | **6** | 9 |
| 186 | **29546** | **6** | 10 |
| 187 | **29613** | **6** | 10 |
| 188 | **7869** | **7** | 10 |
| 189 | **7976** | **7** | 10 |
| 190 | **7982** | **7** | 10 |
| 191 | **8358** | **7** | 9 |
| 192 | **28106** | **7** | 10 |
| 193 | **28208** | **7** | 9 |
| 194 | **28230** | **7** | 10 |
| 195 | **28360** | **7** | 9 |
| 196 | **28411** | **7** | 10 |
| 197 | **28426** | **7** | 9 |
| 198 | **28504** | **7** | 9 |
| 199 | **28508** | **7** | 9 |
| 200 | **28533** | **7** | 10 |
| 201 | **28549** | **7** | 9 |
| 202 | **28582** | **7** | 10 |
| 203 | **28589** | **7** | 10 |
| 204 | **28781** | **7** | 10 |
| 205 | **28787** | **7** | 10 |
| 206 | **28809** | **7** | 10 |
| 207 | **28845** | **7** | 9 |
| 208 | **28851** | **7** | 9 |
| 209 | **28872** | **7** | 8 |
| 210 | **28904** | **7** | 10 |
| 211 | **29051** | **7** | 10 |
| 212 | **29093** | **7** | 9 |
| 213 | **29180** | **7** | 9 |
| 214 | **29181** | **7** | 10 |
| 215 | **29324** | **7** | 9 |
| 216 | **29510** | **7** | 9 |
| 217 | **7956** | **8** | 10 |
| 218 | **8345** | **8** | 10 |
| 219 | **12247** | **8** | 9 |
| 220 | **27937** | **8** | 10 |
| 221 | **27956** | **8** | 10 |

| No. | MGAS number | $TR_{R28}$ number | Number of Ts in $HT_{Spy1336-7}$ |
|---|---|---|---|
| 222 | **27990** | **8** | 10 |
| 223 | **28004** | **8** | 9 |
| 224 | **28048** | **8** | 9 |
| 225 | **28082** | **8** | 9 |
| 226 | **28164** | **8** | 11 |
| 227 | **28209** | **8** | 10 |
| 228 | **28295** | **8** | 10 |
| 229 | **28321** | **8** | 10 |
| 230 | **28400** | **8** | 10 |
| 231 | **28585** | **8** | 9 |
| 232 | **28609** | **8** | 10 |
| 233 | **28612** | **8** | 10 |
| 234 | **28626** | **8** | 11 |
| 235 | **28801** | **8** | 9 |
| 236 | **28836** | **8** | 9 |
| 237 | **28869** | **8** | 9 |
| 238 | **28886** | **8** | 10 |
| 239 | **28900** | **8** | 9 |
| 240 | **28921** | **8** | 10 |
| 241 | **28924** | **8** | 10 |
| 242 | **29076** | **8** | 9 |
| 243 | **29095** | **8** | 10 |
| 244 | **29105** | **8** | 10 |
| 245 | **29188** | **8** | 9 |
| 246 | **29263** | **8** | 10 |
| 247 | **29300** | **8** | 9 |
| 248 | **29360** | **8** | 10 |
| 249 | **29381** | **8** | 9 |
| 250 | **29402** | **8** | 10 |
| 251 | **29416** | **8** | 10 |
| 252 | **29488** | **8** | 10 |
| 253 | **29504** | **8** | 10 |
| 254 | **29507** | **8** | 10 |
| 255 | **29534** | **8** | 9 |
| 256 | **29588** | **8** | 10 |
| 257 | **29594** | **8** | 11 |
| 258 | **29620** | **8** | 10 |

| No. | MGAS number | $TR_{R28}$ number | Number of Ts in $HT_{Spy1336-7}$ |
|---|---|---|---|
| 259 | **7890** | **9** | 10 |
| 260 | **7980** | **9** | 10 |
| 261 | **7994** | **9** | 10 |
| 262 | **8007** | **9** | 10 |
| 263 | **8009** | **9** | 9 |
| 264 | **10807** | **9** | 9 |
| 265 | **12250** | **9** | 9 |
| 266 | **27767** | **9** | 10 |
| 267 | **27787** | **9** | 10 |
| 268 | **27946** | **9** | 10 |
| 269 | **27967** | **9** | 10 |
| 270 | **27981** | **9** | 9 |
| 271 | **27993** | **9** | 9 |
| 272 | **28007** | **9** | 9 |
| 273 | **28022** | **9** | 10 |
| 274 | **28032** | **9** | 10 |
| 275 | **28046** | **9** | 10 |
| 276 | **28049** | **9** | 8 |
| 277 | **28074** | **9** | 10 |
| 278 | **28081** | **9** | 9 |
| 279 | **28086** | **9** | 10 |
| 280 | **28115** | **9** | 10 |
| 281 | **28138** | **9** | 9 |
| 282 | **28172** | **9** | 9 |
| 283 | **28241** | **9** | 9 |
| 284 | **28248** | **9** | 10 |
| 285 | **28265** | **9** | 10 |
| 286 | **28268** | **9** | 10 |
| 287 | **28287** | **9** | 10 |
| 288 | **28330** | **9** | 10 |
| 289 | **28386** | **9** | 11 |
| 290 | **28467** | **9** | 10 |
| 291 | **28471** | **9** | 10 |
| 292 | **28480** | **9** | 10 |
| 293 | **28482** | **9** | 10 |
| 294 | **28505** | **9** | 10 |
| 295 | **28510** | **9** | 10 |

| No. | MGAS number | $TR_{R28}$ number | Number of Ts in $HT_{Spy1336-7}$ |
|---|---|---|---|
| 296 | **28512** | **9** | 11 |
| 297 | **28513** | **9** | 10 |
| 298 | **28523** | **9** | 10 |
| 299 | **28630** | **9** | 10 |
| 300 | **28662** | **9** | 9 |
| 301 | **28707** | **9** | 10 |
| 302 | **28731** | **9** | 9 |
| 303 | **28744** | **9** | 10 |
| 304 | **28748** | **9** | 9 |
| 305 | **28798** | **9** | 10 |
| 306 | **28906** | **9** | 9 |
| 307 | **29161** | **9** | 10 |
| 308 | **29162** | **9** | 10 |
| 309 | **29214** | **9** | 9 |
| 310 | **29240** | **9** | 9 |
| 311 | **29254** | **9** | 10 |
| 312 | **29326** | **9** | 9 |
| 313 | **29331** | **9** | 10 |
| 314 | **29377** | **9** | 9 |
| 315 | **29379** | **9** | 10 |
| 316 | **29382** | **9** | 9 |
| 317 | **29387** | **9** | 10 |
| 318 | **29409** | **9** | 10 |
| 319 | **29426** | **9** | 10 |
| 320 | **29429** | **9** | 10 |
| 321 | **29444** | **9** | 10 |
| 322 | **29511** | **9** | 9 |
| 323 | **29542** | **9** | 9 |
| 324 | **29564** | **9** | 9 |
| 325 | **29583** | **9** | 10 |
| 326 | **7888** | **10** | 10 |
| 327 | **7891** | **10** | 9 |
| 328 | **7914** | **10** | 10 |
| 329 | **7921** | **10** | 10 |
| 330 | **7935** | **10** | 10 |
| 331 | **7959** | **10** | 10 |
| 332 | **7973** | **10** | 10 |

| No. | MGAS number | $TR_{R28}$ number | Number of Ts in $HT_{Spy1336-7}$ |
|---|---|---|---|
| 333 | **8012** | **10** | 10 |
| 334 | **8347** | **10** | 11 |
| 335 | **8365** | **10** | 10 |
| 336 | **8396** | **10** | 10 |
| 337 | **10786** | **10** | 10 |
| 338 | **10793** | **10** | 9 |
| 339 | **10799** | **10** | 10 |
| 340 | **10812** | **10** | 9 |
| 341 | **11103** | **10** | 9 |
| 342 | **11107** | **10** | 10 |
| 343 | **27961** | **10** | 9 |
| 344 | **27962** | **10** | 10 |
| 345 | **28031** | **10** | 9 |
| 346 | **28065** | **10** | 10 |
| 347 | **28107** | **10** | 10 |
| 348 | **28108** | **10** | 9 |
| 349 | **28117** | **10** | 9 |
| 350 | **28191** | **10** | 10 |
| 351 | **28217** | **10** | 10 |
| 352 | **28254** | **10** | 9 |
| 353 | **28261** | **10** | 10 |
| 354 | **28309** | **10** | 9 |
| 355 | **28315** | **10** | 9 |
| 356 | **28392** | **10** | 10 |
| 357 | **28397** | **10** | 9 |
| 358 | **28429** | **10** | 9 |
| 359 | **28473** | **10** | 10 |
| 360 | **28477** | **10** | 10 |
| 361 | **28501** | **10** | 10 |
| 362 | **28536** | **10** | 9 |
| 363 | **28640** | **10** | 10 |
| 364 | **28653** | **10** | 10 |
| 365 | **28654** | **10** | 9 |
| 366 | **28670** | **10** | 10 |
| 367 | **28686** | **10** | 9 |
| 368 | **28715** | **10** | 10 |
| 369 | **28728** | **10** | 10 |

| No. | MGAS number | $TR_{R28}$ number | Number of Ts in $HT_{Spy1336-7}$ |
|---|---|---|---|
| 370 | **28739** | **10** | 9 |
| 371 | **28746** | **10** | 9 |
| 372 | **28747** | **10** | 10 |
| 373 | **28825** | **10** | 10 |
| 374 | **28831** | **10** | 10 |
| 375 | **28864** | **10** | 9 |
| 376 | **28879** | **10** | 9 |
| 377 | **28898** | **10** | 9 |
| 378 | **28925** | **10** | 11 |
| 379 | **28949** | **10** | 9 |
| 380 | **29041** | **10** | 10 |
| 381 | **29045** | **10** | 10 |
| 382 | **29054** | **10** | 10 |
| 383 | **29068** | **10** | 10 |
| 384 | **29086** | **10** | 10 |
| 385 | **29124** | **10** | 10 |
| 386 | **29125** | **10** | 10 |
| 387 | **29201** | **10** | 10 |
| 388 | **29216** | **10** | 10 |
| 389 | **29221** | **10** | 9 |
| 390 | **29233** | **10** | 9 |
| 391 | **29238** | **10** | 10 |
| 392 | **29281** | **10** | 10 |
| 393 | **29310** | **10** | 10 |
| 394 | **29325** | **10** | 9 |
| 395 | **29330** | **10** | 10 |
| 396 | **29347** | **10** | 11 |
| 397 | **29404** | **10** | 9 |
| 398 | **29517** | **10** | 10 |
| 399 | **29559** | **10** | 9 |
| 400 | **29573** | **10** | 9 |
| 401 | **7918** | **11** | 10 |
| 402 | **7922** | **11** | 9 |
| 403 | **7987** | **11** | 10 |
| 404 | **8350** | **11** | 10 |
| 405 | **8374** | **11** | 10 |
| 406 | **10764** | **11** | 10 |

| No. | MGAS number | $TR_{R28}$ number | Number of Ts in $HT_{Spy1336\text{-}7}$ |
|---|---|---|---|
| 407 | **10802** | **11** | 10 |
| 408 | **12226** | **11** | 10 |
| 409 | **12248** | **11** | 9 |
| 410 | **27895** | **11** | 10 |
| 411 | **28041** | **11** | 9 |
| 412 | **28255** | **11** | 9 |
| 413 | **28317** | **11** | 10 |
| 414 | **28449** | **11** | 10 |
| 415 | **28557** | **11** | 10 |
| 416 | **28657** | **11** | 11 |
| 417 | **28661** | **11** | 9 |
| 418 | **28673** | **11** | 10 |
| 419 | **28699** | **11** | 10 |
| 420 | **28716** | **11** | 9 |
| 421 | **28760** | **11** | 11 |
| 422 | **28907** | **11** | 10 |
| 423 | **28932** | **11** | 10 |
| 424 | **29043** | **11** | 9 |
| 425 | **29061** | **11** | 10 |
| 426 | **29094** | **11** | 10 |
| 427 | **29143** | **11** | 9 |
| 428 | **29147** | **11** | 10 |
| 429 | **29230** | **11** | 10 |
| 430 | **29284** | **11** | 10 |
| 431 | **29286** | **11** | 10 |
| 432 | **29292** | **11** | 10 |
| 433 | **29314** | **11** | 10 |
| 434 | **29447** | **11** | 13 |
| 435 | **29494** | **11** | 9 |
| 436 | **29528** | **11** | 10 |
| 437 | **29599** | **11** | 10 |
| 438 | **29633** | **11** | 10 |
| 439 | **7865** | **12** | 9 |
| 440 | **7906** | **12** | 10 |
| 441 | **7972** | **12** | 10 |
| 442 | **8015** | **12** | 10 |
| 443 | **11052** | **12** | 10 |

| No. | MGAS number | $TR_{R28}$ number | Number of Ts in $HT_{Spy1336-7}$ |
|---|---|---|---|
| 444 | **11087** | **12** | 9 |
| 445 | **11098** | **12** | 9 |
| 446 | **11100** | **12** | 9 |
| 447 | **11108** | **12** | 11 |
| 448 | **11113** | **12** | 10 |
| 449 | **11123** | **12** | 10 |
| 450 | **11128** | **12** | 10 |
| 451 | **11138** | **12** | 10 |
| 452 | **27784** | **12** | 9 |
| 453 | **27848** | **12** | 9 |
| 454 | **28078** | **12** | 10 |
| 455 | **28737** | **12** | 9 |
| 456 | **28766** | **12** | 9 |
| 457 | **29064** | **12** | 10 |
| 458 | **29088** | **12** | 9 |
| 459 | **29186** | **12** | 9 |
| 460 | **6180** | **13** | 10 |
| 461 | **7878** | **13** | 10 |
| 462 | **11121** | **13** | 10 |
| 463 | **27754** | **13** | 10 |
| 464 | **28948** | **13** | 9 |
| 465 | **29406** | **13** | 10 |
| 466 | **29538** | **13** | 10 |
| 467 | **29582** | **13** | 10 |
| 468 | **28131** | **14** | 10 |
| 469 | **29556** | **14** | 10 |
| 470 | **28403** | **15** | 10 |
| 471 | **28597** | **15** | 10 |
| 472 | **29486** | **15** | 10 |
| 473 | **28544** | **16** | 10 |
| 474 | **29393** | **16** | 9 |
| 475 | **28727** | **17** | 9 |
| 476 | **7932** | **ND**(1) | 10 |
| 477 | **10761** | **ND** | 9 |
| 478 | **10792** | **No RD2**(2) | - |
| 479 | **10815** | **ND** | 9 |
| 480 | **27813** | **ND** | 9 |

| No. | MGAS number | $TR_{R28}$ number | Number of Ts in $HT_{Spy1336-7}$ |
|---|---|---|---|
| 481 | **28259** | **ND** | 11 |
| 482 | **28363** | **ND** | 9 |
| 483 | **28575** | **No RD2** | - |
| 484 | **28916** | **ND** | 9 |
| 485 | **28928** | **ND** | 11 |
| 486 | **29127** | **ND** | 9 |
| 487 | **29277** | **ND** | 10 |
| 488 | **29336** | **ND** | 9 |
| 489 | **29357** | **ND** | 11 |
| 490 | **29522** | **ND** | 9 |
| 491 | **29525** | **ND** | 9 |
| 492 | **29536** | **ND** | 10 |
| 493 | **29548** | **ND** | 9 |

(1) Not determined

**(2)** The RD2 ICE element is not present

### Table S4. Differentially expressed genes comparing MGAS27961Δ*Spy1337* to the isogenic parental strain MGAS27961-10T

#### A. Upregulated genes.

| | Growth[1] | Spy | gene | FC[2] | Function |
|---|---|---|---|---|---|
| 1 | ME[3] | Spy0023 | *purL* | 2.2 | Phosphoribosylformylglycinamidine synthase |
| 2 | ME | Spy0169 | - | 1.7 | Transposase |
| 3 | ME | Spy0170 | - | 1.9 | Transposase |
| 4 | ME | Spy0184 | *rivR* | 1.7 | RofA-related transcriptional regulator |
| 5 | ME | Spy0478 | - | 2.1 | Thiamine transporter |
| 6 | ME | Spy0873 | - | 1.5 | Mg2+/citrate complex secondary transporter |
| 7 | ME | Spy0878 | *citD* | 1.5 | Citrate lyase acyl carrier protein |
| 8 | ME | Spy0899 | *fhs.1* | 1.8 | Formate--tetrahydrofolate ligase |
| 9 | ME | Spy1209 | - | 1.5 | Xaa-His dipeptidase |
| 10 | ME | Spy1210 | - | 1.5 | Arginine/ornithine antiporter |
| 1 | ES[4] | Spy0106 | - | 2.6 | Sortase |
| 2 | ES | Spy0107 | *cpa* | 1.7 | Collagen-binding protein |
| 3 | ES | Spy0108 | - | 1.6 | Signal peptidase I |
| 4 | ES | Spy0109 | - | 1.5 | Hypothetical protein |
| 5 | ES | Spy0110 | *eftLSL.B* | 1.5 | Hypothetical exported protein |
| 6 | ES | Spy0111 | - | 1.8 | Hypothetical protein |
| 7 | ES | Spy0112 | - | 2.5 | Transcriptional regulator, AraC family |
| 8 | ES | Spy0117 | *atoB* | 2.1 | Acetyl-CoA acetyltransferase |
| 9 | ES | Spy0118 | *atoD.2* | 2.0 | Acetate CoA-transferase alpha subunit |
| 10 | ES | Spy0119 | *atoD.1* | 2.0 | Acetyl-CoA:acetoacetyl-CoA transferase beta subunit |
| 11 | ES | Spy0121 | *ridA* | 3.3 | RidA-enamine deaminase-family protein |
| 12 | ES | Spy0122 | *sloR* | 4.3 | Transcriptional regulator |
| 13 | ES | Spy0123 | - | 4.3 | Hypothetical protein |
| 14 | ES | Spy0124 | *ntpI* | 4.5 | V-type sodium ATP synthase subunit I |
| 15 | ES | Spy0125 | *ntpK* | 4.7 | V-type sodium ATP synthase subunit K |
| 16 | ES | Spy0126 | *ntpE* | 5.0 | V-type sodium ATP synthase subunit E |
| 17 | ES | Spy0127 | *ntpC* | 4.9 | V-type ATP synthase subunit C |
| 18 | ES | Spy0128 | *ntpF* | 5.2 | V-type sodium ATP synthase subunit F |
| 19 | ES | Spy0129 | *ntpA* | 4.4 | V-type sodium ATP synthase subunit A |
| 20 | ES | Spy0130 | *ntpB* | 4.0 | V-type sodium ATP synthase subunit B |
| 21 | ES | Spy0131 | *ntpD* | 3.9 | V-type sodium ATP synthase subunit D |
| 22 | ES | Spy0134 | *purA* | 1.7 | Adenylosuccinate synthetase |
| 23 | ES | Spy0135 | - | 1.7 | Nucleoside-binding protein |
| 24 | ES | Spy0157 | *polA* | 1.7 | DNA polymerase I |
| 25 | ES | Spy0184 | *rivR* | 1.6 | RofA-related transcriptional regulator |
| 26 | ES | Spy0210 | - | 3.2 | Hypothetical membrane spanning protein |
| 27 | ES | Spy0211 | *nanH* | 3.4 | N-acetylneuraminate lyase |
| 28 | ES | Spy0212 | - | 3.7 | N-acetylmannosamine kinase |

| | Growth[1] | Spy | gene | FC[2] | Function |
|---|---|---|---|---|---|
| 29 | ES | Spy0213 | - | 2.0 | Transcriptional regulator, RpiR family |
| 30 | ES | Spy0287 | - | 1.6 | Acylphosphatase |
| 31 | ES | Spy0329 | *prtS* | 2.6 | Lactocepin |
| 32 | ES | Spy0335 | *nrdI* | 1.7 | Hypothetical protein |
| 33 | ES | Spy0336 | *nrdE.1* | 1.7 | Ribonucleoside-diphosphate reductase alpha chain |
| 34 | ES | Spy0338 | - | 1.7 | Hypothetical protein |
| 35 | ES | Spy0361 | *ftsK* | 1.5 | Cell division protein |
| 36 | ES | Spy0398 | - | 1.6 | ATPase |
| 37 | ES | Spy0409 | *gloA* | 1.5 | Lactoylglutathione lyase |
| 38 | ES | Spy0410 | - | 1.5 | NAD(P)H-dependent quinone reductase |
| 39 | ES | Spy0432 | - | 1.5 | Hypothetical protein |
| 40 | ES | Spy0433 | *metK1* | 1.6 | S-adenosylmethionine synthetase |
| 41 | ES | Spy0434 | - | 1.7 | Hypothetical membrane associated protein |
| 42 | ES | Spy0435 | - | 1.6 | Involved in cell wall biogenesis |
| 43 | ES | Spy0436 | - | 1.6 | Hypothetical protein |
| 44 | ES | Spy0498 | *agaD* | 3.3 | PTS system, N-acetylgalactosamine-specific IID component |
| 45 | ES | Spy0499 | *agaC* | 3.5 | PTS system, N-acetylgalactosamine-specific IIC component |
| 46 | ES | Spy0500 | *agaV* | 3.6 | PTS system, N-acetylgalactosamine-specific IIB component |
| 47 | ES | Spy0501 | *ugl* | 3.3 | Unsaturated glucuronyl hydrolase |
| 48 | ES | Spy0502 | *agaF* | 2.9 | PTS system, N-acetylgalactosamine-specific IIA component |
| 49 | ES | Spy0503 | *idnO* | 1.8 | Gluconate 5-dehydrogenase |
| 50 | ES | Spy0504 | - | 1.5 | Galactose-6-phosphate isomerase LacB subunit |
| 51 | ES | Spy0506 | *kgdA* | 1.5 | 2-dehydro-3-deoxyphosphogluconate aldolase |
| 52 | ES | Spy0536 | - | 1.8 | Transposase |
| 53 | ES | Spy0537 | - | 1.9 | Transposase |
| 54 | ES | Spy0538 | - | 1.8 | Transcriptional regulator |
| 55 | ES | Spy0540 | *sagA* | 1.8 | Streptolysin S precursor |
| 56 | ES | Spy0542 | *sagC* | 1.6 | Streptolysin S biosynthesis protein |
| 57 | ES | Spy0543 | *sagD* | 1.7 | Streptolysin S biosynthesis protein |
| 58 | ES | Spy0544 | *sagE* | 1.8 | Streptolysin S putative self-immunity protein |
| 59 | ES | Spy0545 | *sagF* | 1.8 | Streptolysin S biosynthesis protein |
| 60 | ES | Spy0546 | *sagG* | 1.9 | Streptolysin S export ATP-binding protein |
| 61 | ES | Spy0547 | *sagH* | 1.8 | Streptolysin S export transmembrane protein |
| 62 | ES | Spy0548 | *sagI* | 1.7 | Streptolysin S export transmembrane protein |
| 63 | ES | Spy0549 | - | 1.5 | Endonuclease/exonuclease/phosphatase family protein |
| 64 | ES | Spy0577 | *mscL* | 2.2 | Large-conductance mechanosensitive channel |
| 65 | ES | Spy0593 | *pepT* | 1.5 | Peptidase T |
| 66 | ES | Spy0594 | *ebsA* | 1.7 | Pore forming protein |
| 67 | ES | Spy0595 | - | 1.8 | Ferredoxin |
| 68 | ES | Spy0596 | - | 2.0 | Hypothetical membrane associated protein |
| 69 | ES | Spy0597 | *cmk* | 1.7 | Cytidylate kinase |
| 70 | ES | Spy0612 | *capA* | 1.8 | Capsule biosynthesis protein |
| 71 | ES | Spy0661 | - | 1.5 | Two component system histidine kinase |

| | Growth[1] | Spy | gene | FC[2] | Function |
|---|---|---|---|---|---|
| 72 | ES | Spy0674 | *clpL* | 3.8 | ATP-dependent protease ATP-binding subunit |
| 73 | ES | Spy0731 | *acoB* | 1.5 | Pyruvate dehydrogenase E1 component beta subunit |
| 74 | ES | Spy0732 | *acoC* | 1.6 | Dihydrolipoamide acetyltransferase component of pyruvate DH |
| 75 | ES | Spy0733 | - | 1.6 | Hypothetical protein |
| 76 | ES | Spy0734 | *acoL* | 1.6 | Dihydrolipoamide dehydrogenase |
| 77 | ES | Spy0735 | - | 1.6 | Hypothetical protein |
| 78 | ES | Spy0885 | *xerD* | 1.6 | Recombinase |
| 79 | ES | Spy0894 | *pdxK* | 1.9 | Hypothetical membrane spanning protein |
| 80 | ES | Spy0895 | - | 2.0 | Pyridoxine kinase |
| 81 | ES | Spy0896 | - | 2.2 | Transcriptional regulator, GntR family |
| 82 | ES | Spy0943 | - | 2.2 | General stress protein, Gls24 family |
| 83 | ES | Spy0944 | - | 2.3 | Hypothetical protein |
| 84 | ES | Spy0945 | - | 2.3 | General stress protein, Gls24 family |
| 85 | ES | Spy0946 | - | 2.4 | Hypothetical protein |
| 86 | ES | Spy0947 | - | 2.3 | Hypothetical protein |
| 87 | ES | Spy0948 | - | 2.0 | Integral membrane protein |
| 88 | ES | Spy0958 | *glmS* | 1.5 | Isomerizing glucosamine-fructose-6-P aminotransferase |
| 89 | ES | Spy0963 | - | 1.7 | Transcriptional regulator, GntR family |
| 90 | ES | Spy0964 | - | 1.9 | ABC transporter ATP-binding protein |
| 91 | ES | Spy0965 | - | 1.9 | ABC transporter permease protein |
| 92 | ES | Spy0999 | - | 2.0 | 6180.1 phage protein |
| 93 | ES | Spy1009 | - | 1.8 | 6180.1 phage protein |
| 94 | ES | Spy1012 | - | 1.8 | 6180.1 phage protein |
| 95 | ES | Spy1014 | - | 1.7 | 6180.1 phage protein |
| 96 | ES | Spy1015 | - | 2.0 | 6180.1 phage protein |
| 97 | ES | Spy1017 | - | 2.0 | 6180.1 phage protein |
| 98 | ES | Spy1018 | - | 1.9 | 6180.1 phage protein |
| 99 | ES | Spy1019 | - | 2.3 | 6180.1 phage protein |
| 100 | ES | Spy1020 | - | 2.2 | 6180.1 phage protein |
| 101 | ES | Spy1036 | *glgP* | 1.6 | Glycogen phosphorylase |
| 102 | ES | Spy1039 | *malE* | 1.9 | Maltose/maltodextrin-binding protein (ABC transporter) |
| 103 | ES | Spy1040 | *malF* | 1.5 | Maltose transport system permease protein (ABC transporter) |
| 104 | ES | Spy1041 | *malG* | 1.8 | Maltose transport system permease protein (ABC transporter) |
| 105 | ES | Spy1043 | *malA* | 2.7 | Maltodextrose utilization protein |
| 106 | ES | Spy1044 | *malD* | 2.9 | Maltodextrin transport system permease protein |
| 107 | ES | Spy1045 | *malC* | 2.9 | Maltose transport system permease protein |
| 108 | ES | Spy1060 | *celB* | 1.6 | PTS system, cellobiose-specific IIC component |
| 109 | ES | Spy1061 | - | 1.8 | Hypothetical protein |
| 110 | ES | Spy1062 | *celC* | 1.7 | PTS system, cellobiose-specific IIA component |
| 111 | ES | Spy1063 | *celA* | 1.7 | PTS system, cellobiose-specific IIB component |
| 112 | ES | Spy1064 | - | 1.6 | Transcription antiterminator, BglG family |
| 113 | ES | Spy1065 | - | 2.0 | Outer surface protein |
| 114 | ES | Spy1066 | *bglA.2* | 1.9 | Beta-glucosidase |

| | Growth[1] | Spy | gene | FC[2] | Function |
|---|---|---|---|---|---|
| 115 | ES | Spy1085 | - | 1.6 | RD.1 transcriptional regulator, MarR family |
| 116 | ES | Spy1123 | - | 1.6 | CPBP family intramembrane metalloprotease |
| 117 | ES | Spy1161 | - | 1.5 | Lead, cadmium, zinc and mercury transporting ATPase |
| 118 | ES | Spy1179 | *clpE* | 1.7 | ATP-dependent Clp protease ATP-binding subunit |
| 119 | ES | Spy1208 | *arcC* | 2.3 | Carbamate kinase |
| 120 | ES | Spy1209 | - | 2.3 | Xaa-His dipeptidase |
| 121 | ES | Spy1218 | - | 1.5 | Two-component sensor kinase |
| 122 | ES | Spy1289 | - | 1.9 | Hypothetical cytosolic protein |
| 123 | ES | Spy1290 | - | 2.0 | Hypothetical protein |
| 124 | ES | Spy1291 | - | 2.0 | Hypothetical protein |
| 125 | ES | Spy1292 | - | 1.8 | Hypothetical cytosolic protein |
| 126 | ES | Spy1293 | - | 1.6 | Hypothetical protein |
| 127 | ES | Spy1294 | - | 1.6 | Hypothetical protein |
| 128 | ES | Spy1295 | - | 1.6 | ATP-dependent RNA helicase |
| 129 | ES | Spy1307 | - | 2.0 | RD.2 hypothetical exported protein |
| 130 | ES | Spy1308 | - | 2.0 | RD.2 hypothetical exported protein |
| 131 | ES | Spy1311 | - | 1.6 | RD.2 DNA segregation ATPase related protein |
| 132 | ES | Spy1314 | - | 1.7 | RD.2 hypothetical protein |
| 133 | ES | Spy1322 | - | 1.7 | RD.2 FtsK/SpoIIIE family |
| 134 | ES | Spy1323 | - | 1.8 | RD.2 hypothetical protein |
| 135 | ES | Spy1324 | - | 1.6 | RD.2 hypothetical protein |
| 136 | ES | Spy1325 | - | 1.7 | RD.2 putative cell surface protein |
| 137 | ES | Spy1349 | - | 3.1 | Sugar-binding protein |
| 138 | ES | Spy1350 | - | 3.1 | Sugar transport system permease protein |
| 139 | ES | Spy1351 | - | 2.9 | Sugar transport system permease protein |
| 140 | ES | Spy1352 | *nagC* | 3.3 | Glucokinase or transcriptional regulator |
| 141 | ES | Spy1353 | - | 3.7 | Hypothetical protein |
| 142 | ES | Spy1354 | - | 3.3 | Beta-glucosidase |
| 143 | ES | Spy1355 | *hyl* | 3.5 | Hyaluronoglucosaminidase |
| 144 | ES | Spy1356 | - | 3.2 | Transcriptional regulator, GntR family |
| 145 | ES | Spy1357 | - | 3.8 | Hypothetical protein |
| 146 | ES | Spy1358 | - | 4.1 | Alpha-mannosidase |
| 147 | ES | Spy1368 | *comFA* | 1.9 | ComF operon protein 1 |
| 148 | ES | Spy1438 | *lacD.1* | 2.8 | Tagatose-bisphosphate aldolase |
| 149 | ES | Spy1439 | *lacC.1* | 2.7 | Tagatose-6-phosphate kinase |
| 150 | ES | Spy1440 | *lacB.1* | 2.5 | Galactose-6-phosphate isomerase LacB subunit |
| 151 | ES | Spy1441 | *lacA.1* | 2.5 | Galactose-6-phosphate isomerase LacA subunit |
| 152 | ES | Spy1447 | *copZ* | 3.1 | Copper chaperone |
| 153 | ES | Spy1448 | *copA* | 3.5 | Copper-exporting ATPase |
| 154 | ES | Spy1449 | *copY* | 2.3 | CopAB ATPases metal-fist type repressor |
| 155 | ES | Spy1486 | *dnaJ* | 3.2 | Chaperone protein |
| 156 | ES | Spy1487 | *dnaK* | 3.3 | Chaperone protein |
| 157 | ES | Spy1488 | *grpE* | 3.1 | Hypothetical protein |

| | Growth[1] | Spy | gene | FC[2] | Function |
|---|---|---|---|---|---|
| 158 | ES | Spy1489 | *hrcA* | 2.4 | Heat-inducible transcription repressor |
| 159 | ES | Spy1511 | *cbiO* | 1.5 | Cobalt transport ATP-binding protein CbiO |
| 160 | ES | Spy1512 | *cbiQ* | 1.5 | Cobalt transport protein cbiQ |
| 161 | ES | Spy1514 | - | 1.6 | ABC transporter ATP-binding protein |
| 162 | ES | Spy1515 | - | 1.8 | ABC transporter ATP-binding protein |
| 163 | ES | Spy1516 | *fhuC* | 1.7 | Ferrichrome transport ATP-binding protein |
| 164 | ES | Spy1517 | *fhuB* | 1.6 | Ferrichrome transport system permease protein |
| 165 | ES | Spy1518 | *fhuD* | 1.5 | Ferrichrome-binding protein |
| 166 | ES | Spy1527 | *endoS* | 2.7 | Endo-beta-N-acetylglucosaminidase F2 precursor |
| 167 | ES | Spy1528 | - | 2.3 | Hypothetical protein |
| 168 | ES | Spy1529 | *scrA* | 2.5 | PTS system, sucrose-specific IIABC component |
| 169 | ES | Spy1543 | - | 1.5 | Hypothetical protein |
| 170 | ES | Spy1603 | - | 1.6 | Hypothetical protein |
| 171 | ES | Spy1605 | *thiD* | 1.6 | Hydroxymethylpyrimidine kinase |
| 172 | ES | Spy1606 | - | 1.6 | tRNA pseudouridine synthase A |
| 173 | ES | Spy1619 | *salX* | 1.6 | Lantibiotic transport ATP-binding protein |
| 174 | ES | Spy1620 | *salB* | 1.6 | Serine (threonine) dehydratase |
| 175 | ES | Spy1621 | *salA* | 2.0 | Lantibiotic salivaricin A |
| 176 | ES | Spy1622 | *lacG* | 2.4 | 6-phospho-beta-galactosidase |
| 177 | ES | Spy1623 | *lacE* | 3.1 | PTS system, lactose-specific IIBC component |
| 178 | ES | Spy1624 | *lacF* | 3.0 | PTS system, lactose-specific IIA component |
| 179 | ES | Spy1625 | *lacD.2* | 2.8 | Tagatose-bisphosphate aldolase |
| 180 | ES | Spy1626 | *lacC.2* | 2.5 | Tagatose-6-phosphate kinase |
| 181 | ES | Spy1627 | *lacB.2* | 2.4 | Galactose-6-phosphate isomerase LacB subunit |
| 182 | ES | Spy1628 | *lacA.2* | 2.1 | Galactose-6-phosphate isomerase LacA subunit |
| 183 | ES | Spy1649 | - | 1.9 | Transaldolase |
| 184 | ES | Spy1650 | - | 2.2 | Putative transport protein sgaT |
| 185 | ES | Spy1651 | - | 2.4 | PTS system, IIB component |
| 186 | ES | Spy1652 | - | 1.9 | Transcription antiterminator, BglG family |
| 187 | ES | Spy1667 | - | 2.2 | Hypothetical protein |
| 188 | ES | Spy1668 | *pulA* | 2.1 | Pullulanase |
| 189 | ES | Spy1669 | *dexB* | 2.0 | Glucan 1,6-alpha-glucosidase |
| 190 | ES | Spy1677 | *xthA* | 2.0 | Exodeoxyribonuclease III |
| 191 | ES | Spy1678 | - | 2.4 | PTS system, glucose-specific IIABC component |
| 192 | ES | Spy1679 | - | 1.9 | Hypothetical cytosolic protein |
| 193 | ES | Spy1680 | *prmA* | 1.9 | Ribosomal protein L11 methyltransferase |
| 194 | ES | Spy1681 | - | 1.8 | Hypothetical protein |
| 195 | ES | Spy1683 | *trpG* | 1.7 | *p*-aminobenzoate synthase glutamine amidotransferase component II |
| 196 | ES | Spy1684 | - | 1.8 | ATPase, AAA family |
| 197 | ES | Spy1686 | *flaR* | 1.7 | DNA topology modulation protein flar-related protein |
| 198 | ES | Spy1699 | - | 1.5 | Cell surface protein |
| 199 | ES | Spy1706 | - | 2.8 | Hypothetical protein |
| 200 | ES | Spy1707 | *isp* | 3.1 | Immunogenic secreted protein |

| | Growth[1] | Spy | gene | FC[2] | Function |
|---|---|---|---|---|---|
| 201 | ES | Spy1708 | *ihk* | 2.9 | Two component system histidine kinase |
| 202 | ES | Spy1709 | *irr* | 2.9 | Two-component response regulator |
| 203 | ES | Spy1710 | - | 3.5 | ABC transporter permease protein |
| 204 | ES | Spy1711 | - | 4.8 | ABC transporter ATP-binding protein |
| 205 | ES | Spy1712 | - | 4.6 | Component of efflux system |
| 206 | ES | Spy1713 | - | 3.7 | Hypothetical protein |
| 207 | ES | Spy1718 | - | 2.6 | Protein export protein prsA precursor |
| 208 | ES | Spy1719 | - | 2.7 | Hypothetical protein |
| 209 | ES | Spy1720 | - | 2.6 | Streptopain fragment |
| 210 | ES | Spy1721 | *speB* | 2.7 | Streptococcal pyrogenic exotoxin B |
| 211 | ES | Spy1725 | *mf* | 2.1 | Mitogenic factor |
| 212 | ES | Spy1726 | - | 2.3 | Hypothetical protein |
| 213 | ES | Spy1734 | - | 1.7 | Sorbitol operon regulator |
| 214 | ES | Spy1747 | *groEL* | 2.3 | 60 kDa chaperonin |
| 215 | ES | Spy1759 | - | 2.2 | Hypothetical cytosolic protein |
| 216 | ES | Spy1760 | - | 2.3 | Amino acid permease |
| 217 | ES | Spy1761 | *hutH* | 2.2 | Histidine ammonia-lyase |
| 218 | ES | Spy1773 | *nrdG* | 2.5 | Ribonucleoside-triphosphate reductase activating protein |
| 219 | ES | Spy1774 | - | 2.6 | Acetyltransferase |
| 220 | ES | Spy1775 | - | 2.7 | Putative oxidoreductase |
| 221 | ES | Spy1776 | - | 3.0 | Hypothetical protein |
| 222 | ES | Spy1777 | *nrdD* | 3.6 | Anaerobic ribonucleoside-triphosphate reductase |
| 223 | ES | Spy1805 | - | 1.9 | 6180.3 phage protein |
| 224 | ES | Spy1847 | - | 2.3 | 6180.4 phage protein |
| 225 | ES | Spy1873 | *sdhB* | 3.4 | L-serine dehydratase |
| 226 | ES | Spy1874 | *sdhA* | 3.7 | L-serine dehydratase |

### B. Downregulated genes.

| | Growth | Spy | gene | FC | Function |
|---|---|---|---|---|---|
| 1 | ME | Spy0339 | - | -2.3 | Transposase |
| 2 | ME | Spy0540 | *sagA* | -2.7 | Streptolysin S precursor |
| 3 | ME | Spy0541 | *sagB* | -2.1 | Streptolysin S biosynthesis protein |
| 4 | ME | Spy0542 | *sagC* | -1.7 | Streptolysin S biosynthesis protein |
| 5 | ME | Spy0543 | *sagD* | -1.7 | Streptolysin S biosynthesis protein |
| 6 | ME | Spy0544 | *sagE* | -1.7 | Streptolysin S putative self-immunity protein |
| 7 | ME | Spy0545 | *sagF* | -1.7 | Streptolysin S biosynthesis protein |
| 8 | ME | Spy0546 | *sagG* | -1.6 | Streptolysin S export ATP-binding protein |
| 9 | ME | Spy0547 | *sagH* | -1.6 | Streptolysin S export transmembrane protein |
| 10 | ME | Spy0548 | *sagI* | -1.6 | Streptolysin S export transmembrane protein |
| 11 | ME | Spy0951 | - | -1.5 | Hypothetical protein |
| 12 | ME | Spy0999 | - | -1.5 | 6180.1 phage protein |

| | Growth | Spy | gene | FC | Function |
|---|---|---|---|---|---|
| 13 | ME | Spy1001 | - | -1.6 | 6180.1 phage protein |
| 14 | ME | Spy1336 | *Spy1336/R28* | -89.2 | RD.2 R28 virulence factor |
| 15 | ME | Spy1337 | *Spy1337* | -664.8 | RD.2 Spy1337 transcriptional regulator, AraC family |
| 16 | ME | Spy1338 | - | -1.6 | Phospho-2-dehydro-3-deoxyheptonate aldolase |
| 17 | ME | Spy1339 | *aroB* | -1.5 | 3-dehydroquinate synthase |
| 18 | ME | Spy1340 | - | -1.8 | Hypothetical protein |
| 19 | ME | Spy1482 | *acpP.2* | -1.6 | Acyl carrier protein |
| 20 | ME | Spy1483 | *fabH* | -2.0 | 3-oxoacyl-[acyl-carrier-protein] synthase III |
| 21 | ME | Spy1484 | *fabT* | -1.6 | Transcriptional regulator, MarR family |
| 22 | ME | Spy1653 | - | -1.5 | Hypothetical protein |
| 23 | ME | Spy1740 | - | -2.1 | Translation initiation inhibitor |
| 24 | ME | Spy1799 | - | -1.5 | 6180.3 phage protein |
| 1 | ES | Spy0014 | - | -2.3 | Amino acid permease |
| 2 | ES | Spy0063 | *secY* | -1.7 | Protein translocase subunit |
| 3 | ES | Spy0064 | *adk* | -2.2 | Adenylate kinase |
| 4 | ES | Spy0065 | *infA* | -1.7 | Translation initiation factor IF-1 |
| 5 | ES | Spy0066 | *rpmJ* | -1.7 | LSU ribosomal protein L36P |
| 6 | ES | Spy0067 | *rpsM* | -1.7 | SSU ribosomal protein S13P |
| 7 | ES | Spy0068 | *rpsK* | -1.7 | SSU ribosomal protein S11 |
| 8 | ES | Spy0069 | *rpoA* | -1.7 | RNA polymerase alpha chain |
| 9 | ES | Spy0070 | *rplQ* | -1.7 | LSU ribosomal protein L17P |
| 10 | ES | Spy0079 | *tyrS* | -2.6 | Tyrosyl-tRNA synthetase |
| 11 | ES | Spy0092 | *ackA* | -1.6 | Acetate kinase |
| 12 | ES | Spy0100 | *ssb* | -2.3 | Single-strand DNA binding protein |
| 13 | ES | Spy0144 | *metB* | -1.7 | Cystathionine beta-lyase |
| 14 | ES | Spy0153 | - | -1.5 | Transcription antiterminator, BglG family |
| 15 | ES | Spy0261 | - | -1.9 | Hypothetical cytosolic protein |
| 16 | ES | Spy0262 | - | -1.5 | ABC transporter substrate-binding protein |
| 17 | ES | Spy0270 | *gidB* | -1.7 | Glucose inhibited division protein B |
| 18 | ES | Spy0271 | *lemA* | -1.6 | Hypothetical protein |
| 19 | ES | Spy0272 | - | -1.5 | Heat shock protein HtpX |
| 20 | ES | Spy0359 | *mtsC* | -1.6 | Manganese transport system membrane protein |
| 21 | ES | Spy0364 | *rplA* | -2.0 | LSU ribosomal protein L1P |
| 22 | ES | Spy0405 | *pcp* | -1.9 | Pyrrolidone-carboxylate peptidase |
| 23 | ES | Spy0427 | *smc* | -1.6 | Chromosome partition protein smc |
| 24 | ES | Spy0444 | - | -1.6 | RelB-domain protein |
| 25 | ES | Spy0445 | - | -1.7 | ParE-domain protein |
| 26 | ES | Spy0446 | - | -1.7 | Hypothetical cytosolic protein |
| 27 | ES | Spy0447 | - | -1.8 | Hypothetical protein |
| 28 | ES | Spy0449 | - | -1.9 | Transposase |
| 29 | ES | Spy0481 | - | -1.5 | Hypothetical exported protein |
| 30 | ES | Spy0553 | *atpE* | -1.8 | ATP synthase C chain |
| 31 | ES | Spy0554 | *atpB* | -2.0 | ATP synthase A chain |

| | Growth | Spy | gene | FC | Function |
|---|---|---|---|---|---|
| 32 | ES | Spy0555 | *atpF* | -1.9 | ATP synthase B chain |
| 33 | ES | Spy0556 | *atpH* | -1.9 | ATP synthase delta chain |
| 34 | ES | Spy0557 | *atpA* | -1.7 | ATP synthase alpha chain |
| 35 | ES | Spy0558 | *atpG* | -1.6 | ATP synthase gamma chain |
| 36 | ES | Spy0559 | *atpD* | -1.5 | ATPase, beta subunit |
| 37 | ES | Spy0560 | *atpC* | -1.5 | ATP synthase epsilon chain |
| 38 | ES | Spy0561 | - | -1.5 | Hypothetical membrane associated protein |
| 39 | ES | Spy0598 | *infC* | -1.9 | Bacterial protein translation initiation factor IF-3 |
| 40 | ES | Spy0599 | *rpl36* | -2.0 | LSU ribosomal protein L35P |
| 41 | ES | Spy0600 | *rplT* | -2.1 | LSU ribosomal protein L20P |
| 42 | ES | Spy0630 | - | -1.7 | RNA binding protein |
| 43 | ES | Spy0631 | - | -1.9 | Hypothetical protein |
| 44 | ES | Spy0666 | - | -1.9 | 3-hydroxy-3-methylglutaryl-coenzyme A reductase |
| 45 | ES | Spy0667 | *mvaS.1* | -2.0 | Hydroxymethylglutaryl-CoA synthase |
| 46 | ES | Spy0673 | - | -1.5 | Hypothetical protein |
| 47 | ES | Spy0691 | *parE* | -1.5 | Topoisomerase IV subunit B |
| 48 | ES | Spy0692 | *parC* | -1.6 | Topoisomerase IV subunit A |
| 49 | ES | Spy0693 | *bcaT* | -2.2 | Branched-chain amino acid aminotransferase |
| 50 | ES | Spy0694 | - | -2.0 | Hypothetical cytosolic protein |
| 51 | ES | Spy0824 | - | -1.8 | ATP-NAD kinase |
| 52 | ES | Spy0910 | *pgmA* | -1.6 | Phosphomannomutase |
| 53 | ES | Spy0918 | *rpsT* | -1.9 | SSU ribosomal protein S20P |
| 54 | ES | Spy1076 | - | -1.5 | Hypothetical cytosolic protein |
| 55 | ES | Spy1087 | - | -1.7 | RD.1 hypothetical cytosolic protein |
| 56 | ES | Spy1088 | - | -1.7 | RD.1 hypothetical protein |
| 57 | ES | Spy1089 | - | -1.6 | RD.1 hypothetical cytosolic protein |
| 58 | ES | Spy1090 | - | -1.6 | tRNA (uracil-5-)-methyltransferase |
| 59 | ES | Spy1098 | *grab* | -1.9 | Protein G-related alpha 2M-binding protein |
| 60 | ES | Spy1193 | - | -1.7 | Hypothetical membrane spanning protein |
| 61 | ES | Spy1336 | *Spy1336/R28* | -355.4 | RD.2 R28 virulence factor |
| 62 | ES | Spy1337 | *Spy1337* | -478.9 | RD.2 Spy1337 transcriptional regulator, AraC family |
| 63 | ES | Spy1338 | - | -2.5 | Phospho-2-dehydro-3-deoxyheptonate aldolase |
| 64 | ES | Spy1339 | *aroB* | -2.5 | 3-dehydroquinate synthase |
| 65 | ES | Spy1340 | - | -2.5 | Hypothetical protein |
| 66 | ES | Spy1371 | - | -1.7 | s1-type RNA-binding domain |
| 67 | ES | Spy1372 | - | -1.6 | Peptidyl-prolyl cis-trans isomerase |
| 68 | ES | Spy1401 | - | -1.6 | Thioredoxin reductase |
| 69 | ES | Spy1403 | - | -1.9 | Transporter |
| 70 | ES | Spy1404 | - | -1.7 | Amino acid ABC transporter permease protein |
| 71 | ES | Spy1434 | - | -1.8 | DegV family protein |
| 72 | ES | Spy1463 | *lytR* | -1.5 | Transcriptional regulator, LytR family |
| 73 | ES | Spy1468 | *manL* | -1.8 | PTS system, mannose-specific IIAB component |
| 74 | ES | Spy1469 | *manM* | -1.5 | PTS system, mannose-specific IIC component |

| | Growth | Spy | gene | FC | Function |
|---|---|---|---|---|---|
| 75 | ES | Spy1482 | *acpP.2* | -2.4 | Acyl carrier protein |
| 76 | ES | Spy1504 | - | -1.8 | Hydrolase (HAD superfamily) |
| 77 | ES | Spy1591 | *pgk* | -1.6 | Phosphoglycerate kinase |
| 78 | ES | Spy1643 | *cysS* | -1.5 | Cysteinyl-tRNA synthetase |
| 79 | ES | Spy1654 | - | -2.0 | SSU ribosomal protein S15P |
| 80 | ES | Spy1676 | - | -1.6 | NrdI.1 |
| 81 | ES | Spy1825 | *rpmG* | -2.0 | LSU ribosomal protein L33P |
| 82 | ES | Spy1865 | *rpsD* | -2.0 | SSU ribosomal protein S4P |

[(1)] Growth refers to the growth phase at which cells were collected

[(2)] FC, fold change. The FC threshold is 1.5

[(3)] ME, mid-exponential

[(4)] ES, early stationary

### Table S5. Oligonucleotides used for isogenic mutant generation and and PCR amplification of *Spy1336/R28*

| Oligonucleotide | DNA sequence |
|---|---|
| **JE410** | 5'-CGATCTCGTGCTCGATAAGCTGTTTCG-3' |
| **JE412** | 5'-CGCATTTACACTTTCTGGATCACTAGCC-3 |
| **JE431** | 5'-GTTTGTTGGAGAGAGATACATAAC-3'' |
| **JE432** | 5'-GACGAAAATGAAGTTGAACTAGG-3' |
| **JE433** | 5'-GCAGAACCTAAGACAGTTACCATC-3' |
| **HPN-seq** | 5'-ACTAAGACCAATTAGTACAGAAGCT-3' |
| **1336-1** | 5'- GTCCGGATCCTCACCTTCAAGTGAATCATTAGAAC -3' |
| **1336-2** | 5'- TTGAAGAGTCAATAGTTGCTGCATATCTCTCTCCAACAAACTTAATTA -3' |
| **1336-3** | 5'- TAATTAAGTTTGTTGGAGAGAGATATGCAGCAACTATTGACTCTTCAA -3' |
| **1336-4** | 5'- GTCCGGATCCTTTAGCTATTTCTTCTGTTTTAATA -3' |
| **1337-1** | 5'- GTCCGGATCCTAGGTTCTTTAGTAAGATACTTAA -3' |
| **1337-2** | 5'- AGCAAAATGAATATTTTTCGATTACTTTGATTTAATATGTTATCTATT -3' |
| **1337-3** | 5'- AATAGATAACATATTAAATCAAAGTAATCGAAAAATATTCATTTTGCT -3' |
| **1337-4** | 5'- GTCCGGATCCATGGACATGAAGTGTTTGTGGCAT -3' |
| **1336-1337-1** | 5'- GTCCGGATCCTCACCTTCAAGTGAATCATTAGAAC -3' |
| **1336-1337-2** | 5'- TGAAGGGAATATTAAGCAAAATGAGCCTATCGAGATTATTAATTTCGA -3' |
| **1336-1337-3** | 5'- TCGAAATTAATAATCTCGATAGGCTCATTTTGCTTAATATTCCCTTCA -3' |
| **1336-1337-4** | 5'- GTCCGGATCCTCATCAGTACTAGGTAACAGAGATA -3' |

### Table S6. Isogenic mutants generated in this study

| Isogenic strains | Number of Ts in $HT_{Spy1336-7}$ | *Spy1336* present | *Spy1337* present |
|---:|---|---|---|
| **MGAS27961-9T** | 9 | √ | √ |
| **MGAS27961-10T** | 10 | √ | √ |
| **MGAS27961-11T** | 11 | √ | √ |
| **MGAS27961-10T-Δ*Spy1336*** | 10 | - | √ |
| **MGAS27961-10T-Δ*Spy1337*** | 10 | √ | - |
| **MGAS27961-10T-Δ*Spy1336*/Δ*Spy1336*** | 10 | - | - |
